# *In silico* analysis and Predictive linkage of Deubiquitinating Enzymes underlying Early Development

**DOI:** 10.1101/2025.05.03.652026

**Authors:** Fahima Munavar-K, Nibedita Lenka

## Abstract

Deubiquitinating enzymes (DUBs) exert their functions by catalyzing the ubiquitinated proteins and maintaining ubiquitination dynamics during post-translational modification. They carry out wide-gamut of functions in various cellular contexts by being associated with regulatory entities concerning transcription, translation, cell signalling, etc. However, limited studies are available concerning their specific attributes during organismal development. In this study, we have employed the integrated bioinformatics and experimental approaches to investigate the involvement of DUBs in embryonic stem cell (ESC) maintenance and differentiation. The StemMapper database revealed the distinctive expression profiles of various DUBs in ESCs and their differentiated derivatives. Further, experimental validation by qRT-PCR with a selected group of understudied DUBs including USP46, USP47, USP4, USP40, CYLD, and BRCC3 confirmed their expression patterns during cardiac and neural differentiation from ESCs, with USP46, USP47, and USP40 showing an increasing trend in contrast to USP4 during differentiation. While the TRANS-DSI database revealed novel DUB-Substrate interactions (DSI), the Gene Ontology and pathway enrichment analysis helped link these DUBs to critical cellular processes including transcriptional regulation, cell cycle control, and DNA repair. Further, the UbiBrowser 2.0 facilitated identifying several understudied DUBs as potential modulators of evolutionarily conserved developmental signalling cascades, notably Wnt, Notch, Hippo, and Hedgehog pathways. Our *in silico* analysis and predictions do provide crucial insights into the complex regulatory roles of DUBs in early development and establish a foundation for further investigations to unveil the molecular mechanisms and identify potential therapeutics thereof.

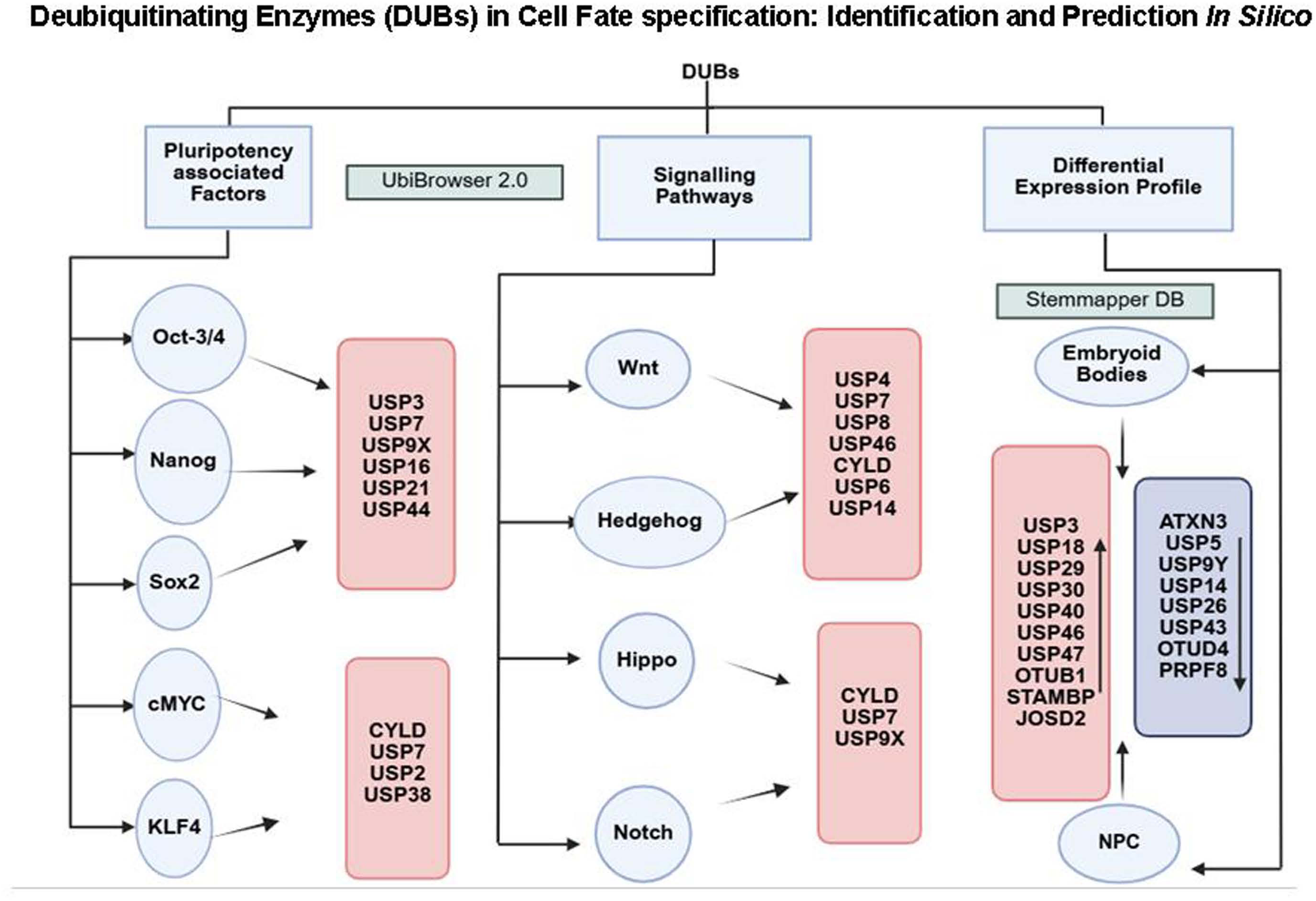

## Introduction

Ubiquitination and deubiquitination are post-translational modifications that are crucial in different cellular processes. The ubiquitination process is carried out by three classes of enzymes, E1 (activating enzyme), E2 (conjugating enzyme) and E3 (ubiquitin ligase). The ubiquitin is activated by E1 in an ATP-dependent manner and then transferred to E2, and finally, E3 adds ubiquitin on a target protein which can be recognized by proteasome for degradation [1]. There are about 600-700 E3 ligases that account for diverse substrate specificity [2]. Based on the number (mono or poly), and site of modification (K9, K11, K48, K63), ubiquitination is involved in different processes like targeting the protein for degradation, membrane targeting and signalling cascades [3]. These modifications regulate protein stability, function, and cellular interactions and thereby impact different cellular functions.

Deubiquitinating enzymes (DUBs) are a group of proteases that catalyze the deconjugation of ubiquitin moiety from proteins, thereby antagonizing the function of ubiquitin E3 ligases. There are about 100 DUBs that have been identified so far and categorized into seven families based on the catalytic domain architecture. Based on the cellular context, a DUB can be attributed to multiple functions. In fact, the functions of the DUBs extend to the regulation of turnover rate, activation, recycling and localization of different proteins [4, 5]. By removing the K48-linked poly-ubiquitin, DUB can rescue the protein tagged for proteasomal degradation thereby increasing the half-life and conferring stability of the substrate protein [4, 6]. Contrastingly, DUBs can modulate a myriad of cellular processes like protein-protein interaction, signalling, endocytosis and protein trafficking by altering K63-linked ubiquitin moieties [4, 7]. Additionally, DUBs altering the mono-ubiquitination status of histone proteins have epigenetic modulatory roles and are involved in DNA damage response, cell cycle control, transcriptional regulation, etc. [8–10]. Of late, DUBs have drawn immense attention concerning their link with various signalling modulators and also in gaining insights into their plausible involvement in various cellular processes during development and tumorigenesis. In fact, a very few DUBs have been well characterized in the context of both disease and development across species. For instance, USP9x is essential for both embryonic and adult neurogenesis modulating the neural progenitor generation and polarity in various species spanning zebra fish to mouse and human [11–13]. The same is also reported to render crucial influence in EMT by regulation of survivin in pancreatic cancer [14]. Similarly, 57 DUBs have been found to be essential for zebra fish embryonic development, the lack of which results in notochord deformities, neuronal defects, etc [15]. In *C. elegans*, 3 DUBs - MATH-33, USP-46 and USP-47 are reported to be critical for centrosome positioning and PAR protein asymmetry in one-cell embryos [16]. USP7 is another important DUB that has a dual role in early development and cancer. Knockdown of USP7 negatively affects genome stability by causing DNA damage and leads to developmental defects and apoptosis [17]. However, it is important to note its contrasting function in cancer progression through its association with p53, NOTCH1 and β-catenin [18].

Since the Ubiquitin-proteasome system (UPS) targets regulatory proteins that play important roles in cell cycle progression, DNA damage response, tumorigenesis as well as early development and differentiation, it is recognized as a crucial mediator/modulator governing organismal development and disease. Dysregulation of these ubiquitination-mediated processes can lead to developmental abnormalities and contribute to tumorigenesis, tumor progression, and therapy resistance. Moreover, their context-dependent functions highlight the importance of ubiquitination-deubiquitination homeostasis underlying various physiological processes and any anomaly in the same dictates disease onset including malignancy. In fact, there is a lot of interest in targeting the E3 ligases in cancer therapeutics [19]. Precise UPS regulation is also required for ESCs to balance self-renewal and differentiation. The core pluripotency factors such as OCT4, NANOG, Sox2 and c-MYC are tightly controlled by ubiquitin-mediated degradation [20]. Moreover, the signalling pathways crucial for normal embryonic development and various cellular functions such as Wnt, Notch, Hippo, and Hedgehog are often dysregulated in disease states including cancer. This underscores the need to explore the key players and their attributes in maintaining cellular homeostasis in addition to determining the causality and interconnectivity manifested among those during normal development and disease. Despite their significance, DUBs remain understudied in developmental context compared to their roles in disease and this study attempts to bridge the gap.

Using different bioinformatics tools, we have conducted the *in silico* predictions to investigate diverse roles of DUBs along with their association with components of the key signalling pathways underlying early development and cancer. The differential expression patterns of the selected candidate DUBs across different stem cells and cancer also suggested their complex interplay and plausible association with development and disease. Hence, understanding the UPS-DUB attributes in stem cells and the mechanistic underpinning would provide insights into the fundamental biological processes that would further assign potential implications in regenerative medicine and cancer biology.

## Materials and Methods

### UbiBrowser

For the prediction of DUBs involved in different signalling pathways such as Wnt, Notch, Hippo, and Hedgehog, we used UbiBrowser 2.0 (http://ubibrowser.ncpsb.org.cn/), as described [21]. It is a comprehensive database comprising information on the Ubiquitin–Proteasome system. This computational platform integrates structural, evolutionary, and functional data from multiple sources including manual curation, GO annotation, protein domain, motif and network topology. Our query was based on potential DUBs reported or predicted to act on transcription activators and ligands of the stated signaling pathways. We chose β-catenin (CTNNB1, AXIN 1 as components of Wnt pathway and NOTCH1 receptor for the Notch pathway, Smoothened (SMO) for Hedgehog pathway and YAP1, LATS1 for Hippo pathway to identify potential DUB interactions with each of these components. The confidence score threshold was set at ≥ 0.5 to ensure high-confidence predictions. The database provided a list of predicted DUBs for each queried protein along with the confidence score and supporting evidences. The candidates were further shortlisted based on their associations with multiple signalling pathways.

### StemMapper

StemMapper database (https://stemmapper.sysbiolab.eu/), developed by Pinto et al. [22], includes the integrated datasets on various lineage-specific gene expression spanning both murine and human stem cells. We took advantage of the same to have a comparative assessment of DUBs expressed in pluripotent ESCs during both maintenance and differentiation. While the initial comparison was conducted using datasets from murine ESCs and ESC-derived embryoid bodies (EBs) or neural progenitor cells (NPCs), further comparison was done considering both murine and human datasets and narrowing down to a few of the selected candidate DUBs based on their differential expression pattern, *in silico* predictions and existing literature review. We compared the expression status of the selected DUBs in ESCs with that of cardiomyocytes and neurons as the mesoderm and ectoderm derivatives respectively. Similarly, for endodermal derivatives we used the datasets from hepatocytes (*Mus musculus*) or pancreatic beta cells (*Homo sapiens*) for comparison. Following projects were used to retrieve the data for normal/wild type cells, (i) ESC vs EBs - GSE8128, GSE119085, GSE3653, (ii) ESC vs NPCs - GSE44175, (iii) ESC vs Cardiomyocyte (Ms) - GSE6689, GSE5671, (iv) ESC vs Neurons (Ms) - GSE44175, (v) ESC vs Hepatocytes (Ms) GSE44923, GSE7038, GSE3653 (vi) ESC vs Cardiomyocyte (Hu) - GSE54186, GSE79413, (vii) ESC vs Neurons (Hu) - GSE41439, GSE75701 (viii) ESC vs Beta cells (Hu) - GSE41439, GSE40709. A threshold of log2 fold change > 1.5 and a p-value threshold of 0.05 to indicate statistically significant difference was considered to narrow down the differentially expressed DUBs and the heatmaps were generated using SR plots tool [23]. We further validated the expression profile of the selected DUBs at different stages of cardiomyogenic and neural differentiation from murine ESCs by conducting qRT-PCR.

### TCGA

The Cancer Genome Atlas (TCGA) is a widely-adopted comprehensive cancer genome database catering to the biomedical researchers world-wide. We utilized the UALCAN portal (http://ualcan.path.uab.edu/) to analyze the expression profiles of selected DUBs across multiple cancer types by comparing the datasets available in the TCGA database [24] for the primary tumor samples and normal tissues. Expression values were reported as transcripts per million (TPM). The expression data were visualized using UALCAN’s built-in plotting tools and the box plots were generated to reflect the expression distribution pattern.

### Trans DSI

Trans-DSI is a computational framework designed to predict protein-protein interactions and it relies on machine learning models trained on broad interaction databases (e.g., BioGRID, STRING) to predict novel protein-protein interactions [25]. Hence, we utilized the information from the Trans-DSI database to predict the potential DUB-substrate interaction (DSI) and curate a list of potential substrates for the candidate DUBs. A confidence score threshold was set at ≥ 0.5 in order to obtain higher confidence predictions. For each DUB we collected information on the predicted substrates along with associated confidence scores for further analysis (Table S1) and some of those were matched with existing literature and databases.

### Gene Ontology (GO) enrichment analysis

We performed Gene Ontology (GO) enrichment analysis of the predicted substrates mentioned in Table S1 to understand the functional relevance of the candidate DUBs with the help of Metascape tool (https://metascape.org), a web-based portal for gene annotation and analysis [26]. The parameters considered were GO Biological Processes and GO Molecular Functions that were grouped into clusters based on the similarity and with a minimum count of 3; p-value ≤ 0.01; and an enrichment factor > 1.5. The top 20 clusters were plotted using SR Plots to provide an overview of the enriched functional terms associated with the predicted DUB substrates. The differentially expressed DUBs were analyzed in the context of their known functions and their association with signaling pathways. This also helped us to predict the potential functional roles of the DUB substrates in their biological roles during cellular differentiation.

### ESC maintenance and differentiation

We have used mouse ESC line D3 (ATCC # CRL-1934; RRID: CVCL_4378) for the maintenance and differentiation studies. ESCs were maintained in DMEM supplemented with 15% FBS, L-glutamine, penicillin-streptomycin, nonessential amino acids (all from Invitrogen), β-mercaptoethanol (Sigma-Aldrich) and Leukemia inhibitory factor (LIF) (1000U/mL; Merck-Millipore) in a humidified incubator at 37 °C with 5% CO2, as described [27]. While spontaneous differentiation into three germ layers was achieved by the withdrawal of LIF, the directed differentiation towards cardiac lineage was carried out by following the protocol described [28]. Embryoid bodies (EBs) were generated in 20μL hanging drops with 600 cells/drop for 2 days (d2 EB) followed by suspension culture for 3 days (d5 EB). The cardiac differentiation was induced by plating the d5 EBs using the maintenance medium without LIF and monitoring the spontaneous pulsating activities in them. Similarly, the neural differentiation was carried out using the monoculture strategy reported earlier [27]. The cells were seeded with DMEM containing 10% FBS without LIF for 2 days followed by 10% Knockout-DMEM (KO) with Knockout serum replacement from d2 onwards. We selected different time points for neural [d2ND, d8-10 (specified as d10ND) and d14ND] and cardiac differentiation [d2CD (d2 EB), d5CD (d5 EB) and d10CD] that corresponded to lineage/differentiation induction-, progenitor- and differentiated stages respectively. The validation of the respective differentiation was carried out by conducting immunocytochemical analysis using the protocol, as described [27, 28]. The antibodies used were OCT4 (Santa Cruz Biotechnology; sc-5279, RRID: AB_628051) for attributing ESCs characteristic during maintenance, Sox1 (Merck; S8318, RRID: AB_1841173) for detecting early neural cells (on d2), Nestin (Merck; MAB353, RRID:AB_94911) and Sox2 (Chemicon; ab5770) for neural progenitors (on d10), Map2 (Merck; M4403, RRID: AB_477193) and GFAP (Merck; G3893, RRID: AB_477010) for detecting differentiated neurons and astrocytes (on d14) respectively, Brachyury (Bry) (Santa Cruz Biotechnology; sc-20109, RRID: AB_2255702) for detecting the pan-mesoderm cells during early stage (on d2), PDGFRα (BD Biosciences; 558774, RRID:AB_397117) for cardiomyogennic progenitors (on d5) and cardiac troponin-T (cTnT: BD Biosciences; 564766, RRID:AB_2738938) for cardiomyocytes (on d10) during ESCs differentiation.

### RNA isolation and qRT-PCR

Total RNA was extracted using Trizol reagent (Invitrogen) followed by cDNA synthesis and DNAse-I treatment, as described [28]. qRT-PCR was carried out with SSOFast Eva green supermix (Biorad) and using either CFX-96 (BioRad) or QuantStudio-5 (Applied Biosystems) thermal cycler. Relative expression level of each gene was normalized with β-actin and calculated using the formula 2^-ΔCT^. The primer sequences are presented in Table S2.

## Results

### Prediction of novel DUBs associated with ESC cell fate specification and pluripotency

To systematically identify novel DUBs that are likely involved in stem cell maintenance and differentiation, we performed an integrated *in-silico* analysis that included the following three criteria; (i) the DUBs that are linked with the core pluripotency-associated factors, (ii) the ones that differentially express during stem cell maintenance and differentiation and (iii) the DUBs that are associated with multiple signalling pathways. To identify the DUBs contributing to ESC pluripotency, we used UbiBrowser 2.0 and looked for the ones linked with the core pluripotency-associated factors such as OCT4, SOX2, NANOG, KLF4 and cMYC. As seen in Fig. 1, USP3 was identified as one of the known DUBs for POU5F1 (OCT4) and NANOG as its targets. However, it appeared under the predicted category for SOX2 and cMYC (Fig. 1a-d). Similarly, USP7 was common between POU5F1 and cMYC among the known DUBs, but a predicted one for SOX2, NANOG and KLF4 (Fig. 1a-e). USP9x on the contrary came under the predicted category for all the factors except SOX2, where it emerged as a known DUB. USP21 was also identified as a known DUB for NANOG in contrast to the same appearing as a predicted one for POU5F1, SOX2 and c-MYC. CYLD however, came as a predicted DUB for POU5F1, cMYC and KLF4. Collectively, our analysis revealed a wide gamut of interacting DUBs associated with factors rendering ESC pluripotency. Further, we examined their expression status along ESC maintenance and differentiation states using the available information from several datasets and the Stemmapper database. Accordingly, a comparative analysis was carried out following the compilation of the datasets available for mouse ESCs and differentiated cells - the embryoid bodies (EB) resembling gastrulating embryos, and also for the neural progenitor cells (NPC). Our prediction was based on the rationale that if a DUB’s expression in undifferentiated ESCs is distinct from that of the differentiated states, there is a greater likelihood that those are associated with the cell fate decision machinery and regulate the same. Among the 6- and 9 datasets considered for ESCs and EBs respectively, a lot of disparity concerning the expression of various DUB candidates was noticed (Fig. 2a) unlike the consistency seen among the 3 datasets each comparing ESCs with NPCs (Fig. 2b). Interestingly, most of the upregulated DUBs were common in EBs and NPCs in different data sets when compared with the ESC datasets (Fig, 2a, b; red*). To name a few, *Usp3, Usp18, Usp29, Usp30, Usp40, Usp46, Usp47, Otub1, Stambp,* and *Josd2* were seen highly expressed in both EBs and NPCs hinting at their association during differentiation. However, majority of the downregulated DUBs in EBs and NPCs showed inconsistency in their expression pattern as discerned among various datasets (Fig. 2a, b; blue*). Both *Usp25* and *Cyld* showed contrasting pattern concerning their expression status in EBs and NPCs compared to their ESC counterparts. While the expression of *Usp25* was higher in EBs, the reverse was true in NPCs (magenta*). Similarly, *Cyld* displayed higher expression in NPCs in contrast to the reduced expression in EBs when compared with their respective ESC datasets. The DUBs such as *Atxn3, Usp5, Usp9y, Usp14, Usp26, USP43, Usp50, Otud4 and Prpf8* were common among the downregulated ones in NPC and EB datasets (Fig. 2a, b; dark grey*). Together our analysis predicted the potential involvement of various DUBs in stem cell differentiation either through a shared or independent mode of regulation of lineage specification. Further validation would be required to discern their specific developmental attributes.

**Fig. 1:**
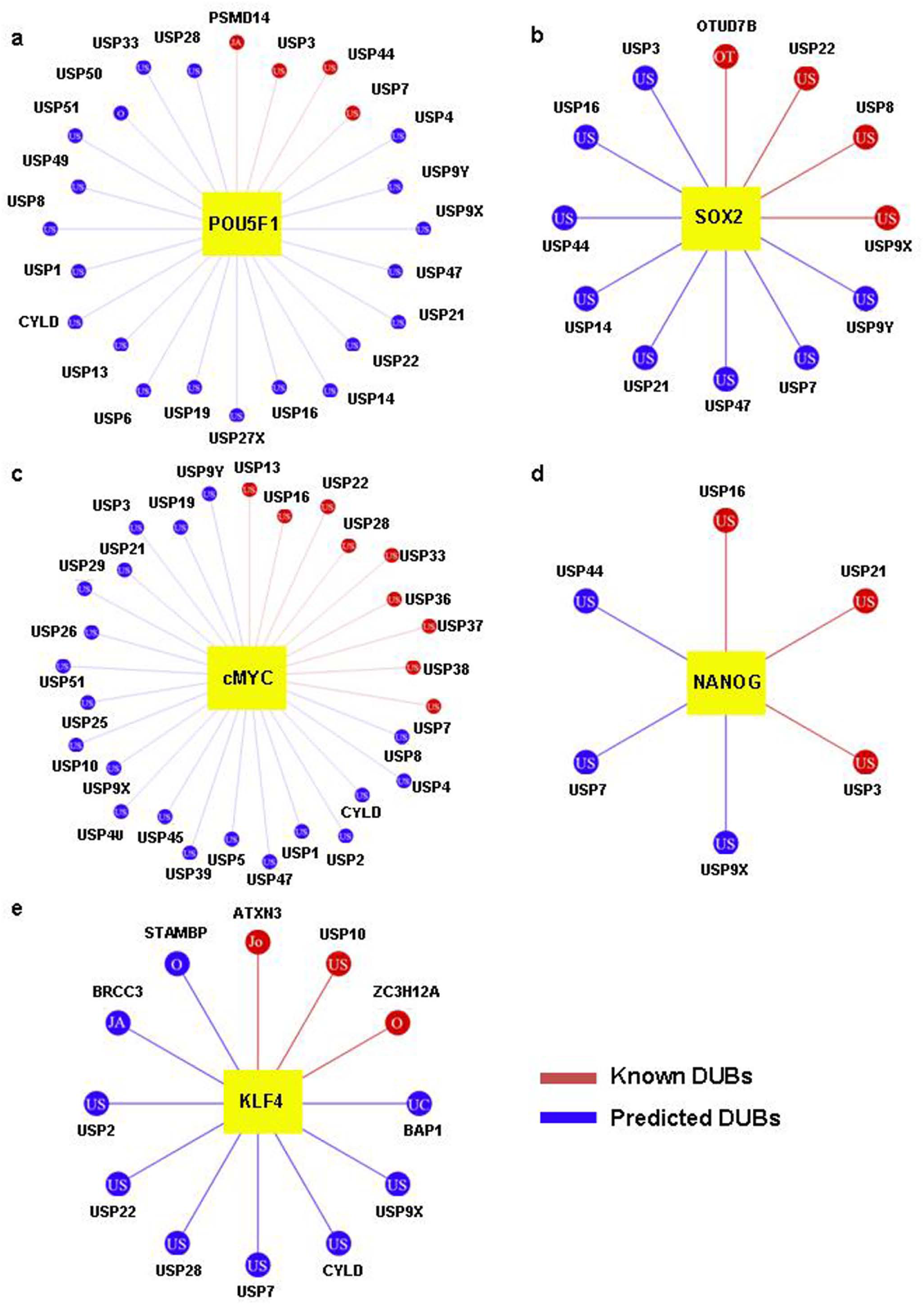
Predicted and known DUBs associated with core pluripotency-associated factors in *Homo sapiens*. Depiction of candidate DUBs associated with pluripotency associated factors (a: OCT4, b: SOX2, c: c-MYC, d: NANOG, and e: KLF4) by using Ubibrowser 2.0. The confidence score threshold was set at ≥ 0.5 to ensure high confidence predictions. The known interactions are indicated in Red and the predicted ones in blue.

**Fig. 2:**
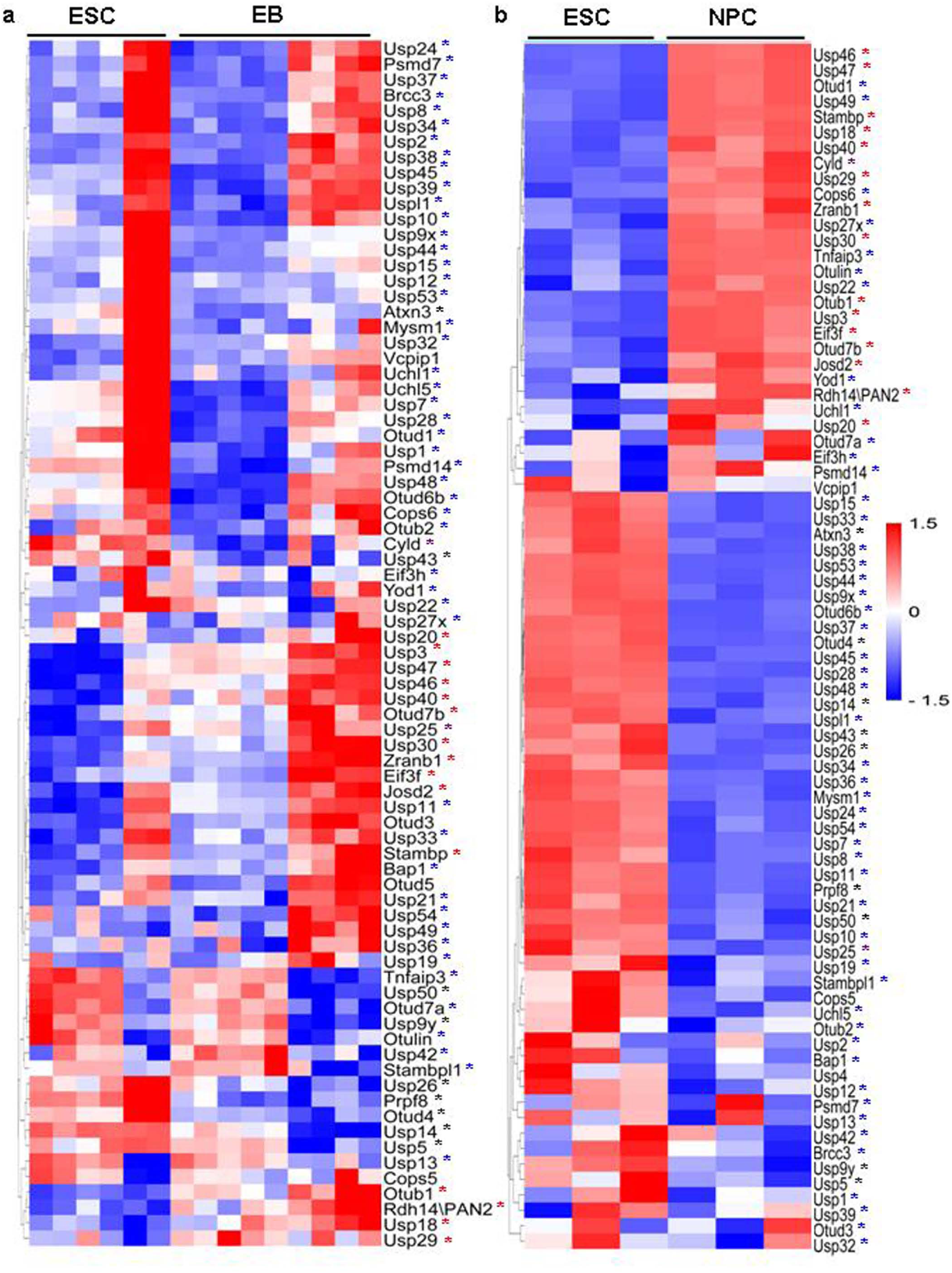
Differential Expression of DUBs in ESCs vs EBs and -NPCs. a, b) Heatmap depicting the differential expression of various DUBs compiled from murine datasets comparing ESC with embryoid body (EB) (a) and ESC with neural progenitor cells (NPCs) (b). The StemMapper database was used to curate the data and represent the differential expression at log 2 scale. Dark grey**\*** indicates the downregulated DUBs in EB and NPC compared to ESC in each; Red**\*,** the upregulated DUBs in EB and NPC compared to ESC in each; Blue*, the inconsistent expression pattern of DUBs in EB and NPC compared to ESC in each; Magenta**\*** denotes the opposite expression pattern of DUBs in ESC-EB group comparing that with ESC-NPC datasets.

### Differential expression of DUBs during lineage specification from ESCs

Based on our integrated approach, we narrowed down to a couple of poorly characterized DUBs (*Usp4, Usp40, Usp46, Usp47, Usp48, Cyld,* and *Brcc3)* along with the two well-studied ones (USP21 and USP3) for further elucidation of their plausible association with differential cell fates from ESCs. Accordingly, we analyzed the status of the afore-stated selected DUBs using the Stemmapper platform. A comparative analysis was performed to assess their expression profiles in ESCs during maintenance and upon subsequent differentiation into all three germ layer derivatives: cardiomyocytes (mesoderm), neurons (ectoderm), and hepatocytes or pancreatic beta cells (endoderm). The data sets from both *Mus musculus* and *Homo sapiens* were considered to assess evolutionary conservation, if any. Our analysis revealed that the expression of USP48 was higher in ESCs in both mouse and human envisioning an evolutionarily conserved role during stem cell maintenance (Fig. 3a-f). Similarly, the conserved expression pattern of USP47 between mouse and human datasets highlighted its plausible functional importance during stem cell differentiation across the species analyzed (Fig. 3a-f). While Usp46 expression was comparatively less in endodermal derivatives in human, the same remained higher in ecto-, endo- and mesodermal derivatives in mouse. In contrast, Brcc3 expression remained higher in mouse ESCs compared to cardiomyocytes and neurons, while the same was seen expressing at a higher level in endodermal derivative, the hepatocytes (Fig. 3a-c). Further, USP4 in both the species, when compared with ESCs in each, displayed relatively higher expression among meso- and endodermal derivatives in contrast to that of neurons, where it remained inconsistent among the curated mouse datasets and lower as per the human datasets (Fig. 3d-f). However, USP40 displayed a context dependent expression in both the species. While its expression was higher in ectodermal (neurons) and mesodermal (cardiomyocyte) derivatives in both the species compared to their ESC counterparts (Fig. 3b, d), it displayed a contrasting expression pattern in endodermal derivatives between the two species analyzed (Fig. 3c, f). Our analysis also corroborated with the established role of USP21 in ESC maintenance and USP3 in ESC differentiation to mesoderm lineage [29, 30]. Together our analysis not only depicted the plausible involvement of these DUBs in cell fate modulation but also provided a basis for further exploration into the specific mechanisms via which these would orchestrate their regulatory role in stem cell fate decision.

**Fig. 3:**
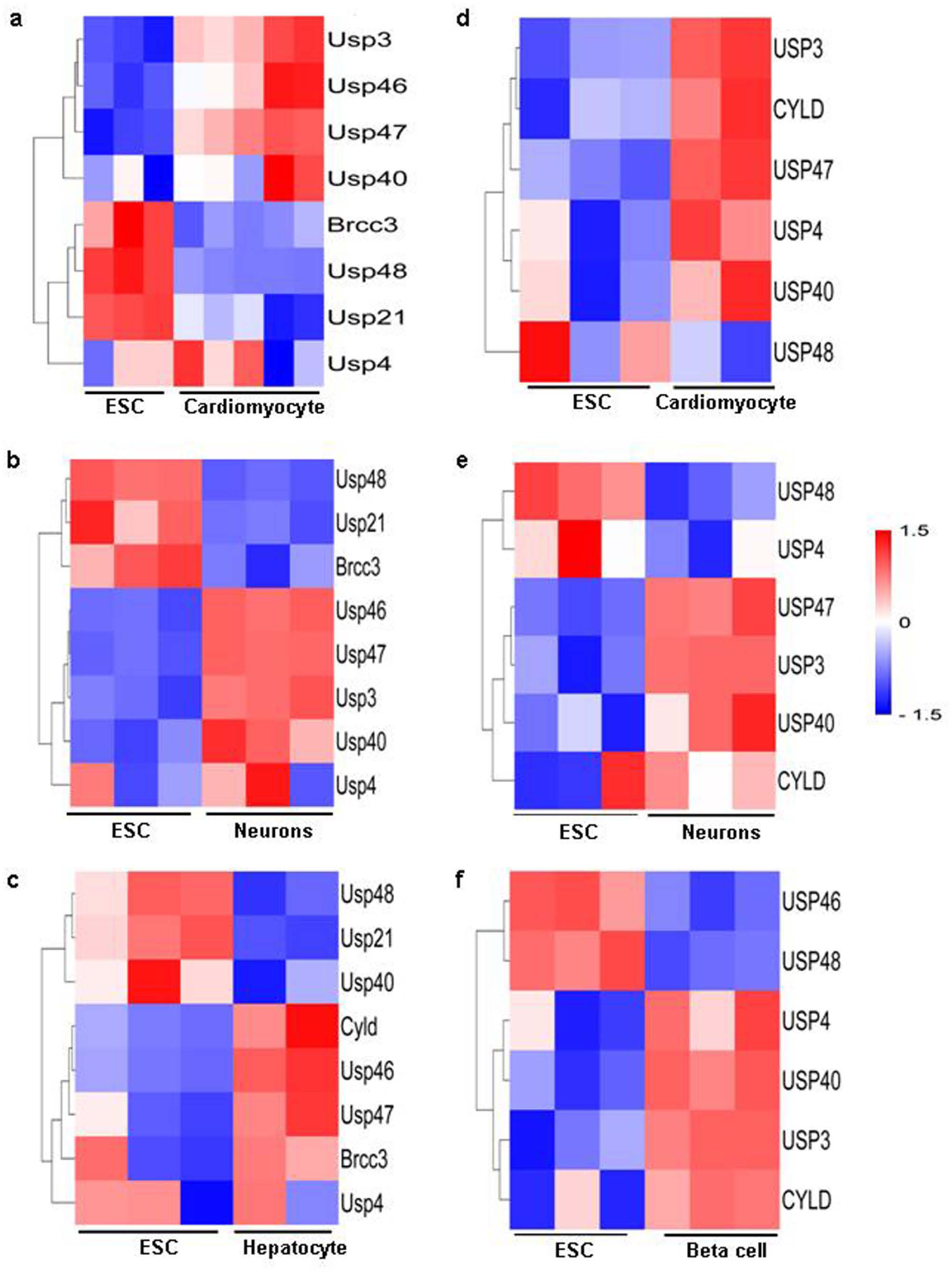
Differential Expression of selected DUBs during ESCs’ differentiation. Heatmap generated utilizing the information available from the murine (a-c) and human (d-f) datasets showing the differential expression of candidate DUBs in cardiomyocytes (a, d), neuron (b, e), hepatocytes (c) and pancreatic beta cells (f) differentiated from the respective ESCs. The comparison was done between ESC and the differentiated cells in each.

### Validation of DUB expression during cardiomyogenic and neural differentiation from ESCs *in vitro*

To validate the expression of selected DUBs (Usp46, Usp47, Usp40, Usp4, and Brcc3) emerged in our *in silico* predictions we considered murine ESCs (Fig. 4a), directed them to differentiate into cardiomyocytes (Fig. 4) and neural cells (Fig. 5) and carried out qRT-PCR during various stages of cardiomyogenic and neural differentiation from ESCs *in vitro*. The rationale being, the stage-specific differential expression during cardiac and neural differentiation would predict the likely involvement of the particular DUBs in the differentiation process and a temporal influence, if any. Fig. 4a shows the live microscopic images of ESCs and their differentiation into mesoderm (d2) and cardiomyogenic derivatives (d5 and d10) by generating EBs in hanging drop (d2) and suspension culture (d5) followed by their seeding for further differentiation into cardiomyocytes (d10). At d10 (shown as white arrow) a number of distinct pulsating clusters appeared proving them as cardiomyocytes. The validation of the same was carried out by observing the immunostained patterns, where cTnT revealed their cardiomyocyte phenotype (Fig. 4b). Similarly OCT4 indicated the undifferentiated ESCs, whereas Bry and PDGFRα reflected early mesoderm (d2) and cardiac progenitors (d5) respectively (Fig. 4b). In line with the *in silico* findings shown in Fig. 3, the expression of *Usp46*, *Usp47* and *Usp40* showed an increasing trend during cardiac differentiation when monitored on d2, d5 and d10 of ESCs’ differentiation, respectively (Fig. 4c-e). It is noteworthy that, the increased expression of these three DUBs during differentiation corroborated with their predicted roles in cardiac lineage specification shown in Fig. 3. While *Usp4* showed higher expression in ESCs compared to the differentiated cells, we did not find any remarkable difference in the expression of *Brcc3* among the time points tested (Fig. 4f, g). The inconsistency in Brcc3 expression during ESCs’ differentiation to cardiomyogenic lineage was in line with its variable expression status shown in Fig. 2 comparing the information on ESCs vs EBs from various retrieved datasets.

**Fig. 4:**
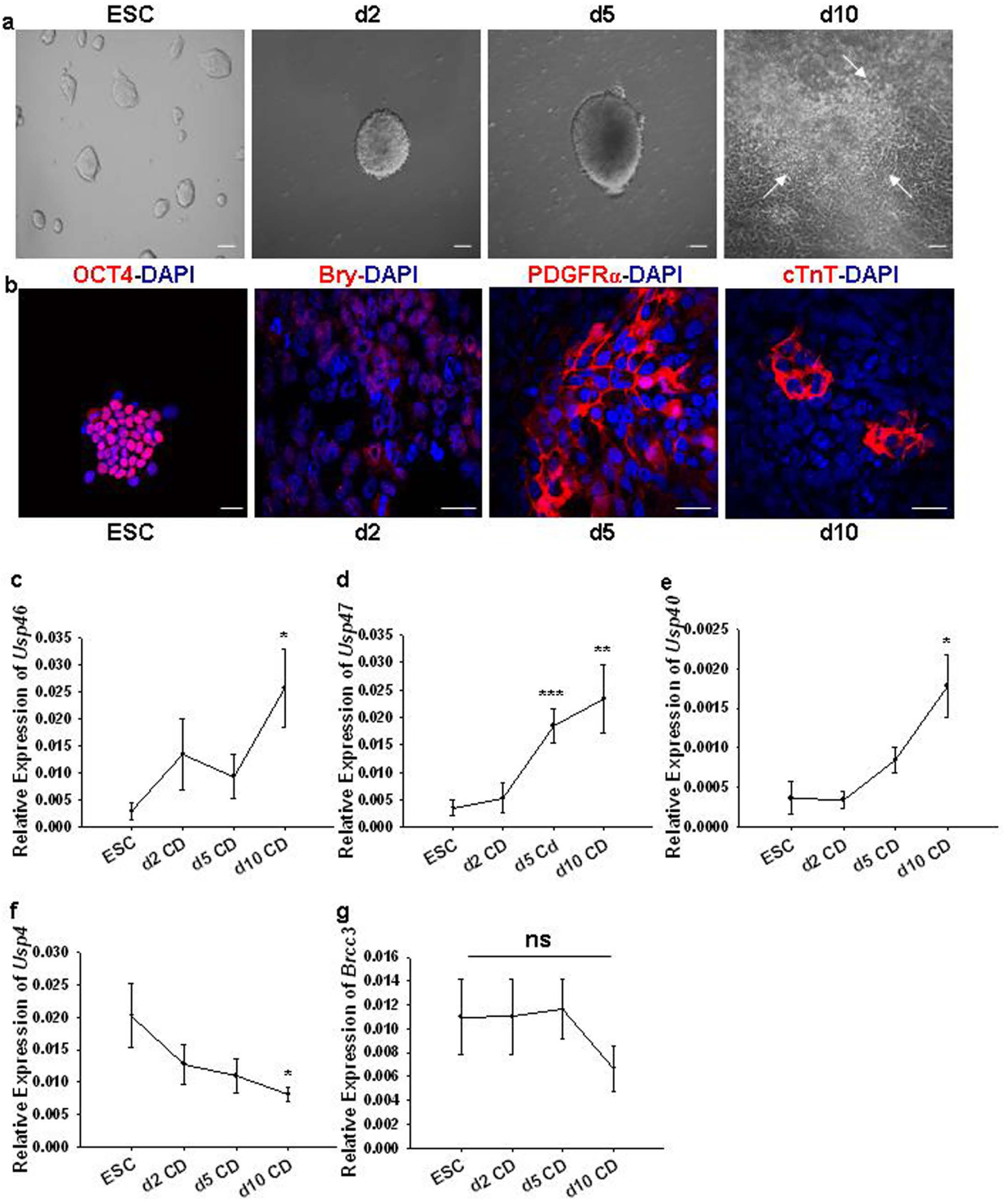
Transcriptional Validation of candidate DUBs during Cardiac Differentiation: a. Brightfield images showing live cells in culture; undifferentiated ESCs and EBs at various stages (d2, d5 and d10) of cardiac differentiation (white arrows in d10 EBs represent pulsating clusters reflecting cardiomyocytes). b) Confocal laser scanning microscopy (CLSM) validates the respective cell types via their positive immunostaining with cell-type specific antibodies (OCT4: undifferentiated ESCs; Bry: Mesoderm; PDGFRα: Cardiac progenitors; cTnT: Cardiomyocytes; DAPI: Nuclear stain). c-g) qRT-PCR showing expression patterns of various DUBs in murine ESCs in comparison with the same at various stages of cardiac differentiation. CD: Cardiac differentiation. Data shown are mean ± SEM; n=4. Statistical significance between the groups was assessed by Student’s *t*-test; ns: P > 0.05, *: P ≤ 0.05, **: P ≤ 0.01, ***: P ≤ 0.001.

**Fig. 5:**
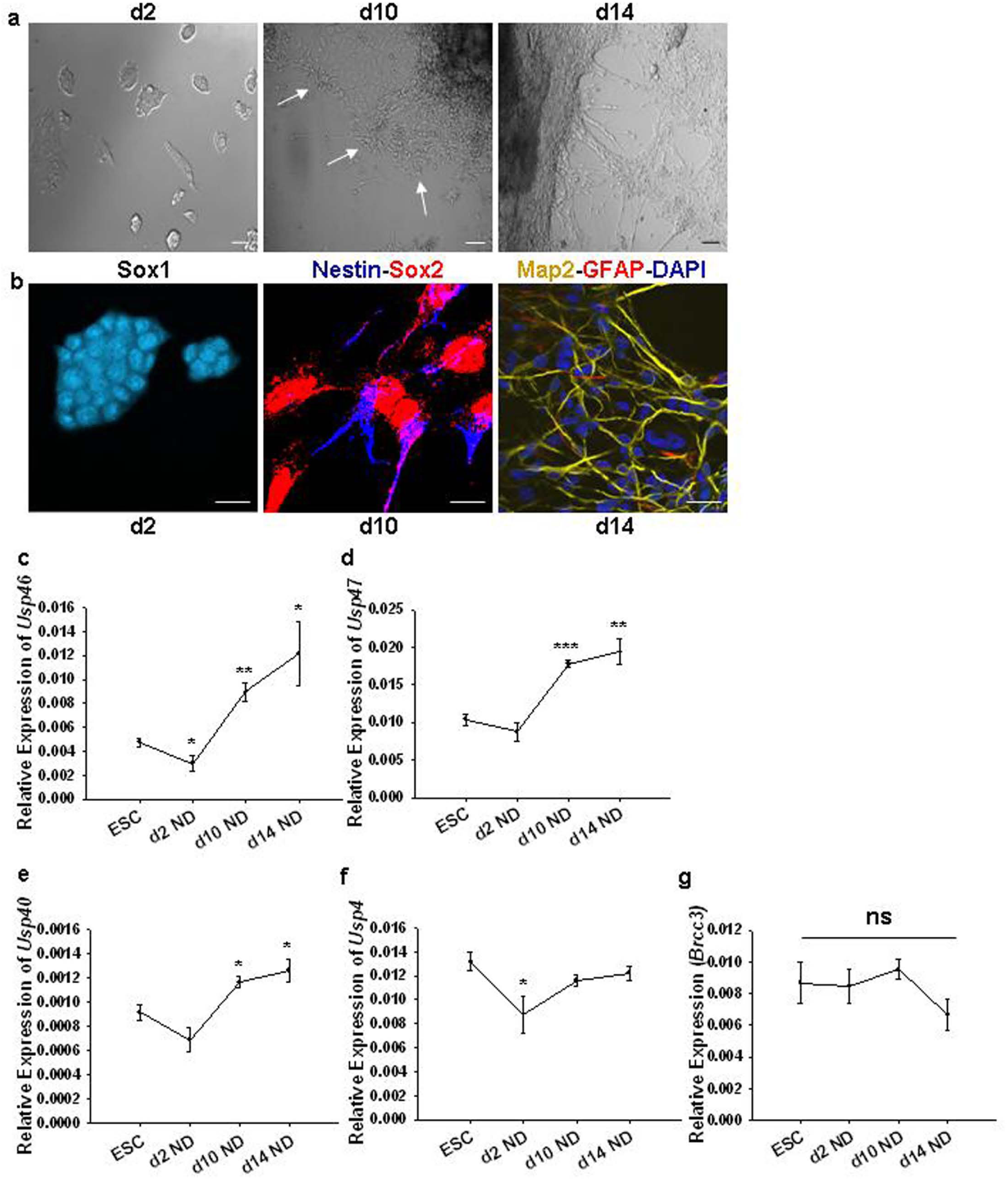
Transcriptional Validation of candidate DUBs during Neural Differentiation: a, b) Brightfield (a) and respective CLSM (b) images showing live cells in culture and their phenotypic validation by immunostaining at various stages (d2, d10 and d14) of neural differentiation (Upper Panel: white arrows in d10 indicate the neural rosettes reflecting their neural progenitor status; Lower Panel: Sox1: Early neural cells at induction; Nestin and Sox2: Neural stem cells/progenitors; Map2 and GFAP: Differentiated neurons and astrocytes respectively; DAPI: Nuclear stain). c-g) qRT-PCR showing expression patterns of various DUBs in murine ESCs in comparison with the same at various stages of neural differentiation. ND: Neural differentiation. Data shown are mean ± SEM; n=4. Statistical significance between the groups was assessed by Student’s *t*-test; ns: P > 0.05, *: P ≤ 0.05, **: P ≤ 0.01, ***: P ≤ 0.001.

In case of neural differentiation, a monolayer culture strategy was considered where the compact colony morphology of ESCs shown in Fig. 4a changed to more flattened ones (Fig. 5a: d2) upon neural induction and showed Sox1 positivity (Fig. 5b) reflecting early neural cells. During further differentiation of these cells, neural rosettes (white arrows: d10 upper panel) representing the neural stem cells appeared in culture and those were positive for both nestin and Sox2 (Fig. 5a, b). The cells exhibited distinct interconnected network structures by two weeks of neural differentiation (Fig. 5a: d14) that included both differentiated neurons (Map2^+^) and astrocytes (GFAP^+^) (fig. 5b). qRT-PCR was conducted to validate the *in silico* prediction of the candidate DUBs at the stated stages of neural differentiation from ESCs. While both *Usp47 and Usp40* showed similar expression in ESCs and at the neural differentiation induction stage (d2ND), there was a decrease in *Usp46* expression in d2ND (Fig. 5c-e). However, during further differentiation to neural progenitors (d10ND) and differentiated neural derivatives (d14ND), the three stated DUBs were significantly upregulated, corroborating with our *in silico*-curated analysis shown in Fig. 3b. In contrast, no striking alterations in the expression profile of *Usp4* and *Brcc3* was obtained across differentiation time regimens, except that on d2ND where *Usp4* level was lower compared to ESCs (Fig. 4f, g). Together, these experimental validations strengthened our *in silico* predictions and provided a framework for further investigation into their specific roles and regulation during stem cell maintenance and differentiation in future.

### Functional Annotation of Candidate DUBs

A multi-platform computational approach was undertaken to elucidate the role of the understudied selected candidate DUBs in the cell fate specification. Accordingly, by using the TRANS-DSI database, which employs machine learning to predict protein-protein interactions we could identify several novel DUB-substrate interactions (DSI) (Table S1). We included the predicted substrates of the selected DUBs with a confidence score ≥ 0.5 and cross-verified the same using the existing database in UbiBrowser 2.0 (data not shown). Further, the GO enrichment analysis of the candidate DUBs using the METASCAPE platform facilitated identification of their functional associations and revealed the key cellular processes that they might be involved in during the stem cell maintenance and differentiation in *Homo sapiens*. As expected, the ubiquitination and proteasomal-associated components emerged as the common interactors of all the selected candidate DUBs. Interestingly, the enriched substrates of USP46 were associated with critical cellular processes such as cell proliferation, cell division, hematopoiesis and neuritogenesis, etc. (Fig. 5a) suggesting its possible role during organismal development. In addition to this, the predicted interactome also revealed the likely involvement of USP46 with different signalling pathways including TGF-β, TNF-α and NOTCH. Similarly, USP47 was found to be associated with the regulation of protein phosphorylation and stability revealing interactions with crucial cellular processes like cell cycle, endocytosis and DNA repair among others (Fig. 5b). While DNA repair and pathways operational during viral infection and cellular stress response were common between USP4 and BRCC3, the former showed a bias towards immunological processes with its substrates related mostly to cytokine signalling, autophagy, antibody and interferon production, etc. (Fig. 5c, d). BRCC3 seemed to be associated with transcriptional and cell cycle regulation involving Tp53 (Fig. 5d). Interestingly, CYLD manifested divergent functional attributes as the enriched terms ranged from cancer pathways – MAPK cascade, apoptosis to immunological response pathway including NFκB and IL1 signalling, among others (Fig. 5e). Our analysis also revealed the well-established association of USP3 with chromatin organization among the myriads of other functions (Fig. 5f). Incidentally, the GO analysis of these six DUBs underscored their link with various disease processes including cancer. Hence, it provided a stepping stone for the future experimental validation of their specific role in stem cell fate specification and cancer.

### DUBs and Cancer

We also accessed the TCGA (The Cancer Genome Atlas Program) database through the UALCAN portal to determine the differential association of candidate DUBs in cancer (Fig. S1). While USP46 expression was upregulated in 12 out of the 24 cancer types analyzed with no difference in sarcoma, USP47 was upregulated in 9 with no difference in renal (KICH) cancer (Fig. S1a, b). BRCC3 was found to be upregulated in majority of the cancer types (16 out of 24) and downregulated in 5 others (Fig. S1d), possibly due to its association with Tp53 and cell cycle regulation shown in Fig. 5d. Both USP4 and CYLD were upregulated in 7 different cancers each and showed downregulation in 15 and 17 different cancers respectively (Fig. S1c, e). Among the selected DUBs, most of those were downregulated in cancers of renal, uterine, prostrate, rectal and thymic origin and upregulated in liver, colon, esophageal cholangiocarcinoma and skin cancers. While USP46 and BRCC3 were upregulated in bladder, breast cervical, lung and pancreatic cancers, USP47, USP4 and CYLD were downregulated. In contrast, in head and neck and neuroendocrine cancers, USP47 and CYLD were upregulated, whereas USP46 and USP4 were downregulated. Together our data indicated differential expression of DUBs across various cancers and future investigation would elucidate their specific relevance with respect to cancer prevalence, onset and progression.

### DUBs Signaling Pathway Crosstalk Analysis – *In silico* prediction

Considering the multitude of DUB interactomes and their association with development and disease, determined and predicted during our *in silico* analysis, we undertook further analysis to identify the DUBs that are regulating the core developmental and oncogenic pathways. Using UbiBrowser 2.0, a predictive tool for ubiquitination network mapping, our analysis prioritized transcriptional activators and mediators across four evolutionarily conserved pathways - Wnt, Notch, Hedgehog, and Hippo, the rationale being the surge of information available for *Homo sapiens* (Fig. 3) and *Mus musculus* (Fig. S2). Moreover, we have focused on specific components of each pathway to identify DUBs that may play crucial roles in their regulation. Among the known DUBs that are associated with Wnt signalling (Fig. 6a, b), USP7 was found to be a common candidate targeting both CTNBB1 (β-catenin) and AXIN1, the two key players of the canonical Wnt pathway. Similarly, among the predicted DUBs, CYLD, USP2, USP25 and USP45 emerged as the common ones for both. While both USP9X and USP14 appeared as the common DUBs for CTNBB1, the same emerged under the predicted category for AXIN1. Concerning the Notch pathway, we investigated only NOTCH1-DUB interactome as other NOTCH receptors did not yield significant predictions. Similar to that seen with Wnt, NOTCH1 and YAP1 were also identified as the known targets of USP7 (Fig. 6c, e). Moreover, USP9X emerged as a common DUB for YAP1 and LATS1, both associated with Hippo signalling (Fig. 6e, f). While USP25 and USP28 emerged as predicted DUBs for CTNBB1/AXIN1, Notch1 was found to be a known target of both these DUBs (Fig. 6a-c). Our analysis also revealed that in human while USP7 was found to be associated with almost all the stated signalling pathways, CYLD appeared to be a common DUB among the predicted ones (Fig. 6a-d, f). Concerning Smoothened (SMO), a component of Hedgehog pathway, most of the DUBs including USP7 belonged to the predicted category except USP8 and UCHL5 (Fig. 6d). Similar observation on the association of predicted DUBs and signalling pathway components were also obtained in *Mus musculus* species by utilizing the UbiBrowser 2.0. analysis tools. As seen in Fig. S2, no known DUBs were found in mouse database that were targeting Wnt and Hedgehog signalling. However, the predicted ones shared the common DUBs that were indentified under the known category (USP4, USP7, USP9x, USP14, USP20, and USP47) targeting CTNNB1 in human (Fig. 6a and Fig. S2a). Interestingly, all the predicted DUBs for Ctnnb1 in mouse were also common to human thereby indicating their likely evolutionarily conserved substrate target. While CYLD, USP46 and USP14, emerged as common predicted DUBs for both Ctnnb1 and Axin1 as possible targets in mouse, CYLD and USP14 were the common DUBs for AXIN1 in both the species (Fig. 6a, b and Fig. S2a, b). Similarly, Usp10 appeared as the sole DUB having Notch1 as its known target in mouse, whereas USP7 that was known to have NOTCH1 as its target in human emerged under the predicted category (Fig. 6c and Fig. S2c). Among the predicted ones USP1, USP5, USP15 and UCHL1 were seen as common between both the species. Concerning the Hedgehog signalling, many of the predicted DUBs including CYLD were common between both mouse and human (Fig. 6d and Fig. S2d). CYLD also emerged as a common DUB for LATS1 falling under the predicted category in both the species studied (Fig. 6f and Fig. S2f). While USP7 was identified as the known DUB for YAP1 common to both the species, Outb1 emerged as the common predicted DUB in them (Fig. 6e and Fig. S2e). Undoubtedly, these findings have generated resources and avenues for further exploration to delineate the mechanistic basis concerning specific DUBs in relation to organismal development and disease-specific attributes.

**Fig. 6:**
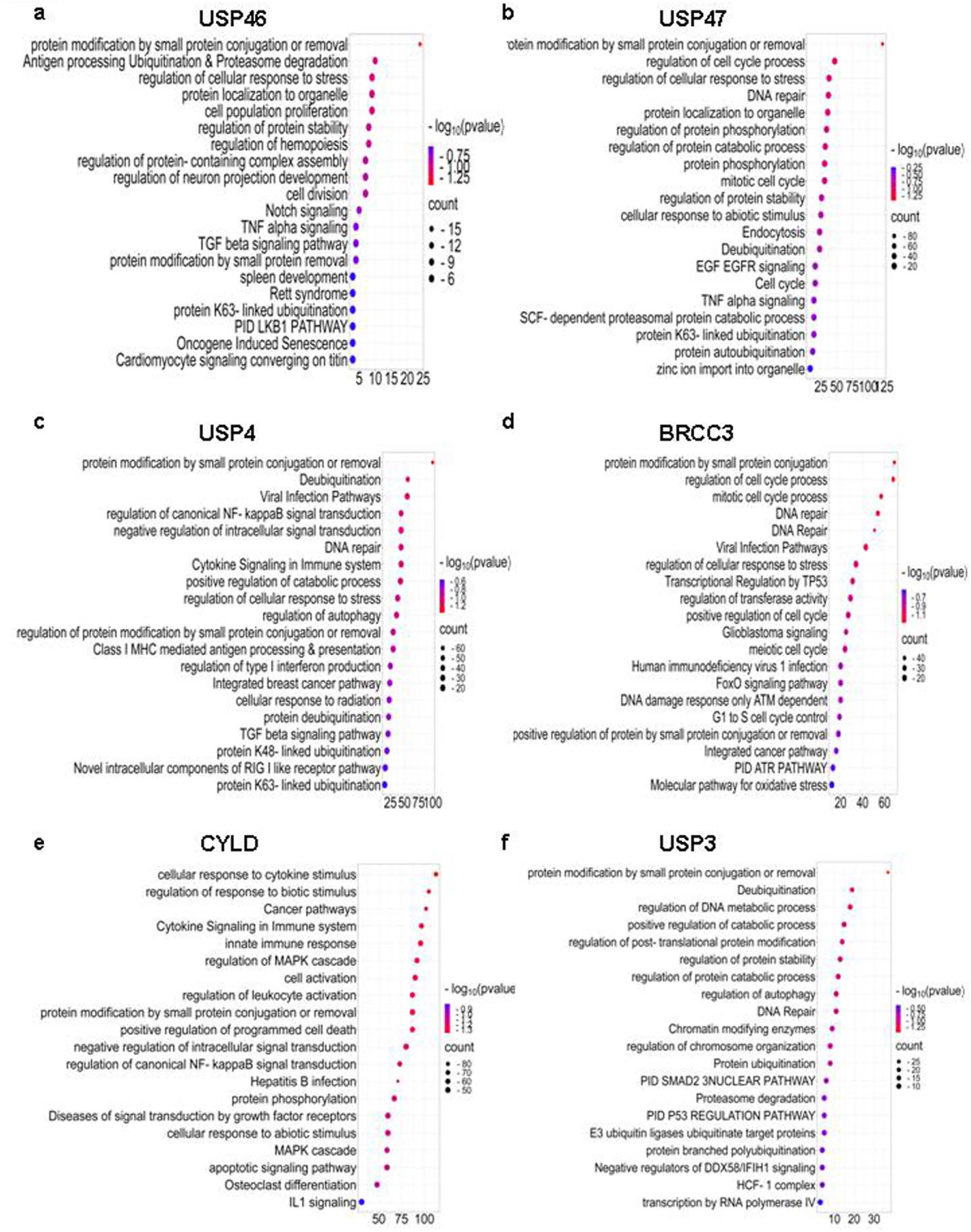
GO analysis for the predicted substrates of the candidate DUBs. The predicted substrates of the candidate DUBs using Trans-DSI database (human) were based on the assigned confidence score ≥ 0.5. The GO analysis of the predicted substrates was performed using Metascape by taking various biological processes into consideration for USP46 (a), USP47 (b), USP4 (c) BRCC3 (d), CYLD (e) and USP3 (f) respectively. GO analysis indicates that the DUBs are involved in various critical cellular processes including cell cycle, development, autophagy and transcriptional regulation.

**Fig. 7:**
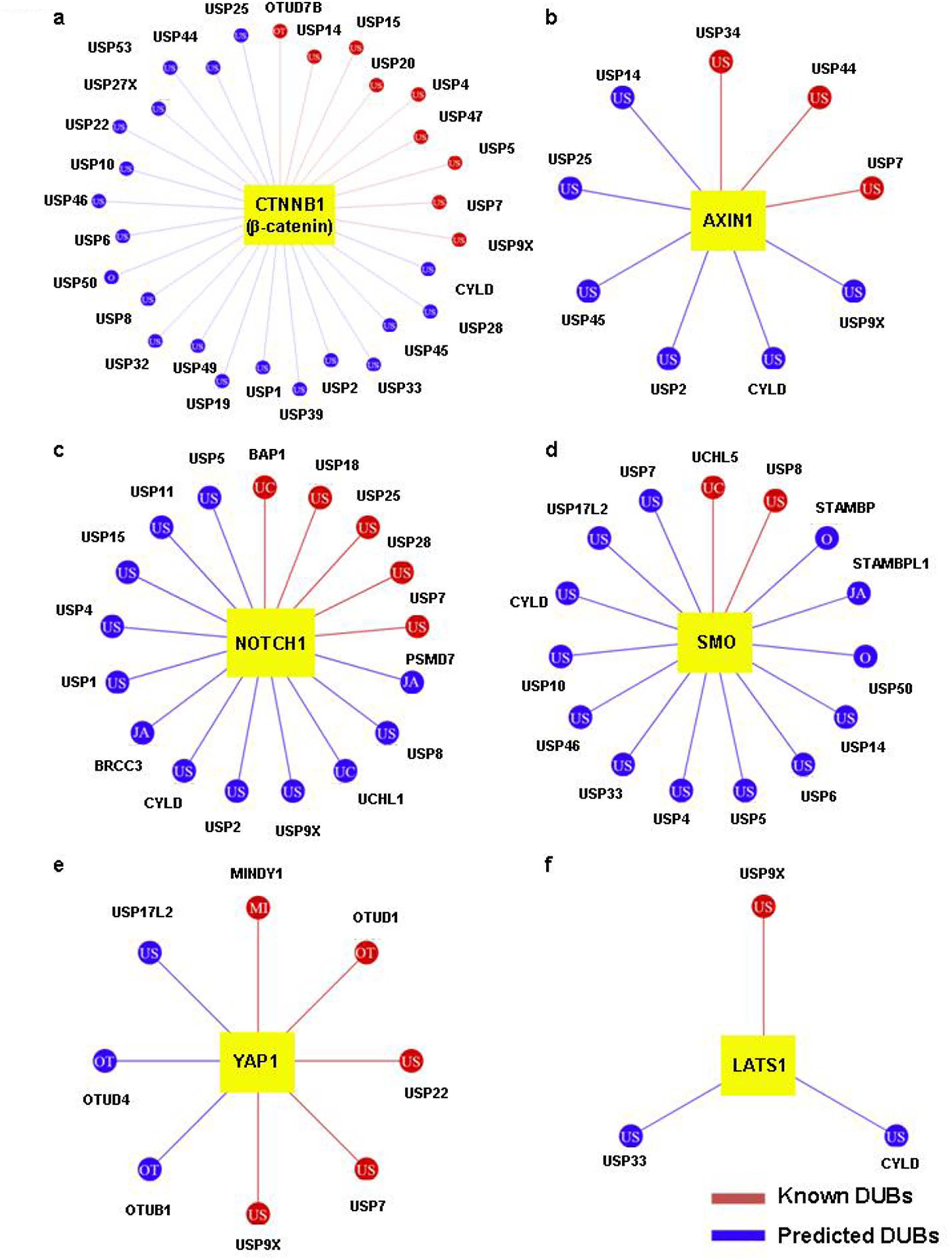
Analysis of DUBs associated with various signalling pathways in *Homo sapiens* using Ubibrowser 2.0. a, b) Analysis of the predicted and known DUBs associated with CTNNB1 (β-catenin) and AXIN1, the Wnt signalling pathway components. c, d) DUBs associated with NOTCH1 (c) and SMO (d) belonging to Notch and Hedgehog signalling respectively. e, f) Components of Hippo signalling pathway – YAP1 and LATS1 and the associated DUBs. The known interactions are indicated in red color and predicted interactions in blue. The confidence score threshold was set at ≥ 0.5 to ensure high confidence predictions.

## Discussion

Ubiquitination and deubiquitination are critical post-translational modifications involved in key cellular processes underlying early development and various pathologies. DUBs are known to exert their functions through interplay with various signalling cascades and modulation of protein stability and also contribute to cell cycle regulation, stem cell maintenance and their differentiation [1, 31–33]. Infact, a fine-tuned balance between ubiquitination-deubiquitination is crucial for various physiological processes [34]. However, unlike ubiquitination, the information on the regulatory roles of DUBs underlying both normal development and pathogenesis is limited. Hence, it is imperative to narrow this knowledge gap and address their functional and regulatory relevance by employing an integrated/interdisciplinary approach. Herein we have considered multiple computational approaches to undertake comprehensive analysis of DUBs associated with pluripotent stem cells as well as cancer and provided a prediction framework on their role underlying stem cell fate decision machinery.

In this study, we have utilized *in silico* predictions, transcriptomic profiling, and experimental validation to unravel the roles of DUBs in cell fate determination. The elucidation of USP3 in our previous study [30] and also the ongoing study with USP31 (to be published elsewhere) as potential cell fate modulators encouraged us further to conduct a comprehensive analysis on the association of DUBs during early development and cancer by utilizing the available datasets. Conducting *in silico* analysis has indeed helped identify several DUBs, both the known and the predicted ones, involved in cell fate decision. Moreover, the Stemmapper analysis has revealed the stage-specific expression of DUBs that act as molecular switches during lineage specification. An elegant study by Vlasschaert et al. [35] has demonstrated the evolutionary conservation along the DUB repertoire, including the orthologs and paralogs, by examining their evolutionary origin across species and functional diversification. In line with the same, our analysis involving mouse and human datasets during ESCs differentiation revealed an evolutionary conservation of several DUBs, particularly USP47, USP40, and USP48 between the two species. This conservation suggests the role of DUBs in stem cell maintenance and differentiation and underscores their fundamental roles in early development [11, 31, 32]. We have also validated some of them during both cardiomyogenic and neural differentiation from murine ESCs. Their distinct expression pattern suggests that DUBs may have context-dependent roles during differentiation into distinct lineages. Dysregulation of these processes can lead to developmental abnormalities and contribute to tumorigenesis, emphasizing the importance of understanding DUB functions in both development and disease [19]. We have noticed both commonality and striking diversity among the DUB candidates having pluripotency-associated factors as their targets. While cMYC and KLF4 are the known oncogenes, POU5F1, SOX2 and NANOG are associated with embryonic as well as many of the adult stem cells including cancer stem cells [36–38]. Hence, this differential mode of DUB interaction in stem cells envisaged their likely context-dependent functional attributes in rendering stemness during development and cancer onset/progression. Interestingly, the expression of USP48 from the curated datasets was seen to be comparatively higher in the undifferentiated ESCs than their differentiation counterparts and thereby hinted at their likely association with the pluripotent state maintenance. Similarly, the contrasting expression pattern of USP46 seen in the hepatocytes and pancreatic islet cells, the two distinct endodermal derivatives, compared to their respective ESC counterparts suggested its plausible association with either differential endodermal fate modulation or species-specific contrasting response. Such temporal regulations highlight the significance of DUBs in fine-tuning signalling pathways to govern cell fate decisions. Our analysis also corroborated with the established role of USP21 in ESC maintenance [29]. Together, our in vitro findings and *in-silico* predictions have provided the groundwork for further investigation to assess the specific modulations of key DUBs during early development.

The consistent upregulation pattern of USP46, USP47 and USP40 in ESC-derived cardiomyocytes and neurons strongly suggested their involvement in lineage specification. Incidentally, USP46, the histone DUB, is reported to be linked with gastrulation proceedings [39] and also participates in the regulation of AMPAR neurotransmitter turnover and synaptic plasticity [40]. Moreover, in contrast to its positive manifestation concerning Wnt signalling pathway by deubiquitinating LRP6 [41], it serves as a negative regulator of the AKT signalling pathway [42]. Similarly, USP47 is reported to promote cancer progression via activation of Wnt and Hippo signalling pathways [43, 44], whereas USP40 is also known to stabilize Claudin1 and promote hepatocellular carcinoma [45]. However, the roles of these DUBs in the context of stem cell maintenance or differentiation are not well studied. We demonstrated that USP40, USP46, and USP47 could be involved in cardiomyogenic and neural differentiation from ESCs based on our *in silico* curation and gene expression validation studies. Incidentally, this aligned with the predicted roles of USP46 and USP47 in Wnt/β-catenin pathway modulation in the context of cell fate determination [41, 44]. However, our *in vitro* validation on USP4 status during cardiac and neural differentiation revealed a contrasting response. While the *in silico* analysis among the curated datasets in mouse revealed an inconsistency concerning *Usp4* expression in cardiomyocytes and neurons compared to ESCs during maintenance, our validation study revealed a down-ward trend during cardiomyogenic differentiation from ESCs. Further exploration on these candidates would pave way to gain insight into their precise modulation and association with various signalling pathways that dictate cardiomyogenic and neurogenic proceedings. Concerning BRCC3, although our *in vitro* validation did not yield a concrete pattern during ESCs’ differentiation into cardiomyocytes and neural cells, further investigation on BRCC3 that has been studied in the context of DNA repair earlier [46] may uncover a possible link between its DNA damage response and stem cell maintenance. The same might hold true for its influence on cancer as it was seen overexpressed in majority of the cancers in TCGA database.

Our analysis also revealed that several DUBs, including USP46, USP47, USP4, USP40, CYLD, and BRCC3, are potential regulators of multiple developmental signalling pathways, such as Wnt, Notch, Hedgehog, and Hippo. This suggests a complex regulatory network where DUBs orchestrate cell fate decisions by coordinated modulation of multiple signalling pathways. Interestingly, CYLD, USP2, USP7, USP9X, USP25 and USP45 were found to be associated with both β-catenin and AXIN1. Similar cross-talk between DUBs and signalling pathways has been observed in other studies, where DUBs like USP9X and USP7 have been implicated in early embryonic and neuronal development, and are also reported to play dual roles in serving as both oncogenes and tumour suppressors, depending on the type of cancer [12, 13]. Similarly, a recent report indicated that USP28 modulates Notch signalling by stabilizing NICD1 and affects cancer progression and stem cell differentiation [48]. Our analysis has also revealed NOTCH1 as a target for USP7, USP25 and USP28. These findings also highlighted the complex interplay between DUBs and the key signalling pathways involved in embryonic development and cancer. In the similar line, our analysis on the expression of the candidate DUBs across various cancer types revealed a differential expression pattern across cancer types. The decreased expression of USP4 and CYLD in majority of the cancer types is in alignment with previous studies suggesting their tumour suppressor role [49, 50]. However, the varied expression of USP46 and USP47 in different cancer types is an indicator of the context-dependent role of these DUBs. USP4 and USP47 exhibited reduced expression in lung adenocarcinoma (LUAD) and colorectal tumors, consistent with their reported Wnt/β-catenin regulatory roles [51, 52]. However, USP46 displayed context-dependent expression aligning with its dual regulatory roles reported in suppression of AKT signalling [42] and promotion of Wnt signalling through LRP6 deubiquitination [44]. BRCC3 showed elevated expression in majority of the cancers and it has been reported to activate NF-κB in bladder cancer [53]. On the contrary, CYLD, the tumor suppressor, was downregulated in multiple cancers. We believe further investigation on these DUBs would help understand the complex dynamics and identify specific therapeutic candidates for preventing cancer progression, remission and other developmental disorders. Such multipronged approaches would also help generate new hypotheses and aid in further experimental investigations in understanding the precise regulatory roles of DUBs.

### Future Prospects

Our study has provided a predictive analysis on the DUB expression and function during stem cell maintenance and differentiation. We have identified several promising candidates with potential roles in cell fate specification for further investigation and lay the groundwork for future studies aimed at elucidating their molecular functions. Further validation of the same would also be required to gain a concrete insight into their precise mode of regulation. Moreover, studies focusing on understanding the specific substrates and mechanisms of action for these DUBs in stem cell maintenance and lineage commitment are essential. Hence, there is a need for further studies to dissect the context-dependent functions of DUBs, where investigating the interplay between DUBs and other post-translational modifications during differentiation would reveal new facets of regulations in cell fate decisions. These studies may also uncover novel therapeutic targets for further exploration and intervention in regenerative medicine and cancer therapy. Undoubtedly, the integration of predictive modeling with functional studies would provide a blueprint for targeting DUB networks and devising pertinent strategies in the stated domains.

## Supporting information

Suuplementary Figures- Fig.S1 and S2

## Acknowledgements

The authors wish to acknowledge the support received from NCCS intramural funding. FM is a graduate student supported by fellowship from Dept. of Biotechnology, Govt. of India.

## Ethical Approval

Not applicable

## Consent to Participate

Not applicable

## Consent to Publish

Not applicable

## Author Contribution

FM: Execution, Data Collection, Analysis, Writing, Editing and Finalization of the manuscript; NL: Conceptualization, Resource generation, Project supervision, Execution, Data Collection, Analysis, Writing, Editing and Finalization of the manuscript

## Data Availability

The datasets and analysis related to this article shall be made available from the corresponding author on reasonable request.

## Figure Legends

**Fig. S1. Pan-cancer analysis of DUBs.**

The box plot depiction of candidate DUBs in different cancers retrieved from TGCA data base (a: USP46, b: USP47, c: USP4, d: BRCC3, and e: CYLD). The blue colour represents the normal samples and the red, the tumour.

**Fig. S2. Analysis of DUBs associated with various signalling pathways in *Mus musculus us*ing Ubibrowser 2.0.** a, b) Analysis of the predicted and known DUBs associated with Wnt signalling pathway components; β-catenin and Axin1. c, d) DUBs associated with Notch (Notch1) and Hedgehog (Smo) signalling respectively. e, f) Components of Hippo signalling pathway – Yap1 and Lats1 and the associated DUBs. The known interactions are indicated in Red color and the predicted interactions in blue. The confidence score threshold was set at ≥ 0.5 to ensure the high confidence predictions.

