## Supplementary figures and images for "*In silico* analysis and Predictive linkage of Deubiquitinating Enzymes underlying Early Development"

### Suuplementary Figures- Fig.S1 and S2

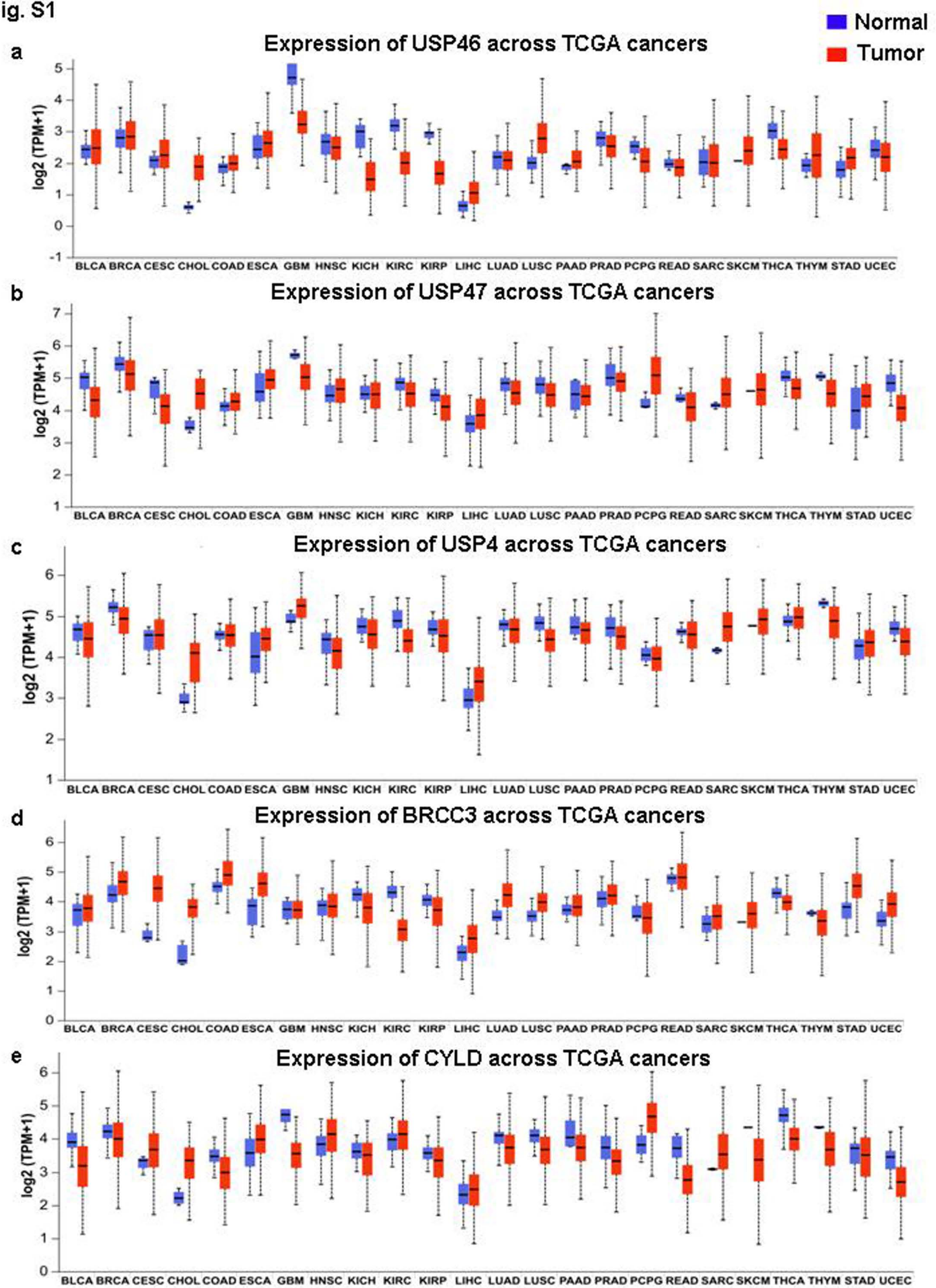

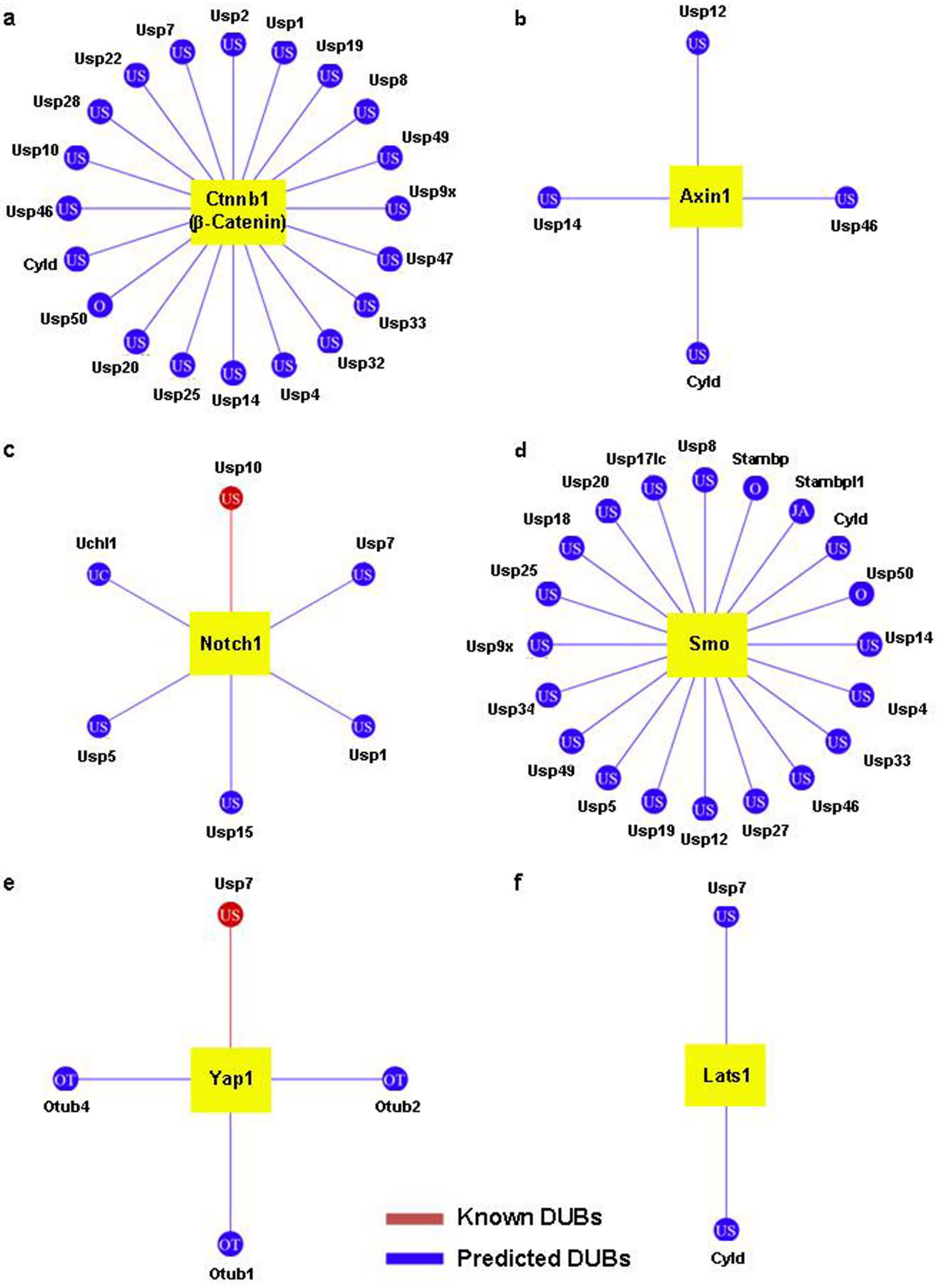
