## Supplementary Tables - Table S1 and S2 for "*In silico* analysis and Predictive linkage of Deubiquitinating Enzymes underlying Early Development"

**Prediction of novel DUB-Substarte Interaction using TRANS DSI platform**

| SwissProt ID (DUB) | SwissProt ID (SUB) | Entrez Gene Symbol (DUB) | Entrez Gene Symbol (SUB) | TransDSI Score |
|---|---|---|---|---|
| O00487 | P01106 | PSMD14 | MYC | 0.945 |
| O15372 | P01106 | EIF3H | MYC | 0.945 |
| O75317 | P01106 | USP12 | MYC | 0.959 |
| O75604 | P01106 | USP2 | MYC | 0.952 |
| O94782 | P01106 | USP1 | MYC | 0.965 |
| O94966 | P01106 | USP19 | MYC | 0.956 |
| O95630 | P01106 | STAMBP | MYC | 0.920 |
| P09936 | P01106 | UCHL1 | MYC | 0.962 |
| P12956 | P01106 | XRCC6 | MYC | 0.885 |
| P21580 | P01106 | TNFAIP3 | MYC | 0.956 |
| P35125 | P01106 | USP6 | MYC | 0.954 |
| P40818 | P01106 | USP8 | MYC | 0.933 |
| P45974 | P01106 | USP5 | MYC | 0.933 |
| P51784 | P01106 | USP11 | MYC | 0.958 |
| P54252 | P01106 | ATXN3 | MYC | 0.961 |
| P54578 | P01106 | USP14 | MYC | 0.965 |
| P62068 | P01106 | USP46 | MYC | 0.961 |
| Q0WX57 | P01106 | USP17L24 | MYC | 0.927 |
| Q13107 | P01106 | USP4 | MYC | 0.957 |
| Q14694 | P01106 | USP10 | MYC | 0.940 |
| Q5D1E8 | P01106 | ZC3H12A | MYC | 0.912 |
| Q5VVJ2 | P01106 | MYSM1 | MYC | 0.956 |
| Q6R6M4 | P01106 | USP17L2 | MYC | 0.920 |
| Q86UV5 | P01106 | USP48 | MYC | 0.949 |
| Q8N5J2 | P01106 | MINDY1 | MYC | 0.956 |
| Q92560 | P01106 | BAP1 | MYC | 0.965 |
| Q92905 | P01106 | COPS5 | MYC | 0.939 |
| Q93008 | P01106 | USP9X | MYC | 0.925 |
| Q96FW1 | P01106 | OTUB1 | MYC | 0.974 |
| Q9H0E7 | P01106 | USP44 | MYC | 0.965 |
| Q9HBJ7 | P01106 | USP29 | MYC | 0.944 |
| Q9NQC7 | P01106 | CYLD | MYC | 0.965 |
| Q9UHP3 | P01106 | USP25 | MYC | 0.940 |
| Q9UMW8 | P01106 | USP18 | MYC | 0.962 |
| Q9UPU5 | P01106 | USP24 | MYC | 0.918 |
| Q9Y4E8 | P01106 | USP15 | MYC | 0.961 |
| Q9Y5K5 | P01106 | UCHL5 | MYC | 0.966 |
| P09936 | P48431 | UCHL1 | SOX2 | 0.925 |
| P12956 | P48431 | XRCC6 | SOX2 | 0.751 |
| Q70CQ2 | P48431 | USP34 | SOX2 | 0.899 |
| Q93009 | P48431 | USP7 | SOX2 | 0.903 |
| Q9C040 | P48431 | TRIM2 | SOX2 | 0.875 |

| | | | | |
|---|---|---|---|---|
| Q9H0E7 | P48431 | USP44 | SOX2 | 0.933 |
| P21580 | O43474 | TNFAIP3 | KLF4 | 0.881 |
| Q53GS9 | O43474 | USP39 | KLF4 | 0.923 |
| Q93009 | O43474 | USP7 | KLF4 | 0.843 |
| Q96RU2 | O43474 | USP28 | KLF4 | 0.863 |
| P09936 | Q01860 | UCHL1 | POU5F1 | 0.964 |
| P12956 | Q01860 | XRCC6 | POU5F1 | 0.906 |
| Q93008 | Q01860 | USP9X | POU5F1 | 0.918 |
| P09936 | Q9H9S0 | UCHL1 | NANOG | 0.964 |
| P12956 | Q9H9S0 | XRCC6 | NANOG | 0.896 |
| P40818 | Q9H9S0 | USP8 | NANOG | 0.931 |
| P54252 | Q9H9S0 | ATXN3 | NANOG | 0.967 |
| Q93008 | Q9H9S0 | USP9X | NANOG | 0.908 |
| Q93009 | Q9H9S0 | USP7 | NANOG | 0.940 |
| Q9BXU7 | Q9H9S0 | USP26 | NANOG | 0.958 |
| Q9H0E7 | Q9H9S0 | USP44 | NANOG | 0.966 |
| O00487 | P35222 | PSMD14 | CTNNB1 | 0.904 |
| P09936 | P35222 | UCHL1 | CTNNB1 | 0.934 |
| P12956 | P35222 | XRCC6 | CTNNB1 | 0.861 |
| P21580 | P35222 | TNFAIP3 | CTNNB1 | 0.949 |
| P35125 | P35222 | USP6 | CTNNB1 | 0.918 |
| Q5VVJ2 | P35222 | MYSM1 | CTNNB1 | 0.929 |
| Q92905 | P35222 | COPS5 | CTNNB1 | 0.896 |
| Q9C040 | P35222 | TRIM2 | CTNNB1 | 0.923 |
| Q9NQC7 | P35222 | CYLD | CTNNB1 | 0.961 |
| Q9UGI0 | P35222 | ZRANB1 | CTNNB1 | 0.932 |
| Q9Y5E6 | P35222 | PCDHB3 | CTNNB1 | 0.904 |
| P40818 | P56704 | USP8 | WNT3A | 0.781 |
| O00487 | O15169 | PSMD14 | AXIN1 | 0.921 |
| O95630 | O15169 | STAMBP | AXIN1 | 0.872 |
| P21580 | O15169 | TNFAIP3 | AXIN1 | 0.933 |
| Q8N5J2 | O15169 | MINDY1 | AXIN1 | 0.941 |
| Q92905 | O15169 | COPS5 | AXIN1 | 0.915 |
| Q96RU2 | O15169 | USP28 | AXIN1 | 0.924 |
| Q9C040 | O15169 | TRIM2 | AXIN1 | 0.925 |
| Q9NQC7 | O15169 | CYLD | AXIN1 | 0.948 |
| Q9UGI0 | O15169 | ZRANB1 | AXIN1 | 0.920 |
| Q9UHP3 | O15169 | USP25 | AXIN1 | 0.913 |
| Q9Y4E8 | O15169 | USP15 | AXIN1 | 0.940 |
| O00487 | Q9Y2T1 | PSMD14 | AXIN2 | 0.898 |
| Q9C040 | Q9Y2T1 | TRIM2 | AXIN2 | 0.903 |

| | | | | |
|---|---|---|---|---|
| Q9UHP3 | Q9Y2T1 | USP25 | AXIN2 | 0.887 |
| Q5VVJ2 | Q04912 | MYSM1 | MST1R | 0.819 |
| P15374 | P46937 | UCHL3 | YAP1 | 0.789 |
| P54578 | P46937 | USP14 | YAP1 | 0.815 |
| Q13107 | P46937 | USP4 | YAP1 | 0.789 |
| Q5VVJ2 | P46937 | MYSM1 | YAP1 | 0.775 |
| Q70CQ2 | P46937 | USP34 | YAP1 | 0.810 |
| Q70EL4 | P46937 | USP43 | YAP1 | 0.750 |
| Q86UV5 | P46937 | USP48 | YAP1 | 0.753 |
| Q96K76 | P46937 | USP47 | YAP1 | 0.720 |
| Q96RU2 | P46937 | USP28 | YAP1 | 0.769 |
| Q9UHP3 | P46937 | USP25 | YAP1 | 0.762 |
| Q9UMW8 | P46937 | USP18 | YAP1 | 0.804 |
| Q9UPU5 | P46937 | USP24 | YAP1 | 0.721 |
| Q5VVJ2 | O95835 | MYSM1 | LATS1 | 0.864 |
| Q5VVQ6 | O95835 | YOD1 | LATS1 | 0.885 |
| Q92560 | O95835 | BAP1 | LATS1 | 0.876 |
| Q96RU2 | O95835 | USP28 | LATS1 | 0.859 |
| Q5VVJ2 | Q9NRM7 | MYSM1 | LATS2 | 0.829 |
| Q92560 | Q9NRM7 | BAP1 | LATS2 | 0.837 |
| Q93009 | Q9NRM7 | USP7 | LATS2 | 0.792 |
| Q96RU2 | Q9NRM7 | USP28 | LATS2 | 0.823 |
| Q93009 | Q9Y6K1 | USP7 | DNMT3A | 0.548 |
| Q93009 | Q9UBC3 | USP7 | DNMT3B | 0.611 |
| Q9H0E7 | Q9UBC3 | USP44 | DNMT3B | 0.569 |
| Q4G0A6 | Q9BRJ9 | MINDY4 | MESP1 | 0.564 |
| P09936 | P16234 | UCHL1 | PDGFRA | 0.865 |
| Q93009 | P16234 | USP7 | PDGFRA | 0.809 |
| A6NNY8 | Q96QV6 | USP27X | H2AC1 | 0.820 |
| P46736 | Q96QV6 | BRCC3 | H2AC1 | 0.785 |
| Q5VVJ2 | Q96QV6 | MYSM1 | H2AC1 | 0.821 |
| Q92560 | Q96QV6 | BAP1 | H2AC1 | 0.856 |
| Q96F44 | Q96QV6 | TRIM11 | H2AC1 | 0.802 |
| Q9C040 | Q96QV6 | TRIM2 | H2AC1 | 0.757 |
| Q9UK80 | Q96QV6 | USP21 | H2AC1 | 0.768 |
| Q9UPT9 | Q96QV6 | USP22 | H2AC1 | 0.868 |
| Q9Y5T5 | Q96QV6 | USP16 | H2AC1 | 0.825 |
| Q9Y6I4 | Q96QV6 | USP3 | H2AC1 | 0.797 |
| P12956 | Q96QV6 | XRCC6 | H2AC1 | 0.640 |

| | | | | |
|---|---|---|---|---|
| A6NNY8 | O60814 | USP27X | H2BC12 | 0.874 |
| O94966 | O60814 | USP19 | H2BC12 | 0.875 |
| P35125 | O60814 | USP6 | H2BC12 | 0.867 |
| P46736 | O60814 | BRCC3 | H2BC12 | 0.842 |
| P51784 | O60814 | USP11 | H2BC12 | 0.877 |
| Q13107 | O60814 | USP4 | H2BC12 | 0.864 |
| Q5VVJ2 | O60814 | MYSM1 | H2BC12 | 0.873 |
| Q70CQ1 | O60814 | USP49 | H2BC12 | 0.883 |
| Q70CQ3 | O60814 | USP30 | H2BC12 | 0.889 |
| Q70EK9 | O60814 | USP51 | H2BC12 | 0.896 |
| Q70EL4 | O60814 | USP43 | H2BC12 | 0.860 |
| Q8TEY7 | O60814 | USP33 | H2BC12 | 0.895 |
| Q93009 | O60814 | USP7 | H2BC12 | 0.853 |
| Q96F44 | O60814 | TRIM11 | H2BC12 | 0.858 |
| Q9C040 | O60814 | TRIM2 | H2BC12 | 0.838 |
| Q9H0E7 | O60814 | USP44 | H2BC12 | 0.903 |
| Q9UPT9 | O60814 | USP22 | H2BC12 | 0.912 |
| Q9Y2K6 | O60814 | USP20 | H2BC12 | 0.872 |
| Q9Y4E8 | O60814 | USP15 | H2BC12 | 0.878 |
| Q9Y6I4 | O60814 | USP3 | H2BC12 | 0.855 |
| P12956 | O60814 | XRCC6 | H2BC12 | 0.658 |
| A6NNY8 | Q96A08 | USP27X | H2BC1 | 0.855 |
| O94966 | Q96A08 | USP19 | H2BC1 | 0.860 |
| P35125 | Q96A08 | USP6 | H2BC1 | 0.851 |
| P46736 | Q96A08 | BRCC3 | H2BC1 | 0.825 |
| P51784 | Q96A08 | USP11 | H2BC1 | 0.859 |
| Q13107 | Q96A08 | USP4 | H2BC1 | 0.848 |
| Q5VVJ2 | Q96A08 | MYSM1 | H2BC1 | 0.858 |
| Q70CQ1 | Q96A08 | USP49 | H2BC1 | 0.865 |
| Q70CQ3 | Q96A08 | USP30 | H2BC1 | 0.873 |
| Q70EK9 | Q96A08 | USP51 | H2BC1 | 0.881 |
| Q70EL4 | Q96A08 | USP43 | H2BC1 | 0.839 |
| Q8TEY7 | Q96A08 | USP33 | H2BC1 | 0.880 |
| Q93009 | Q96A08 | USP7 | H2BC1 | 0.829 |
| Q96F44 | Q96A08 | TRIM11 | H2BC1 | 0.840 |
| Q9C040 | Q96A08 | TRIM2 | H2BC1 | 0.821 |
| Q9H0E7 | Q96A08 | USP44 | H2BC1 | 0.889 |
| Q9UPT9 | Q96A08 | USP22 | H2BC1 | 0.900 |
| Q9Y2K6 | Q96A08 | USP20 | H2BC1 | 0.857 |
| Q9Y4E8 | Q96A08 | USP15 | H2BC1 | 0.861 |
| Q9Y6I4 | Q96A08 | USP3 | H2BC1 | 0.831 |
| P12956 | Q96A08 | XRCC6 | H2BC1 | 0.635 |
| A6NNY8 | P33778 | USP27X | H2BC3 | 0.870 |
| O94966 | P33778 | USP19 | H2BC3 | 0.872 |
| P35125 | P33778 | USP6 | H2BC3 | 0.864 |

| | | | | |
|---|---|---|---|---|
| P46736 | P33778 | BRCC3 | H2BC3 | 0.839 |
| P51784 | P33778 | USP11 | H2BC3 | 0.873 |
| Q13107 | P33778 | USP4 | H2BC3 | 0.861 |
| Q5VVJ2 | P33778 | MYSM1 | H2BC3 | 0.870 |
| Q70CQ1 | P33778 | USP49 | H2BC3 | 0.879 |
| Q70CQ3 | P33778 | USP30 | H2BC3 | 0.885 |
| Q70EK9 | P33778 | USP51 | H2BC3 | 0.893 |
| Q70EL4 | P33778 | USP43 | H2BC3 | 0.855 |
| Q8TEY7 | P33778 | USP33 | H2BC3 | 0.892 |
| Q93009 | P33778 | USP7 | H2BC3 | 0.848 |
| Q96F44 | P33778 | TRIM11 | H2BC3 | 0.854 |
| Q9C040 | P33778 | TRIM2 | H2BC3 | 0.835 |
| Q9H0E7 | P33778 | USP44 | H2BC3 | 0.900 |
| Q9H9J4 | P33778 | USP42 | H2BC3 | 0.828 |
| Q9P275 | P33778 | USP36 | H2BC3 | 0.890 |
| Q9UPT9 | P33778 | USP22 | H2BC3 | 0.910 |
| Q9Y2K6 | P33778 | USP20 | H2BC3 | 0.869 |
| Q9Y4E8 | P33778 | USP15 | H2BC3 | 0.875 |
| Q9Y5T5 | P33778 | USP16 | H2BC3 | 0.865 |
| Q9Y6I4 | P33778 | USP3 | H2BC3 | 0.849 |
| P12956 | P33778 | XRCC6 | H2BC3 | 0.652 |
| O75604 | Q99835 | USP2 | SMO | 0.882 |
| Q70EL3 | Q99835 | USP50 | SMO | 0.896 |
| Q70EL4 | Q99835 | USP43 | SMO | 0.857 |
| Q86T82 | Q99835 | USP37 | SMO | 0.860 |
| Q9BXU7 | Q99835 | USP26 | SMO | 0.875 |
| Q9C040 | Q99835 | TRIM2 | SMO | 0.824 |
| Q9HBJ7 | Q99835 | USP29 | SMO | 0.832 |
| Q9UK80 | Q99835 | USP21 | SMO | 0.863 |
| O00487 | P08151 | PSMD14 | GLI1 | 0.920 |
| Q9UK80 | P08151 | USP21 | GLI1 | 0.921 |
| O00487 | P10070 | PSMD14 | GLI2 | 0.957 |
| Q93009 | P10070 | USP7 | GLI2 | 0.946 |
| Q9NQC7 | P10070 | CYLD | GLI2 | 0.976 |
| Q9UK80 | P10070 | USP21 | GLI2 | 0.959 |
| Q96DC9 | Q70CQ4 | OTUB2 | USP31 | 0.503 |
| Q96FW1 | Q70CQ4 | OTUB1 | USP31 | 0.503 |
| Q9Y6I4 | P11802 | USP3 | CDK4 | 0.693 |
| Q9Y6I4 | Q92905 | USP3 | COPS5 | 0.693 |
| Q9Y6I4 | Q7Z6Z7 | USP3 | HUWE1 | 0.691 |
| Q9Y6I4 | P61086 | USP3 | UBE2K | 0.689 |
| Q9Y6I4 | Q9H9V4 | USP3 | RNF122 | 0.687 |
| Q9Y6I4 | Q96I51 | USP3 | RCC1L | 0.683 |

| | | | | |
|---|---|---|---|---|
| Q9Y6I4 | Q9BZK7 | USP3 | TBL1XR1 | 0.682 |
| Q9Y6I4 | P11441 | USP3 | UBL4A | 0.682 |
| Q9Y6I4 | Q2TAZ0 | USP3 | ATG2A | 0.680 |
| Q9Y6I4 | Q6ZSG1 | USP3 | RNF165 | 0.678 |
| Q9Y6I4 | Q9UIF8 | USP3 | BAZ2B | 0.677 |
| Q9Y6I4 | Q9C0D7 | USP3 | ZC3H12C | 0.675 |
| Q9Y6I4 | Q96FS4 | USP3 | SIPA1 | 0.674 |
| Q9Y6I4 | Q96Q15 | USP3 | SMG1 | 0.670 |
| Q9Y6I4 | O95630 | USP3 | STAMBP | 0.670 |
| Q9Y6I4 | Q96J02 | USP3 | ITCH | 0.668 |
| Q9Y6I4 | Q6P444 | USP3 | MTFR2 | 0.668 |
| Q9Y6I4 | Q9GZS3 | USP3 | WDR61 | 0.666 |
| Q9Y6I4 | Q8TEK3 | USP3 | DOT1L | 0.664 |
| Q9Y6I4 | Q13105 | USP3 | ZBTB17 | 0.662 |
| Q9Y6I4 | Q9P2Q2 | USP3 | FRMD4A | 0.662 |
| Q9Y6I4 | Q92995 | USP3 | USP13 | 0.659 |
| Q9Y6I4 | Q9H270 | USP3 | VPS11 | 0.656 |
| Q9Y6I4 | P35227 | USP3 | PCGF2 | 0.655 |
| Q9Y6I4 | Q13347 | USP3 | EIF3I | 0.652 |
| Q9Y6I4 | Q13535 | USP3 | ATR | 0.651 |
| Q9Y6I4 | Q9BUL5 | USP3 | PHF23 | 0.650 |
| Q9Y6I4 | Q6P4R8 | USP3 | NFRKB | 0.650 |
| Q9Y6I4 | Q05086 | USP3 | UBE3A | 0.648 |
| Q9Y6I4 | P53804 | USP3 | TTC3 | 0.646 |
| Q9Y6I4 | Q5HYM0 | USP3 | ZC3H12B | 0.646 |
| Q9Y6I4 | Q9BXM7 | USP3 | PINK1 | 0.645 |
| Q9Y6I4 | Q9UIF9 | USP3 | BAZ2A | 0.643 |
| Q9Y6I4 | Q5VVX9 | USP3 | UBE2U | 0.642 |
| Q9Y6I4 | Q9BT67 | USP3 | NDFIP1 | 0.641 |
| Q9Y6I4 | O60318 | USP3 | MCM3AP | 0.641 |
| Q9Y6I4 | P51668 | USP3 | UBE2D1 | 0.636 |
| Q9Y6I4 | Q9Y4P3 | USP3 | TBL2 | 0.636 |
| Q9Y6I4 | Q14669 | USP3 | TRIP12 | 0.636 |
| Q9Y6I4 | Q9P2H3 | USP3 | IFT80 | 0.634 |
| Q9Y6I4 | O75164 | USP3 | KDM4A | 0.633 |
| Q9Y6I4 | Q9Y4E6 | USP3 | WDR7 | 0.631 |
| Q9Y6I4 | Q15047 | USP3 | SETDB1 | 0.626 |
| Q9Y6I4 | Q15645 | USP3 | TRIP13 | 0.624 |
| Q9Y6I4 | Q9HBJ7 | USP3 | USP29 | 0.624 |
| Q9Y6I4 | P21580 | USP3 | TNFAIP3 | 0.624 |
| Q9Y6I4 | Q8N7F7 | USP3 | UBL4B | 0.622 |
| Q9Y6I4 | Q5U5Q3 | USP3 | MEX3C | 0.619 |
| Q9Y6I4 | O14802 | USP3 | POLR3A | 0.618 |
| Q9Y6I4 | Q92830 | USP3 | KAT2A | 0.617 |
| Q9Y6I4 | O95602 | USP3 | POLR1A | 0.617 |
| Q9Y6I4 | Q9BYB4 | USP3 | GNB1L | 0.612 |
| Q9Y6I4 | Q15819 | USP3 | UBE2V2 | 0.612 |
| Q9Y6I4 | Q9BSC4 | USP3 | NOL10 | 0.612 |

| | | | | |
|---|---|---|---|---|
| Q9Y6I4 | Q96FJ0 | USP3 | STAMBPL1 | 0.611 |
| Q9Y6I4 | P08217 | USP3 | CELA2A | 0.610 |
| Q9Y6I4 | Q15059 | USP3 | BRD3 | 0.606 |
| Q9Y6I4 | Q86WP2 | USP3 | GPBP1 | 0.605 |
| Q9Y6I4 | O95071 | USP3 | UBR5 | 0.603 |
| Q9Y6I4 | P24928 | USP3 | POLR2A | 0.602 |
| Q9Y6I4 | Q14139 | USP3 | UBE4A | 0.597 |
| Q9Y6I4 | Q8IZU2 | USP3 | WDR17 | 0.596 |
| Q9Y6I4 | Q9BRA2 | USP3 | TXNDC17 | 0.595 |
| Q9Y6I4 | Q9UBR5 | USP3 | CKLF | 0.593 |
| Q9Y6I4 | Q8IZW8 | USP3 | TNS4 | 0.592 |
| Q9Y6I4 | Q9BYW2 | USP3 | SETD2 | 0.591 |
| Q9Y6I4 | O15372 | USP3 | EIF3H | 0.590 |
| Q9Y6I4 | O15265 | USP3 | ATXN7 | 0.590 |
| Q9Y6I4 | Q9C0C9 | USP3 | UBE2O | 0.588 |
| Q9Y6I4 | Q92831 | USP3 | KAT2B | 0.587 |
| Q9Y6I4 | Q5T2W1 | USP3 | PDZK1 | 0.586 |
| Q9Y6I4 | Q9ULK2 | USP3 | ATXN7L1 | 0.586 |
| Q9Y6I4 | Q15751 | USP3 | HERC1 | 0.586 |
| Q9Y6I4 | Q5T4S7 | USP3 | UBR4 | 0.585 |
| Q9Y6I4 | P42166 | USP3 | TMPO | 0.585 |
| Q9Y6I4 | P51610 | USP3 | HCFC1 | 0.576 |
| Q9Y6I4 | Q15843 | USP3 | NEDD8 | 0.576 |
| Q9Y6I4 | P09936 | USP3 | UCHL1 | 0.575 |
| Q9Y6I4 | Q5QP82 | USP3 | DCAF10 | 0.574 |
| Q9Y6I4 | Q9Y2I8 | USP3 | WDR37 | 0.573 |
| Q9Y6I4 | Q9C037 | USP3 | TRIM4 | 0.573 |
| Q9Y6I4 | Q9H4Z3 | USP3 | PCIF1 | 0.573 |
| Q9Y6I4 | Q9NRR5 | USP3 | UBQLN4 | 0.571 |
| Q9Y6I4 | Q8TF72 | USP3 | SHROOM3 | 0.571 |
| Q9Y6I4 | Q9BVQ7 | USP3 | SPATA5L1 | 0.570 |
| Q9Y6I4 | O15047 | USP3 | SETD1A | 0.569 |
| Q9Y6I4 | Q9UPU5 | USP3 | USP24 | 0.565 |
| Q9Y6I4 | P61088 | USP3 | UBE2N | 0.564 |
| Q9Y6I4 | P25440 | USP3 | BRD2 | 0.561 |
| Q9Y6I4 | Q9BXF3 | USP3 | CECR2 | 0.560 |
| Q9Y6I4 | P54727 | USP3 | RAD23B | 0.560 |
| Q9Y6I4 | Q6UWZ7 | USP3 | ABRAXAS1 | 0.558 |
| Q9Y6I4 | Q58F21 | USP3 | BRDT | 0.554 |
| Q9Y6I4 | Q86WR6 | USP3 | C17orf64 | 0.554 |
| Q9Y6I4 | Q9H2U1 | USP3 | DHX36 | 0.553 |
| Q9Y6I4 | Q3MJ13 | USP3 | WDR72 | 0.553 |
| Q9Y6I4 | P54725 | USP3 | RAD23A | 0.552 |
| Q9Y6I4 | Q15542 | USP3 | TAF5 | 0.550 |
| Q9Y6I4 | Q15646 | USP3 | OASL | 0.549 |
| Q9Y6I4 | P15374 | USP3 | UCHL3 | 0.548 |
| Q9Y6I4 | O00762 | USP3 | UBE2C | 0.547 |
| Q9Y6I4 | Q14118 | USP3 | DAG1 | 0.546 |

| | | | | |
|---|---|---|---|---|
| Q9Y6I4 | O00303 | USP3 | EIF3F | 0.536 |
| Q9Y6I4 | Q9UNX4 | USP3 | WDR3 | 0.535 |
| Q9Y6I4 | Q5VTH9 | USP3 | DNAI4 | 0.534 |
| Q9Y6I4 | P26196 | USP3 | DDX6 | 0.533 |
| Q9Y6I4 | P35125 | USP3 | USP6 | 0.531 |
| Q9Y6I4 | Q9HAV5 | USP3 | EDA2R | 0.529 |
| Q9Y6I4 | Q9BXB5 | USP3 | OSBPL10 | 0.523 |
| Q9Y6I4 | Q9H0E9 | USP3 | BRD8 | 0.519 |
| Q9Y6I4 | Q5W0U4 | USP3 | BSPRY | 0.518 |
| Q9Y6I4 | P62195 | USP3 | PSMC5 | 0.513 |
| Q9Y6I4 | Q86T82 | USP3 | USP37 | 0.512 |
| Q9Y6I4 | Q9BUR4 | USP3 | WRAP53 | 0.512 |
| Q9Y6I4 | Q8WWQ0 | USP3 | PHIP | 0.510 |
| Q9Y6I4 | Q9Y263 | USP3 | PLAA | 0.510 |
| Q9Y6I4 | Q96MT7 | USP3 | CFAP44 | 0.509 |
| Q9Y6I4 | Q8WVY7 | USP3 | UBLCP1 | 0.506 |
| Q9Y6I4 | P63146 | USP3 | UBE2B | 0.505 |
| Q9Y6I4 | O60331 | USP3 | PIP5K1C | 0.504 |
| Q9Y6I4 | Q8N9V3 | USP3 | WDSUB1 | 0.503 |
| Q9Y6I4 | Q96ST3 | USP3 | SIN3A | 0.501 |
| Q9Y6I4 | O75182 | USP3 | SIN3B | 0.501 |
| Q9Y6I4 | P11802 | USP3 | CDK4 | 0.693 |
| Q9Y6I4 | Q92905 | USP3 | COPS5 | 0.693 |
| Q9Y6I4 | Q7Z6Z7 | USP3 | HUWE1 | 0.691 |
| Q9Y6I4 | P61086 | USP3 | UBE2K | 0.689 |
| Q9Y6I4 | Q9H9V4 | USP3 | RNF122 | 0.687 |
| Q9Y6I4 | Q96I51 | USP3 | RCC1L | 0.683 |
| Q9Y6I4 | Q9BZK7 | USP3 | TBL1XR1 | 0.682 |
| Q9Y6I4 | P11441 | USP3 | UBL4A | 0.682 |
| Q9Y6I4 | Q2TAZ0 | USP3 | ATG2A | 0.680 |
| Q9Y6I4 | Q6ZSG1 | USP3 | RNF165 | 0.678 |
| Q9Y6I4 | Q9UIF8 | USP3 | BAZ2B | 0.677 |
| Q9Y6I4 | Q9C0D7 | USP3 | ZC3H12C | 0.675 |
| Q9Y6I4 | Q96FS4 | USP3 | SIPA1 | 0.674 |
| Q9Y6I4 | Q96Q15 | USP3 | SMG1 | 0.670 |
| Q9Y6I4 | O95630 | USP3 | STAMBP | 0.670 |
| Q9Y6I4 | Q96J02 | USP3 | ITCH | 0.668 |
| Q9Y6I4 | Q6P444 | USP3 | MTFR2 | 0.668 |
| Q9Y6I4 | Q9GZS3 | USP3 | WDR61 | 0.666 |
| Q9Y6I4 | Q8TEK3 | USP3 | DOT1L | 0.664 |
| Q9Y6I4 | Q13105 | USP3 | ZBTB17 | 0.662 |
| Q9Y6I4 | Q9P2Q2 | USP3 | FRMD4A | 0.662 |
| Q9Y6I4 | Q92995 | USP3 | USP13 | 0.659 |
| Q9Y6I4 | Q9H270 | USP3 | VPS11 | 0.656 |
| Q9Y6I4 | P35227 | USP3 | PCGF2 | 0.655 |
| Q9Y6I4 | Q13347 | USP3 | EIF3I | 0.652 |
| Q9Y6I4 | Q13535 | USP3 | ATR | 0.651 |
| Q9Y6I4 | Q9BUL5 | USP3 | PHF23 | 0.650 |

| | | | | |
|---|---|---|---|---|
| Q9Y6I4 | Q6P4R8 | USP3 | NFRKB | 0.650 |
| Q9Y6I4 | Q05086 | USP3 | UBE3A | 0.648 |
| Q9Y6I4 | P53804 | USP3 | TTC3 | 0.646 |
| Q9Y6I4 | Q5HYM0 | USP3 | ZC3H12B | 0.646 |
| Q9Y6I4 | Q9BXM7 | USP3 | PINK1 | 0.645 |
| Q9Y6I4 | Q9UIF9 | USP3 | BAZ2A | 0.643 |
| Q9Y6I4 | Q5VVX9 | USP3 | UBE2U | 0.642 |
| Q9Y6I4 | Q9BT67 | USP3 | NDFIP1 | 0.641 |
| Q9Y6I4 | O60318 | USP3 | MCM3AP | 0.641 |
| Q9Y6I4 | P51668 | USP3 | UBE2D1 | 0.636 |
| Q9Y6I4 | Q9Y4P3 | USP3 | TBL2 | 0.636 |
| Q9Y6I4 | Q14669 | USP3 | TRIP12 | 0.636 |
| Q9Y6I4 | Q9P2H3 | USP3 | IFT80 | 0.634 |
| Q9Y6I4 | O75164 | USP3 | KDM4A | 0.633 |
| Q9Y6I4 | Q9Y4E6 | USP3 | WDR7 | 0.631 |
| Q9Y6I4 | Q15047 | USP3 | SETDB1 | 0.626 |
| Q9Y6I4 | Q15645 | USP3 | TRIP13 | 0.624 |
| Q9Y6I4 | Q9HBJ7 | USP3 | USP29 | 0.624 |
| Q9Y6I4 | P21580 | USP3 | TNFAIP3 | 0.624 |
| Q9Y6I4 | Q8N7F7 | USP3 | UBL4B | 0.622 |
| Q9Y6I4 | Q5U5Q3 | USP3 | MEX3C | 0.619 |
| Q9Y6I4 | O14802 | USP3 | POLR3A | 0.618 |
| Q9Y6I4 | Q92830 | USP3 | KAT2A | 0.617 |
| Q9Y6I4 | O95602 | USP3 | POLR1A | 0.617 |
| Q9Y6I4 | Q9BYB4 | USP3 | GNB1L | 0.612 |
| Q9Y6I4 | Q15819 | USP3 | UBE2V2 | 0.612 |
| Q9Y6I4 | Q9BSC4 | USP3 | NOL10 | 0.612 |
| Q9Y6I4 | Q96FJ0 | USP3 | STAMBPL1 | 0.611 |
| Q9Y6I4 | P08217 | USP3 | CELA2A | 0.610 |
| Q9Y6I4 | Q15059 | USP3 | BRD3 | 0.606 |
| Q9Y6I4 | Q86WP2 | USP3 | GPBP1 | 0.605 |
| Q9Y6I4 | O95071 | USP3 | UBR5 | 0.603 |
| Q9Y6I4 | P24928 | USP3 | POLR2A | 0.602 |
| Q9Y6I4 | Q14139 | USP3 | UBE4A | 0.597 |
| Q9Y6I4 | Q8IZU2 | USP3 | WDR17 | 0.596 |
| Q9Y6I4 | Q9BRA2 | USP3 | TXNDC17 | 0.595 |
| Q9Y6I4 | Q9UBR5 | USP3 | CKLF | 0.593 |
| Q9Y6I4 | Q8IZW8 | USP3 | TNS4 | 0.592 |
| Q9Y6I4 | Q9BYW2 | USP3 | SETD2 | 0.591 |
| Q9Y6I4 | O15372 | USP3 | EIF3H | 0.590 |
| Q9Y6I4 | O15265 | USP3 | ATXN7 | 0.590 |
| Q9Y6I4 | Q9C0C9 | USP3 | UBE2O | 0.588 |
| Q9Y6I4 | Q92831 | USP3 | KAT2B | 0.587 |
| Q9Y6I4 | Q5T2W1 | USP3 | PDZK1 | 0.586 |
| Q9Y6I4 | Q9ULK2 | USP3 | ATXN7L1 | 0.586 |
| Q9Y6I4 | Q15751 | USP3 | HERC1 | 0.586 |
| Q9Y6I4 | Q5T4S7 | USP3 | UBR4 | 0.585 |
| Q9Y6I4 | P42166 | USP3 | TMPO | 0.585 |

| | | | | |
|---|---|---|---|---|
| Q9Y6I4 | P51610 | USP3 | HCFC1 | 0.576 |
| Q9Y6I4 | Q15843 | USP3 | NEDD8 | 0.576 |
| Q9Y6I4 | P09936 | USP3 | UCHL1 | 0.575 |
| Q9Y6I4 | Q5QP82 | USP3 | DCAF10 | 0.574 |
| Q9Y6I4 | Q9Y2I8 | USP3 | WDR37 | 0.573 |
| Q9Y6I4 | Q9C037 | USP3 | TRIM4 | 0.573 |
| Q9Y6I4 | Q9H4Z3 | USP3 | PCIF1 | 0.573 |
| Q9Y6I4 | Q9NRR5 | USP3 | UBQLN4 | 0.571 |
| Q9Y6I4 | Q8TF72 | USP3 | SHROOM3 | 0.571 |
| Q9Y6I4 | Q9BVQ7 | USP3 | SPATA5L1 | 0.570 |
| Q9Y6I4 | O15047 | USP3 | SETD1A | 0.569 |
| Q9Y6I4 | Q9UPU5 | USP3 | USP24 | 0.565 |
| Q9Y6I4 | P61088 | USP3 | UBE2N | 0.564 |
| Q9Y6I4 | P25440 | USP3 | BRD2 | 0.561 |
| Q9Y6I4 | Q9BXF3 | USP3 | CECR2 | 0.560 |
| Q9Y6I4 | P54727 | USP3 | RAD23B | 0.560 |
| Q9Y6I4 | Q6UWZ7 | USP3 | ABRAXAS1 | 0.558 |
| Q9Y6I4 | Q58F21 | USP3 | BRDT | 0.554 |
| Q9Y6I4 | Q86WR6 | USP3 | C17orf64 | 0.554 |
| Q9Y6I4 | Q9H2U1 | USP3 | DHX36 | 0.553 |
| Q9Y6I4 | Q3MJ13 | USP3 | WDR72 | 0.553 |
| Q9Y6I4 | P54725 | USP3 | RAD23A | 0.552 |
| Q9Y6I4 | Q15542 | USP3 | TAF5 | 0.550 |
| Q9Y6I4 | Q15646 | USP3 | OASL | 0.549 |
| Q9Y6I4 | P15374 | USP3 | UCHL3 | 0.548 |
| Q9Y6I4 | O00762 | USP3 | UBE2C | 0.547 |
| Q9Y6I4 | Q14118 | USP3 | DAG1 | 0.546 |
| Q9Y6I4 | O00303 | USP3 | EIF3F | 0.536 |
| Q9Y6I4 | Q9UNX4 | USP3 | WDR3 | 0.535 |
| Q9Y6I4 | Q5VTH9 | USP3 | DNAI4 | 0.534 |
| Q9Y6I4 | P26196 | USP3 | DDX6 | 0.533 |
| Q9Y6I4 | P35125 | USP3 | USP6 | 0.531 |
| Q9Y6I4 | Q9HAV5 | USP3 | EDA2R | 0.529 |
| Q9Y6I4 | Q9BXB5 | USP3 | OSBPL10 | 0.523 |
| Q9Y6I4 | Q9H0E9 | USP3 | BRD8 | 0.519 |
| Q9Y6I4 | Q5W0U4 | USP3 | BSPRY | 0.518 |
| Q9Y6I4 | P62195 | USP3 | PSMC5 | 0.513 |
| Q9Y6I4 | Q86T82 | USP3 | USP37 | 0.512 |
| Q9Y6I4 | Q9BUR4 | USP3 | WRAP53 | 0.512 |
| Q9Y6I4 | Q8WWQ0 | USP3 | PHIP | 0.510 |
| Q9Y6I4 | Q9Y263 | USP3 | PLAA | 0.510 |
| Q9Y6I4 | Q96MT7 | USP3 | CFAP44 | 0.509 |
| Q9Y6I4 | Q8WVY7 | USP3 | UBLCP1 | 0.506 |
| Q9Y6I4 | P63146 | USP3 | UBE2B | 0.505 |
| Q9Y6I4 | O60331 | USP3 | PIP5K1C | 0.504 |
| Q9Y6I4 | Q8N9V3 | USP3 | WDSUB1 | 0.503 |
| Q9Y6I4 | Q96ST3 | USP3 | SIN3A | 0.501 |
| Q9Y6I4 | O75182 | USP3 | SIN3B | 0.501 |

| | | | | |
|---|---|---|---|---|
| P62068 | P55273 | USP46 | CDKN2D | 0.803 |
| P62068 | Q6TDP4 | USP46 | KLHL17 | 0.799 |
| P62068 | Q9NWX5 | USP46 | ASB6 | 0.799 |
| P62068 | Q8WXD9 | USP46 | CASKIN1 | 0.799 |
| P62068 | Q8N8V4 | USP46 | ANKS4B | 0.798 |
| P62068 | O14976 | USP46 | GAK | 0.798 |
| P62068 | P42345 | USP46 | MTOR | 0.795 |
| P62068 | O00221 | USP46 | NFKBIE | 0.795 |
| P62068 | Q8NBL3 | USP46 | TMEM178A | 0.795 |
| P62068 | Q9UNP9 | USP46 | PPIE | 0.794 |
| P62068 | Q8NB46 | USP46 | ANKRD52 | 0.794 |
| P62068 | Q8N9B4 | USP46 | ANKRD42 | 0.793 |
| P62068 | Q969H0 | USP46 | FBXW7 | 0.792 |
| P62068 | P10074 | USP46 | ZBTB48 | 0.791 |
| P62068 | Q15020 | USP46 | SART3 | 0.790 |
| P62068 | Q9UJP4 | USP46 | KLHL21 | 0.788 |
| P62068 | P42772 | USP46 | CDKN2B | 0.787 |
| P62068 | Q96S82 | USP46 | UBL7 | 0.787 |
| P62068 | Q6NXT1 | USP46 | ANKRD54 | 0.786 |
| P62068 | Q8IWQ3 | USP46 | BRSK2 | 0.786 |
| P62068 | O75061 | USP46 | DNAJC6 | 0.784 |
| P62068 | P34903 | USP46 | GABRA3 | 0.784 |
| P62068 | Q9BYH8 | USP46 | NFKBIZ | 0.779 |
| P62068 | O60907 | USP46 | TBL1X | 0.778 |
| P62068 | P49915 | USP46 | GMPS | 0.778 |
| P62068 | Q68DC2 | USP46 | ANKS6 | 0.778 |
| P62068 | Q92905 | USP46 | COPS5 | 0.776 |
| P62068 | Q13489 | USP46 | BIRC3 | 0.776 |
| P62068 | P42771 | USP46 | CDKN2A | 0.776 |
| P62068 | P15621 | USP46 | ZNF44 | 0.775 |
| P62068 | Q15327 | USP46 | ANKRD1 | 0.775 |
| P62068 | Q6NT16 | USP46 | SLC18B1 | 0.773 |
| P62068 | Q96NW4 | USP46 | ANKRD27 | 0.773 |
| P62068 | Q4L235 | USP46 | AASDH | 0.772 |
| P62068 | Q9Y5Z7 | USP46 | HCFC2 | 0.766 |
| P62068 | Q06547 | USP46 | GABPB1 | 0.765 |
| P62068 | Q9UKT8 | USP46 | FBXW2 | 0.764 |
| P62068 | Q8IVT5 | USP46 | KSR1 | 0.763 |
| P62068 | Q9NXR5 | USP46 | ANKRD10 | 0.760 |
| P62068 | Q92527 | USP46 | ANKRD7 | 0.755 |
| P62068 | Q9H6Y7 | USP46 | RNF167 | 0.755 |
| P62068 | O95630 | USP46 | STAMBP | 0.755 |
| P62068 | Q8WVL7 | USP46 | ANKRD49 | 0.755 |
| P62068 | Q9UJX2 | USP46 | CDC23 | 0.754 |
| P62068 | Q8IYU2 | USP46 | HACE1 | 0.751 |
| P62068 | Q8N6D5 | USP46 | ANKRD29 | 0.751 |

| | | | | |
|---|---|---|---|---|
| P62068 | Q9BPW5 | USP46 | RASL11B | 0.748 |
| P62068 | Q8WYP3 | USP46 | RIN2 | 0.748 |
| P62068 | Q9Y2Z0 | USP46 | SUGT1 | 0.748 |
| P62068 | Q9BQI6 | USP46 | SLF1 | 0.747 |
| P62068 | Q9BZK7 | USP46 | TBL1XR1 | 0.746 |
| P62068 | Q96JH7 | USP46 | VCPIP1 | 0.744 |
| P62068 | P11441 | USP46 | UBL4A | 0.742 |
| P62068 | Q6ZVZ8 | USP46 | ASB18 | 0.741 |
| P62068 | P46531 | USP46 | NOTCH1 | 0.739 |
| P62068 | Q92630 | USP46 | DYRK2 | 0.737 |
| P62068 | Q8TBC4 | USP46 | UBA3 | 0.735 |
| P62068 | Q92995 | USP46 | USP13 | 0.735 |
| P62068 | Q9GZV1 | USP46 | ANKRD2 | 0.734 |
| P62068 | Q92625 | USP46 | ANKS1A | 0.732 |
| P62068 | Q6ZVH7 | USP46 | ESPNL | 0.731 |
| P62068 | Q8N3Y1 | USP46 | FBXW8 | 0.730 |
| P62068 | Q86SG2 | USP46 | ANKRD23 | 0.726 |
| P62068 | Q8WYH8 | USP46 | ING5 | 0.725 |
| P62068 | O43791 | USP46 | SPOP | 0.725 |
| P62068 | Q8NG08 | USP46 | HELB | 0.724 |
| P62068 | Q14139 | USP46 | UBE4A | 0.723 |
| P62068 | Q00653 | USP46 | NFKB2 | 0.720 |
| P62068 | Q96J02 | USP46 | ITCH | 0.719 |
| P62068 | Q8WWH4 | USP46 | ASZ1 | 0.718 |
| P62068 | Q9Y252 | USP46 | RNF6 | 0.718 |
| P62068 | Q96PU5 | USP46 | NEDD4L | 0.717 |
| P62068 | Q96Q27 | USP46 | ASB2 | 0.716 |
| P62068 | Q6P6B7 | USP46 | ANKRD16 | 0.716 |
| P62068 | A7E2S9 | USP46 | ANKRD30BL | 0.715 |
| P62068 | Q9P2R3 | USP46 | ANKFY1 | 0.715 |
| P62068 | P82279 | USP46 | CRB1 | 0.713 |
| P62068 | O95271 | USP46 | TNKS | 0.713 |
| P62068 | Q3KP44 | USP46 | ANKRD55 | 0.711 |
| P62068 | Q96I34 | USP46 | PPP1R16A | 0.710 |
| P62068 | Q8TDC3 | USP46 | BRSK1 | 0.709 |
| P62068 | Q6ZW76 | USP46 | ANKS3 | 0.708 |
| P62068 | O15372 | USP46 | EIF3H | 0.707 |
| P62068 | Q96NS5 | USP46 | ASB16 | 0.707 |
| P62068 | Q9Y283 | USP46 | INVS | 0.707 |
| P62068 | Q04721 | USP46 | NOTCH2 | 0.705 |
| P62068 | Q5IJ48 | USP46 | CRB2 | 0.703 |
| Q13107 | P51587 | USP4 | BRCA2 | 0.995 |
| Q13107 | Q9HAW4 | USP4 | CLSPN | 0.993 |
| Q13107 | P12956 | USP4 | XRCC6 | 0.992 |
| Q13107 | P12004 | USP4 | PCNA | 0.980 |
| Q13107 | Q00987 | USP4 | MDM2 | 0.974 |
| Q13107 | O15151 | USP4 | MDM4 | 0.974 |

| | | | | |
|---|---|---|---|---|
| Q13107 | Q49A92 | USP4 | C8orf34 | 0.971 |
| Q13107 | Q99708 | USP4 | RBBP8 | 0.968 |
| Q13107 | O15379 | USP4 | HDAC3 | 0.968 |
| Q13107 | Q13547 | USP4 | HDAC1 | 0.967 |
| Q13107 | Q9BXW9 | USP4 | FANCD2 | 0.959 |
| Q13107 | P01106 | USP4 | MYC | 0.957 |
| Q13107 | P62877 | USP4 | RBX1 | 0.956 |
| Q13107 | Q6PIZ9 | USP4 | TRAT1 | 0.954 |
| Q13107 | Q86UT6 | USP4 | NLRX1 | 0.953 |
| Q13107 | Q12778 | USP4 | FOXO1 | 0.953 |
| Q13107 | Q9BY41 | USP4 | HDAC8 | 0.953 |
| Q13107 | P98177 | USP4 | FOXO4 | 0.946 |
| Q13107 | Q07820 | USP4 | MCL1 | 0.946 |
| Q13107 | P15880 | USP4 | RPS2 | 0.940 |
| Q13107 | Q92844 | USP4 | TANK | 0.940 |
| Q13107 | O43422 | USP4 | THAP12 | 0.940 |
| Q13107 | Q9NS91 | USP4 | RAD18 | 0.939 |
| Q13107 | O43541 | USP4 | SMAD6 | 0.938 |
| Q13107 | Q9P035 | USP4 | HACD3 | 0.937 |
| Q13107 | A6NEH6 | USP4 | TMEM247 | 0.936 |
| Q13107 | Q86WI3 | USP4 | NLRC5 | 0.936 |
| Q13107 | O95140 | USP4 | MFN2 | 0.936 |
| Q13107 | Q8IWA4 | USP4 | MFN1 | 0.936 |
| Q13107 | Q14258 | USP4 | TRIM25 | 0.935 |
| Q13107 | Q5VWC8 | USP4 | HACD4 | 0.935 |
| Q13107 | Q13309 | USP4 | SKP2 | 0.934 |
| Q13107 | Q8IYW5 | USP4 | RNF168 | 0.933 |
| Q13107 | Q9NUU6 | USP4 | OTULINL | 0.929 |
| Q13107 | Q96BN8 | USP4 | OTULIN | 0.929 |
| Q13107 | O15105 | USP4 | SMAD7 | 0.929 |
| Q13107 | P62249 | USP4 | RPS16 | 0.928 |
| Q13107 | P82933 | USP4 | MRPS9 | 0.928 |
| Q13107 | P53597 | USP4 | SUCLG1 | 0.928 |
| Q13107 | O15111 | USP4 | CHUK | 0.927 |
| Q13107 | Q9BYX4 | USP4 | IFIH1 | 0.923 |
| Q13107 | Q5SWX8 | USP4 | ODR4 | 0.923 |
| Q13107 | Q9UQ84 | USP4 | EXO1 | 0.920 |
| Q13107 | O15524 | USP4 | SOCS1 | 0.919 |
| Q13107 | Q92466 | USP4 | DDB2 | 0.917 |
| Q13107 | Q96EP0 | USP4 | RNF31 | 0.914 |
| Q13107 | P51786 | USP4 | ZNF157 | 0.911 |
| Q13107 | Q7RTR2 | USP4 | NLRC3 | 0.911 |
| Q13107 | P51617 | USP4 | IRAK1 | 0.908 |
| Q13107 | Q9HB75 | USP4 | PIDD1 | 0.908 |
| Q13107 | Q14653 | USP4 | IRF3 | 0.907 |
| Q13107 | Q6Y1H2 | USP4 | HACD2 | 0.906 |
| Q13107 | O60285 | USP4 | NUAK1 | 0.905 |
| Q13107 | O14757 | USP4 | CHEK1 | 0.902 |

| | | | | |
|---|---|---|---|---|
| Q13107 | O14920 | USP4 | IKBKB | 0.898 |
| Q13107 | Q13114 | USP4 | TRAF3 | 0.894 |
| Q13107 | P48730 | USP4 | CSNK1D | 0.894 |
| Q13107 | B0YJ81 | USP4 | HACD1 | 0.894 |
| Q13107 | P0C1H6 | USP4 | H2BW2 | 0.893 |
| Q13107 | Q93052 | USP4 | LPP | 0.893 |
| Q13107 | Q9UBS8 | USP4 | RNF14 | 0.893 |
| Q13107 | Q15831 | USP4 | STK11 | 0.891 |
| Q13107 | Q9UBF6 | USP4 | RNF7 | 0.890 |
| Q13107 | O00444 | USP4 | PLK4 | 0.890 |
| Q13107 | Q9C0F3 | USP4 | ZNF436 | 0.889 |
| Q13107 | Q99728 | USP4 | BARD1 | 0.888 |
| Q13107 | Q96EP1 | USP4 | CHFR | 0.886 |
| Q13107 | Q03519 | USP4 | TAP2 | 0.883 |
| Q13107 | Q7Z2G1 | USP4 | H2BW1 | 0.882 |
| Q13107 | Q6U7Q0 | USP4 | ZNF322 | 0.881 |
| Q13107 | Q9UEG4 | USP4 | ZNF629 | 0.878 |
| Q13107 | Q9NQB0 | USP4 | TCF7L2 | 0.875 |
| Q13107 | P58753 | USP4 | TIRAP | 0.874 |
| Q13107 | Q9Y6K9 | USP4 | IKBKG | 0.874 |
| Q13107 | Q96CV9 | USP4 | OPTN | 0.874 |
| Q13107 | P20749 | USP4 | BCL3 | 0.874 |
| Q13107 | Q8NH60 | USP4 | OR52J3 | 0.873 |
| Q13107 | Q3MII6 | USP4 | TBC1D25 | 0.873 |
| Q13107 | Q14457 | USP4 | BECN1 | 0.872 |
| Q13107 | O00635 | USP4 | TRIM38 | 0.872 |
| Q13107 | Q15942 | USP4 | ZYX | 0.871 |
| Q13107 | O14896 | USP4 | IRF6 | 0.868 |
| Q13107 | Q13976 | USP4 | PRKG1 | 0.867 |
| Q13107 | Q9P243 | USP4 | ZFAT | 0.867 |
| Q13107 | P36894 | USP4 | BMPR1A | 0.867 |
| Q13107 | Q13043 | USP4 | STK4 | 0.867 |
| Q13107 | Q9Y577 | USP4 | TRIM17 | 0.866 |
| Q13107 | Q99879 | USP4 | H2BC14 | 0.866 |
| Q13107 | Q99836 | USP4 | MYD88 | 0.865 |
| Q13107 | Q2M1V0 | USP4 | ISX | 0.865 |
| Q13107 | Q9H4P4 | USP4 | RNF41 | 0.865 |
| Q13107 | O60814 | USP4 | H2BC12 | 0.864 |
| Q13107 | Q7Z3U7 | USP4 | MON2 | 0.863 |
| Q13107 | Q99880 | USP4 | H2BC13 | 0.863 |
| Q13107 | Q92547 | USP4 | TOPBP1 | 0.863 |
| Q13107 | Q8NGJ4 | USP4 | OR52E2 | 0.863 |
| Q13107 | Q6ZWH5 | USP4 | NEK10 | 0.862 |
| Q13107 | P29074 | USP4 | PTPN4 | 0.862 |
| Q13107 | Q9NWZ3 | USP4 | IRAK4 | 0.861 |
| Q13107 | Q6ZVD8 | USP4 | PHLPP2 | 0.861 |
| Q13107 | P33778 | USP4 | H2BC3 | 0.861 |
| Q13107 | Q13315 | USP4 | ATM | 0.861 |

| | | | | |
|---|---|---|---|---|
| Q13107 | Q9Y468 | USP4 | L3MBTL1 | 0.860 |
| Q13107 | P98170 | USP4 | XIAP | 0.860 |
| Q13107 | P37275 | USP4 | ZEB1 | 0.859 |
| Q13107 | Q9NR21 | USP4 | PARP11 | 0.859 |
| Q13107 | Q9H4I2 | USP4 | ZHX3 | 0.857 |
| Q13107 | Q96CX3 | USP4 | ZNF501 | 0.857 |
| Q13107 | Q9C035 | USP4 | TRIM5 | 0.857 |
| Q13107 | Q8IXI2 | USP4 | RHOT1 | 0.857 |
| Q13107 | P58876 | USP4 | H2BC5 | 0.856 |
| Q13107 | Q99877 | USP4 | H2BC15 | 0.856 |
| Q13107 | O75165 | USP4 | DNAJC13 | 0.854 |
| Q13107 | O00463 | USP4 | TRAF5 | 0.853 |
| Q13107 | Q5QNW6 | USP4 | H2BC18 | 0.853 |
| Q13107 | O43187 | USP4 | IRAK2 | 0.853 |
| Q13107 | P62807 | USP4 | H2BC4 | 0.853 |
| Q13107 | Q93079 | USP4 | H2BC9 | 0.852 |
| Q13107 | Q8NI38 | USP4 | NFKBID | 0.852 |
| Q13107 | Q9UHD2 | USP4 | TBK1 | 0.852 |
| Q13107 | P06899 | USP4 | H2BC11 | 0.851 |
| Q13107 | Q9H6E5 | USP4 | TUT1 | 0.851 |
| Q13107 | Q96C10 | USP4 | DHX58 | 0.850 |
| Q13107 | P57053 | USP4 | H2BS1 | 0.849 |
| Q13107 | Q16690 | USP4 | DUSP5 | 0.849 |
| Q13107 | Q96RU7 | USP4 | TRIB3 | 0.848 |
| Q13107 | Q96A08 | USP4 | H2BC1 | 0.848 |
| Q13107 | Q5QJU3 | USP4 | ACER2 | 0.847 |
| Q13107 | Q5MCW4 | USP4 | ZNF569 | 0.846 |
| Q13107 | Q9BXS5 | USP4 | AP1M1 | 0.845 |
| Q13107 | Q8N257 | USP4 | H2BU1 | 0.845 |
| Q13107 | Q15365 | USP4 | PCBP1 | 0.844 |
| Q13107 | O60346 | USP4 | PHLPP1 | 0.844 |
| Q13107 | P21453 | USP4 | S1PR1 | 0.842 |
| Q13107 | Q96EB6 | USP4 | SIRT1 | 0.840 |
| Q13107 | Q96AQ6 | USP4 | PBXIP1 | 0.840 |
| Q13107 | P84022 | USP4 | SMAD3 | 0.840 |
| Q13107 | Q9HCE7 | USP4 | SMURF1 | 0.839 |
| Q13107 | Q16778 | USP4 | H2BC21 | 0.839 |
| Q13107 | P23527 | USP4 | H2BC17 | 0.839 |
| Q13107 | O95405 | USP4 | ZFYVE9 | 0.838 |
| Q13107 | P31751 | USP4 | AKT2 | 0.838 |
| Q13107 | Q9HCP0 | USP4 | CSNK1G1 | 0.836 |
| Q13107 | Q12974 | USP4 | PTP4A2 | 0.836 |
| Q13107 | P50570 | USP4 | DNM2 | 0.835 |
| Q13107 | Q9UKV5 | USP4 | AMFR | 0.835 |
| Q13107 | Q96M94 | USP4 | KLHL15 | 0.834 |
| Q13107 | Q9NTX7 | USP4 | RNF146 | 0.833 |
| Q13107 | Q9HAU4 | USP4 | SMURF2 | 0.833 |
| Q13107 | Q9P289 | USP4 | STK26 | 0.833 |

| | | | | |
|---|---|---|---|---|
| Q13107 | Q15796 | USP4 | SMAD2 | 0.833 |
| Q13107 | O96017 | USP4 | CHEK2 | 0.827 |
| Q13107 | Q9BQI6 | USP4 | SLF1 | 0.825 |
| Q13107 | P38398 | USP4 | BRCA1 | 0.824 |
| Q13107 | O43432 | USP4 | EIF4G3 | 0.824 |
| Q13107 | O60858 | USP4 | TRIM13 | 0.822 |
| Q13107 | Q9NZB8 | USP4 | MOCS1 | 0.822 |
| Q13107 | Q504Q3 | USP4 | PAN2 | 0.822 |
| Q13107 | Q9H2X6 | USP4 | HIPK2 | 0.822 |
| Q13107 | O60260 | USP4 | PRKN | 0.820 |
| Q13107 | Q8TDR2 | USP4 | STK35 | 0.820 |
| Q13107 | Q13490 | USP4 | BIRC2 | 0.820 |
| Q13107 | P31749 | USP4 | AKT1 | 0.819 |
| Q13107 | O14976 | USP4 | GAK | 0.817 |
| Q13107 | P78314 | USP4 | SH3BP2 | 0.814 |
| Q13107 | Q96EQ8 | USP4 | RNF125 | 0.814 |
| Q13107 | O94972 | USP4 | TRIM37 | 0.813 |
| Q13107 | Q9H2U2 | USP4 | PPA2 | 0.812 |
| Q13107 | Q92973 | USP4 | TNPO1 | 0.810 |
| Q13107 | P51965 | USP4 | UBE2E1 | 0.809 |
| Q13107 | Q9HCS5 | USP4 | EPB41L4A | 0.807 |
| Q13107 | P63010 | USP4 | AP2B1 | 0.807 |
| Q13107 | Q15750 | USP4 | TAB1 | 0.804 |
| Q13107 | Q7LFL8 | USP4 | CXXC5 | 0.803 |
| Q13107 | Q8N1C3 | USP4 | GABRG1 | 0.803 |
| Q13107 | Q6ZMK1 | USP4 | CYHR1 | 0.803 |
| Q13107 | O15033 | USP4 | AREL1 | 0.802 |
| Q13107 | Q15628 | USP4 | TRADD | 0.800 |
| Q13107 | Q6ZNA4 | USP4 | RNF111 | 0.800 |
| Q13107 | Q8N5S9 | USP4 | CAMKK1 | 0.800 |
| Q13107 | P16104 | USP4 | H2AX | 0.799 |
| Q13107 | Q9ULT6 | USP4 | ZNRF3 | 0.799 |
| Q13107 | P07602 | USP4 | PSAP | 0.796 |
| Q13107 | P42566 | USP4 | EPS15 | 0.796 |
| Q13107 | Q96S59 | USP4 | RANBP9 | 0.794 |
| Q13107 | O60313 | USP4 | OPA1 | 0.793 |
| Q13107 | Q99942 | USP4 | RNF5 | 0.790 |
| Q13107 | Q15020 | USP4 | SART3 | 0.789 |
| Q13107 | P46937 | USP4 | YAP1 | 0.789 |
| Q13107 | A6ND36 | USP4 | FAM83G | 0.788 |
| Q13107 | Q92556 | USP4 | ELMO1 | 0.786 |
| Q13107 | Q96RR4 | USP4 | CAMKK2 | 0.782 |
| Q13107 | Q7Z569 | USP4 | BRAP | 0.780 |
| Q13107 | Q9Y2U9 | USP4 | KLHDC2 | 0.779 |
| Q13107 | Q12888 | USP4 | TP53BP1 | 0.779 |
| Q13107 | O14787 | USP4 | TNPO2 | 0.779 |
| Q13107 | Q9Y2E6 | USP4 | DTX4 | 0.777 |
| Q13107 | O76064 | USP4 | RNF8 | 0.775 |

| | | | | |
|---|---|---|---|---|
| Q13107 | Q16777 | USP4 | H2AC20 | 0.775 |
| Q13107 | Q53EL6 | USP4 | PDCD4 | 0.774 |
| Q13107 | Q15654 | USP4 | TRIP6 | 0.773 |
| Q13107 | Q96JH7 | USP4 | VCPIP1 | 0.772 |
| Q13107 | Q5D1E8 | USP4 | ZC3H12A | 0.771 |
| Q13107 | Q8NA03 | USP4 | FSIP1 | 0.767 |
| Q13107 | Q9UIF8 | USP4 | BAZ2B | 0.767 |
| Q13107 | Q8N9H8 | USP4 | EXD3 | 0.765 |
| Q13107 | Q7L2J0 | USP4 | MEPCE | 0.765 |
| Q13107 | O15013 | USP4 | ARHGEF10 | 0.763 |
| Q13107 | Q9UK12 | USP4 | ZNF222 | 0.762 |
| Q13107 | Q969K4 | USP4 | ABTB1 | 0.760 |
| Q13107 | Q13158 | USP4 | FADD | 0.759 |
| Q13107 | Q86VP6 | USP4 | CAND1 | 0.759 |
| Q13107 | O43255 | USP4 | SIAH2 | 0.757 |
| Q13107 | Q9NVW2 | USP4 | RLIM | 0.757 |
| Q13107 | Q13489 | USP4 | BIRC3 | 0.756 |
| Q13107 | Q9UMS4 | USP4 | PRPF19 | 0.756 |
| Q13107 | Q8TAF7 | USP4 | ZNF461 | 0.753 |
| Q13107 | P46776 | USP4 | RPL27A | 0.752 |
| Q13107 | Q96JY6 | USP4 | PDLIM2 | 0.752 |
| Q13107 | Q9HCE6 | USP4 | ARHGEF10L | 0.750 |
| Q13107 | Q9NVH0 | USP4 | EXD2 | 0.748 |
| Q13107 | Q9UEW8 | USP4 | STK39 | 0.746 |
| Q13107 | Q9UIF9 | USP4 | BAZ2A | 0.746 |
| Q13107 | Q9NYT6 | USP4 | ZNF226 | 0.744 |
| Q13107 | Q9ULT8 | USP4 | HECTD1 | 0.744 |
| Q13107 | Q9Y4A0 | USP4 | JRKL | 0.743 |
| Q13107 | Q9BYH1 | USP4 | SEZ6L | 0.743 |
| Q13107 | Q99729 | USP4 | HNRNPAB | 0.739 |
| Q13107 | Q6PIW4 | USP4 | FIGNL1 | 0.738 |
| Q13107 | P61421 | USP4 | ATP6V0D1 | 0.737 |
| Q13107 | Q96PU5 | USP4 | NEDD4L | 0.736 |
| Q13107 | P51530 | USP4 | DNA2 | 0.736 |
| Q13107 | Q7L9B9 | USP4 | EEPD1 | 0.736 |
| Q13107 | Q14103 | USP4 | HNRNPD | 0.733 |
| Q13107 | Q9H6R0 | USP4 | DHX33 | 0.730 |
| Q13107 | Q96L34 | USP4 | MARK4 | 0.729 |
| Q13107 | P45983 | USP4 | MAPK8 | 0.729 |
| Q13107 | Q2NKQ1 | USP4 | SGSM1 | 0.728 |
| Q13107 | Q96J02 | USP4 | ITCH | 0.728 |
| Q13107 | Q9Y2Z2 | USP4 | MTO1 | 0.725 |
| Q13107 | Q14669 | USP4 | TRIP12 | 0.720 |
| Q13107 | Q8IW03 | USP4 | SIAH3 | 0.719 |
| Q13107 | Q9H4L5 | USP4 | OSBPL3 | 0.719 |
| Q13107 | Q7Z6J4 | USP4 | FGD2 | 0.718 |
| Q13107 | O43151 | USP4 | TET3 | 0.717 |
| Q13107 | Q05086 | USP4 | UBE3A | 0.717 |

| | | | | |
|---|---|---|---|---|
| Q13107 | Q6ZS81 | USP4 | WDFY4 | 0.715 |
| Q13107 | Q9UBT2 | USP4 | UBA2 | 0.714 |
| Q13107 | A6NKF1 | USP4 | SAC3D1 | 0.713 |
| Q13107 | Q8NFZ0 | USP4 | FBH1 | 0.711 |
| Q13107 | Q9UKD2 | USP4 | MRTO4 | 0.711 |
| Q13107 | Q92783 | USP4 | STAM | 0.710 |
| Q13107 | Q8N6T7 | USP4 | SIRT6 | 0.709 |
| Q13107 | P46934 | USP4 | NEDD4 | 0.708 |
| Q13107 | P15621 | USP4 | ZNF44 | 0.708 |
| Q13107 | Q9BYW2 | USP4 | SETD2 | 0.708 |
| Q13107 | Q14160 | USP4 | SCRIB | 0.707 |
| Q13107 | O43889 | USP4 | CREB3 | 0.706 |
| Q13107 | Q5XPI4 | USP4 | RNF123 | 0.706 |
| Q13107 | Q13751 | USP4 | LAMB3 | 0.705 |
| Q13107 | Q9UKF6 | USP4 | CPSF3 | 0.702 |
| Q5VVQ6 | Q13107 | YOD1 | USP4 | 0.744 |
| Q13107 | O14979 | USP4 | HNRNPDL | 0.701 |
| Q13107 | O60318 | USP4 | MCM3AP | 0.701 |
| Q13107 | Q92995 | USP4 | USP13 | 0.701 |
| Q13107 | Q9H6B1 | USP4 | ZNF385D | 0.701 |
| Q13107 | A5PLN7 | USP4 | FAM149A | 0.694 |
| Q13107 | Q8IUD6 | USP4 | RNF135 | 0.694 |
| Q13107 | P11717 | USP4 | IGF2R | 0.693 |
| Q13107 | O95222 | USP4 | OR6A2 | 0.693 |
| Q13107 | P11441 | USP4 | UBL4A | 0.691 |
| Q13107 | O00206 | USP4 | TLR4 | 0.691 |
| Q13107 | Q96S82 | USP4 | UBL7 | 0.690 |
| Q13107 | Q9Y385 | USP4 | UBE2J1 | 0.687 |
| Q13107 | P01375 | USP4 | TNF | 0.686 |
| Q13107 | Q9NZI7 | USP4 | UBP1 | 0.686 |
| Q13107 | Q7Z2W9 | USP4 | MRPL21 | 0.684 |
| Q13107 | Q6P4R8 | USP4 | NFRKB | 0.684 |
| Q13107 | P21580 | USP4 | TNFAIP3 | 0.684 |
| Q13107 | Q9NXA8 | USP4 | SIRT5 | 0.679 |
| Q13107 | Q9HBJ7 | USP4 | USP29 | 0.678 |
| Q13107 | O60522 | USP4 | TDRD6 | 0.676 |
| Q13107 | A1L390 | USP4 | PLEKHG3 | 0.676 |
| Q13107 | O95071 | USP4 | UBR5 | 0.675 |
| Q13107 | Q8TEK3 | USP4 | DOT1L | 0.672 |
| Q13107 | Q9H0M0 | USP4 | WWP1 | 0.668 |
| Q13107 | O15230 | USP4 | LAMA5 | 0.668 |
| Q13107 | Q13588 | USP4 | GRAP | 0.666 |
| Q13107 | Q96Q89 | USP4 | KIF20B | 0.665 |
| Q13107 | A6H8M9 | USP4 | CDHR4 | 0.664 |
| Q13107 | Q92905 | USP4 | COPS5 | 0.664 |
| Q13107 | O95602 | USP4 | POLR1A | 0.660 |
| Q13107 | O14802 | USP4 | POLR3A | 0.658 |
| Q13107 | Q99741 | USP4 | CDC6 | 0.658 |

| | | | | |
|---|---|---|---|---|
| Q13107 | Q13415 | USP4 | ORC1 | 0.658 |
| Q13107 | Q02809 | USP4 | PLOD1 | 0.658 |
| Q13107 | O15085 | USP4 | ARHGEF11 | 0.654 |
| Q13107 | Q8WWF5 | USP4 | ZNRF4 | 0.653 |
| Q13107 | P43897 | USP4 | TSFM | 0.652 |
| Q13107 | O95630 | USP4 | STAMBP | 0.651 |
| Q13107 | Q8WWT9 | USP4 | SLC13A3 | 0.649 |
| Q13107 | Q8IUQ4 | USP4 | SIAH1 | 0.648 |
| Q13107 | Q9UPU5 | USP4 | USP24 | 0.647 |
| Q13107 | O15047 | USP4 | SETD1A | 0.646 |
| Q13107 | Q5T4S7 | USP4 | UBR4 | 0.646 |
| Q13107 | P09001 | USP4 | MRPL3 | 0.645 |
| Q13107 | O75717 | USP4 | WDHD1 | 0.644 |
| Q13107 | P27348 | USP4 | YWHAQ | 0.643 |
| Q13107 | P24928 | USP4 | POLR2A | 0.643 |
| Q13107 | O75688 | USP4 | PPM1B | 0.643 |
| Q13107 | Q96QB1 | USP4 | DLC1 | 0.642 |
| Q13107 | Q8NDG6 | USP4 | TDRD9 | 0.640 |
| Q13107 | Q9NXJ5 | USP4 | PGPEP1 | 0.639 |
| Q13107 | P49792 | USP4 | RANBP2 | 0.639 |
| Q13107 | P05388 | USP4 | RPLP0 | 0.635 |
| Q13107 | P05026 | USP4 | ATP1B1 | 0.630 |
| Q13107 | Q92796 | USP4 | DLG3 | 0.624 |
| Q13107 | Q15878 | USP4 | CACNA1E | 0.622 |
| Q13107 | Q8N7F7 | USP4 | UBL4B | 0.622 |
| Q13107 | Q9C0C9 | USP4 | UBE2O | 0.619 |
| Q13107 | Q8IZQ1 | USP4 | WDFY3 | 0.616 |
| Q13107 | P40763 | USP4 | STAT3 | 0.615 |
| Q13107 | Q96A65 | USP4 | EXOC4 | 0.615 |
| Q13107 | P35125 | USP4 | USP6 | 0.614 |
| Q13107 | Q9BZL1 | USP4 | UBL5 | 0.612 |
| Q13107 | Q5VVX9 | USP4 | UBE2U | 0.611 |
| Q13107 | Q70EL1 | USP4 | USP54 | 0.609 |
| Q13107 | P23588 | USP4 | EIF4B | 0.609 |
| Q13107 | O95235 | USP4 | KIF20A | 0.608 |
| Q13107 | Q9UPN9 | USP4 | TRIM33 | 0.607 |
| Q13107 | Q5U5Q3 | USP4 | MEX3C | 0.601 |
| Q13107 | Q9P0M9 | USP4 | MRPL27 | 0.599 |
| Q13107 | Q9H2U1 | USP4 | DHX36 | 0.598 |
| Q13107 | Q9UGI0 | USP4 | ZRANB1 | 0.598 |
| Q13107 | O60508 | USP4 | CDC40 | 0.598 |
| Q13107 | O43447 | USP4 | PPIH | 0.594 |
| Q13107 | O43791 | USP4 | SPOP | 0.593 |
| Q13107 | Q9UNE7 | USP4 | STUB1 | 0.592 |
| Q13107 | Q9UKZ4 | USP4 | TENM1 | 0.591 |
| Q13107 | Q9ULI0 | USP4 | ATAD2B | 0.590 |
| Q13107 | Q86TM6 | USP4 | SYVN1 | 0.590 |
| Q13107 | P51991 | USP4 | HNRNPA3 | 0.590 |

| | | | | |
|---|---|---|---|---|
| Q13107 | Q15018 | USP4 | ABRAXAS2 | 0.587 |
| Q13107 | Q96FJ0 | USP4 | STAMBPL1 | 0.586 |
| Q13107 | Q9BVQ7 | USP4 | SPATA5L1 | 0.585 |
| Q13107 | Q9BQ95 | USP4 | ECSIT | 0.583 |
| Q13107 | Q6P158 | USP4 | DHX57 | 0.583 |
| Q13107 | Q13191 | USP4 | CBLB | 0.583 |
| Q13107 | O00303 | USP4 | EIF3F | 0.583 |
| Q13107 | Q9Y263 | USP4 | PLAA | 0.583 |
| Q13107 | Q86T82 | USP4 | USP37 | 0.580 |
| Q13107 | Q01804 | USP4 | OTUD4 | 0.579 |
| Q13107 | Q5T5U3 | USP4 | ARHGAP21 | 0.579 |
| Q13107 | A0AVT1 | USP4 | UBA6 | 0.578 |
| Q13107 | Q92820 | USP4 | GGH | 0.578 |
| Q13107 | P07910 | USP4 | HNRNPC | 0.577 |
| Q13107 | Q9UBP0 | USP4 | SPAST | 0.576 |
| Q13107 | Q86Y37 | USP4 | CACUL1 | 0.575 |
| Q13107 | Q15646 | USP4 | OASL | 0.575 |
| Q13107 | Q06710 | USP4 | PAX8 | 0.572 |
| Q13107 | Q9BUR4 | USP4 | WRAP53 | 0.565 |
| Q13107 | P51668 | USP4 | UBE2D1 | 0.557 |
| Q13107 | Q9UNX4 | USP4 | WDR3 | 0.557 |
| Q13107 | Q15843 | USP4 | NEDD8 | 0.554 |
| Q13107 | Q8WVY7 | USP4 | UBLCP1 | 0.551 |
| Q13107 | Q9BYD1 | USP4 | MRPL13 | 0.550 |
| Q13107 | Q9Y3Q3 | USP4 | TMED3 | 0.548 |
| Q13107 | P82675 | USP4 | MRPS5 | 0.547 |
| Q13107 | O43815 | USP4 | STRN | 0.546 |
| Q13107 | Q00341 | USP4 | HDLBP | 0.545 |
| Q13107 | Q7KZ85 | USP4 | SUPT6H | 0.544 |
| Q13107 | O15372 | USP4 | EIF3H | 0.544 |
| Q13107 | Q99961 | USP4 | SH3GL1 | 0.543 |
| Q13107 | P45974 | USP4 | USP5 | 0.541 |
| Q13107 | Q8N668 | USP4 | COMMD1 | 0.541 |
| Q13107 | P62195 | USP4 | PSMC5 | 0.534 |
| Q13107 | A6NKT7 | USP4 | RGPD3 | 0.533 |
| Q13107 | Q9UPS6 | USP4 | SETD1B | 0.532 |
| Q13107 | P54727 | USP4 | RAD23B | 0.532 |
| Q13107 | Q6P589 | USP4 | TNFAIP8L2 | 0.531 |
| Q13107 | Q8IZ57 | USP4 | NRSN1 | 0.531 |
| Q13107 | Q9C037 | USP4 | TRIM4 | 0.530 |
| Q13107 | P09651 | USP4 | HNRNPA1 | 0.528 |
| Q13107 | P09936 | USP4 | UCHL1 | 0.528 |
| Q13107 | P39023 | USP4 | RPL3 | 0.525 |
| Q13107 | Q92901 | USP4 | RPL3L | 0.525 |
| Q13107 | Q7L2E3 | USP4 | DHX30 | 0.525 |
| Q13107 | Q9BV94 | USP4 | EDEM2 | 0.524 |
| Q13107 | Q8NFA0 | USP4 | USP32 | 0.524 |
| Q13107 | Q6ZRS2 | USP4 | SRCAP | 0.524 |

| | | | | |
|---|---|---|---|---|
| Q13107 | Q32P51 | USP4 | HNRNPA1L2 | 0.523 |
| Q13107 | Q9BZR9 | USP4 | TRIM8 | 0.523 |
| Q13107 | P49406 | USP4 | MRPL19 | 0.522 |
| Q13107 | Q15652 | USP4 | JMJD1C | 0.522 |
| Q13107 | Q9NYJ8 | USP4 | TAB2 | 0.522 |
| Q13107 | Q8N5C8 | USP4 | TAB3 | 0.522 |
| Q13107 | Q9Y3T6 | USP4 | R3HCC1 | 0.519 |
| Q13107 | Q9BQY4 | USP4 | RHOXF2 | 0.517 |
| Q13107 | O60812 | USP4 | HNRNPCL1 | 0.517 |
| Q13107 | Q9Y3C5 | USP4 | RNF11 | 0.516 |
| Q13107 | Q8TAF3 | USP4 | WDR48 | 0.515 |
| Q13107 | Q70CQ3 | USP4 | USP30 | 0.513 |
| Q13107 | Q9BUZ4 | USP4 | TRAF4 | 0.513 |
| Q13107 | P51124 | USP4 | GZMM | 0.513 |
| Q13107 | Q14139 | USP4 | UBE4A | 0.512 |
| Q13107 | P54709 | USP4 | ATP1B3 | 0.508 |
| Q13107 | Q8IX18 | USP4 | DHX40 | 0.508 |
| Q13107 | Q13151 | USP4 | HNRNPA0 | 0.506 |
| Q13107 | Q13404 | USP4 | UBE2V1 | 0.504 |
| Q13107 | Q09472 | USP4 | EP300 | 0.504 |
| Q13107 | Q9Y3B3 | USP4 | TMED7 | 0.501 |
| Q13107 | P15374 | USP4 | UCHL3 | 0.500 |
| Q14694 | Q13107 | USP10 | USP4 | 0.508 |
| Q504Q3 | Q13107 | PAN2 | USP4 | 0.523 |
| Q70EL4 | Q13107 | USP43 | USP4 | 0.506 |
| Q96DC9 | Q13107 | OTUB2 | USP4 | 0.537 |
| Q96FW1 | Q13107 | OTUB1 | USP4 | 0.537 |
| Q9H3M9 | Q13107 | ATXN3L | USP4 | 0.513 |
| Q9UMW8 | Q13107 | USP18 | USP4 | 0.508 |
| Q9UPT9 | Q13107 | USP22 | USP4 | 0.513 |
| Q5VVQ6 | Q9NVE5 | YOD1 | USP40 | 0.617 |
| Q9UK80 | Q9HAW4 | USP21 | CLSPN | 0.992 |
| Q9UK80 | Q9H040 | USP21 | SPRTN | 0.975 |
| Q9UK80 | P04637 | USP21 | TP53 | 0.970 |
| Q9UK80 | Q00987 | USP21 | MDM2 | 0.966 |
| Q9UK80 | Q96RL1 | USP21 | UIMC1 | 0.963 |
| Q9UK80 | Q13547 | USP21 | HDAC1 | 0.962 |
| Q9UK80 | P10070 | USP21 | GLI2 | 0.959 |
| Q9UK80 | Q92769 | USP21 | HDAC2 | 0.958 |
| Q9UK80 | P05231 | USP21 | IL6 | 0.955 |
| Q9UK80 | Q01831 | USP21 | XPC | 0.952 |
| Q9UK80 | Q99708 | USP21 | RBBP8 | 0.947 |
| Q9UK80 | P27816 | USP21 | MAP4 | 0.935 |
| Q9UK80 | Q969S8 | USP21 | HDAC10 | 0.932 |
| Q9UK80 | Q9Y4K3 | USP21 | TRAF6 | 0.931 |
| Q9UK80 | Q04206 | USP21 | RELA | 0.924 |

| | | | | |
|---|---|---|---|---|
| Q9UK80 | P08151 | USP21 | GLI1 | 0.921 |
| Q9UK80 | P56524 | USP21 | HDAC4 | 0.920 |
| Q9UK80 | Q96BN8 | USP21 | OTULIN | 0.917 |
| Q9UK80 | Q9NUU6 | USP21 | OTULINL | 0.917 |
| Q9UK80 | O60353 | USP21 | FZD6 | 0.913 |
| Q9UK80 | Q8IYW5 | USP21 | RNF168 | 0.913 |
| Q9UK80 | Q9H7Z6 | USP21 | KAT8 | 0.905 |
| Q9UK80 | Q8N531 | USP21 | FBXL6 | 0.904 |
| Q9UK80 | O00327 | USP21 | ARNTL | 0.902 |
| Q9UK80 | P42356 | USP21 | PI4KA | 0.894 |
| Q9UK80 | Q92889 | USP21 | ERCC4 | 0.893 |
| Q9UK80 | Q99496 | USP21 | RNF2 | 0.890 |
| Q9UK80 | Q7Z7B1 | USP21 | PIGW | 0.888 |
| Q9UK80 | O15111 | USP21 | CHUK | 0.880 |
| Q9UK80 | Q9BYX4 | USP21 | IFIH1 | 0.879 |
| Q9UK80 | Q14332 | USP21 | FZD2 | 0.878 |
| Q9UK80 | Q5W0Q7 | USP21 | USPL1 | 0.874 |
| Q9UK80 | Q9NPG1 | USP21 | FZD3 | 0.874 |
| Q9UK80 | Q9BY79 | USP21 | MFRP | 0.872 |
| Q9UK80 | O15055 | USP21 | PER2 | 0.872 |
| Q9UK80 | O15534 | USP21 | PER1 | 0.870 |
| Q9UK80 | P25963 | USP21 | NFKBIA | 0.867 |
| Q9UK80 | Q86Y13 | USP21 | DZIP3 | 0.865 |
| Q9UK80 | Q96EP0 | USP21 | RNF31 | 0.864 |
| Q9UK80 | Q6FHJ7 | USP21 | SFRP4 | 0.863 |
| Q9UK80 | Q99835 | USP21 | SMO | 0.863 |
| Q9UK80 | Q99743 | USP21 | NPAS2 | 0.856 |
| Q9UK80 | O43318 | USP21 | MAP3K7 | 0.855 |
| Q9UK80 | P48730 | USP21 | CSNK1D | 0.853 |
| Q9UK80 | Q8WYA1 | USP21 | ARNTL2 | 0.852 |
| Q9UK80 | Q9Y573 | USP21 | IPP | 0.852 |
| Q9UK80 | Q9Y462 | USP21 | ZNF711 | 0.852 |
| Q9UK80 | Q9ULW2 | USP21 | FZD10 | 0.850 |
| Q9UK80 | O14757 | USP21 | CHEK1 | 0.849 |
| Q9UK80 | Q9H3N8 | USP21 | HRH4 | 0.849 |
| Q9UK80 | O00421 | USP21 | CCRL2 | 0.846 |
| Q9UK80 | O14920 | USP21 | IKBKB | 0.843 |
| Q9UK80 | Q9C035 | USP21 | TRIM5 | 0.841 |
| Q9UK80 | P50542 | USP21 | PEX5 | 0.837 |
| Q9UK80 | P27540 | USP21 | ARNT | 0.836 |
| Q9UK80 | Q13467 | USP21 | FZD5 | 0.835 |
| Q9UK80 | Q8IYD8 | USP21 | FANCM | 0.835 |
| Q9UK80 | Q86UD3 | USP21 | MARCHF3 | 0.833 |
| Q9UK80 | Q96GD4 | USP21 | AURKB | 0.831 |
| Q9UK80 | P98170 | USP21 | XIAP | 0.827 |
| Q9UK80 | O95976 | USP21 | IGSF6 | 0.827 |
| Q9UK80 | P51965 | USP21 | UBE2E1 | 0.827 |
| Q9UK80 | Q9H4P4 | USP21 | RNF41 | 0.825 |

| | | | | |
|---|---|---|---|---|
| Q9UK80 | Q13635 | USP21 | PTCH1 | 0.824 |
| Q9UK80 | Q9H596 | USP21 | DUSP21 | 0.824 |
| Q9UK80 | Q7LBR1 | USP21 | CHMP1B | 0.823 |
| Q9UK80 | O96009 | USP21 | NAPSA | 0.823 |
| Q9UK80 | Q96Q07 | USP21 | BTBD9 | 0.822 |
| Q9UK80 | Q9Y6C5 | USP21 | PTCH2 | 0.821 |
| Q9UK80 | P19838 | USP21 | NFKB1 | 0.820 |
| Q9UK80 | O00429 | USP21 | DNM1L | 0.817 |
| Q9UK80 | P56645 | USP21 | PER3 | 0.815 |
| Q9UK80 | Q9Y6K9 | USP21 | IKBKG | 0.815 |
| Q9UK80 | Q9HD42 | USP21 | CHMP1A | 0.814 |
| Q9UK80 | Q8N165 | USP21 | PDIK1L | 0.813 |
| Q9UK80 | P30556 | USP21 | AGTR1 | 0.813 |
| Q9UK80 | Q9P2J3 | USP21 | KLHL9 | 0.813 |
| Q9UK80 | Q16778 | USP21 | H2BC21 | 0.812 |
| Q9UK80 | Q9UQB9 | USP21 | AURKC | 0.811 |
| Q9UK80 | Q99728 | USP21 | BARD1 | 0.810 |
| Q9UK80 | Q9NYD6 | USP21 | HOXC10 | 0.810 |
| Q9UK80 | P35226 | USP21 | BMI1 | 0.809 |
| Q9UK80 | Q8NC69 | USP21 | KCTD6 | 0.809 |
| Q9UK80 | Q9BY84 | USP21 | DUSP16 | 0.809 |
| Q9UK80 | P20142 | USP21 | PGC | 0.807 |
| Q9UK80 | Q9ULV1 | USP21 | FZD4 | 0.806 |
| Q9UK80 | Q9UP38 | USP21 | FZD1 | 0.804 |
| Q9UK80 | Q13114 | USP21 | TRAF3 | 0.803 |
| Q9UK80 | Q05193 | USP21 | DNM1 | 0.803 |
| Q9UK80 | P49674 | USP21 | CSNK1E | 0.802 |
| Q9UK80 | Q8N1E6 | USP21 | FBXL14 | 0.802 |
| Q9UK80 | Q14145 | USP21 | KEAP1 | 0.800 |
| Q9UK80 | Q9H173 | USP21 | SIL1 | 0.799 |
| Q9UK80 | P01574 | USP21 | IFNB1 | 0.799 |
| Q9UK80 | P27361 | USP21 | MAPK3 | 0.795 |
| Q9UK80 | Q7L7X3 | USP21 | TAOK1 | 0.795 |
| Q9UK80 | Q96Q05 | USP21 | TRAPPC9 | 0.795 |
| Q9UK80 | Q8IZH2 | USP21 | XRN1 | 0.794 |
| Q9UK80 | P20671 | USP21 | H2AC7 | 0.793 |
| Q9UK80 | Q7L7L0 | USP21 | H2AW | 0.793 |
| Q9UK80 | Q9HC98 | USP21 | NEK6 | 0.791 |
| Q9UK80 | P14091 | USP21 | CTSE | 0.790 |
| Q9UK80 | Q15750 | USP21 | TAB1 | 0.790 |
| Q9UK80 | Q8IYB4 | USP21 | PEX5L | 0.787 |
| Q9UK80 | Q92765 | USP21 | FRZB | 0.784 |
| Q9UK80 | O15321 | USP21 | TM9SF1 | 0.784 |
| Q9UK80 | Q93077 | USP21 | H2AC6 | 0.782 |
| Q9UK80 | P16104 | USP21 | H2AX | 0.782 |
| Q9UK80 | Q96KK5 | USP21 | H2AC12 | 0.781 |
| Q9UK80 | Q96H20 | USP21 | SNF8 | 0.781 |
| Q9UK80 | Q13490 | USP21 | BIRC2 | 0.781 |

| | | | | |
|---|---|---|---|---|
| Q9UK80 | P10145 | USP21 | CXCL8 | 0.779 |
| Q9UK80 | Q96C10 | USP21 | DHX58 | 0.774 |
| Q9UK80 | Q99878 | USP21 | H2AC14 | 0.773 |
| Q9UK80 | P38398 | USP21 | BRCA1 | 0.770 |
| Q9UK80 | Q9H3S7 | USP21 | PTPN23 | 0.769 |
| Q9UK80 | Q96QV6 | USP21 | H2AC1 | 0.768 |
| Q9UK80 | O00144 | USP21 | FZD9 | 0.761 |
| Q9UK80 | Q2KHN1 | USP21 | RNF151 | 0.761 |
| Q9UK80 | P07339 | USP21 | CTSD | 0.761 |
| Q9UK80 | Q9HAU4 | USP21 | SMURF2 | 0.759 |
| Q9UK80 | Q9Y6Y0 | USP21 | IVNS1ABP | 0.759 |
| Q9UK80 | Q5SY16 | USP21 | NOL9 | 0.758 |
| Q9UK80 | O14503 | USP21 | BHLHE40 | 0.758 |
| Q9UK80 | Q9HCE7 | USP21 | SMURF1 | 0.757 |
| Q9UK80 | Q9Y468 | USP21 | L3MBTL1 | 0.757 |
| Q9UK80 | P41743 | USP21 | PRKCI | 0.756 |
| Q9UK80 | P01375 | USP21 | TNF | 0.754 |
| Q9UK80 | Q12933 | USP21 | TRAF2 | 0.753 |
| Q9UK80 | Q9H3R0 | USP21 | KDM4C | 0.753 |
| Q9UK80 | Q9UIX4 | USP21 | KCNG1 | 0.753 |
| Q9UK80 | P50570 | USP21 | DNM2 | 0.753 |
| Q9UK80 | Q8IUE6 | USP21 | H2AC21 | 0.749 |
| Q9UK80 | Q15628 | USP21 | TRADD | 0.749 |
| Q9UK80 | Q9HBZ2 | USP21 | ARNT2 | 0.747 |
| Q9UK80 | O15033 | USP21 | AREL1 | 0.745 |
| Q9UK80 | Q8IUC4 | USP21 | RHPN2 | 0.745 |
| Q9UK80 | Q9NZL4 | USP21 | HSPBP1 | 0.743 |
| Q9UK80 | Q9UQ16 | USP21 | DNM3 | 0.741 |
| Q9UK80 | O75084 | USP21 | FZD7 | 0.738 |
| Q9UK80 | Q96S82 | USP21 | UBL7 | 0.737 |
| Q9UK80 | Q8IYT8 | USP21 | ULK2 | 0.737 |
| Q9UK80 | O00463 | USP21 | TRAF5 | 0.736 |
| Q9UK80 | O76064 | USP21 | RNF8 | 0.735 |
| Q9UK80 | O15516 | USP21 | CLOCK | 0.735 |
| Q9UK80 | Q9GZX5 | USP21 | ZNF350 | 0.727 |
| Q9UK80 | P42566 | USP21 | EPS15 | 0.727 |
| Q9UK80 | Q12888 | USP21 | TP53BP1 | 0.725 |
| Q9UK80 | Q9NVD3 | USP21 | SETD4 | 0.720 |
| Q9UK80 | O14965 | USP21 | AURKA | 0.719 |
| Q9UK80 | Q8TDR2 | USP21 | STK35 | 0.718 |
| Q9UK80 | Q13489 | USP21 | BIRC3 | 0.718 |
| Q9UK80 | O14964 | USP21 | HGS | 0.716 |
| Q9UK80 | P08235 | USP21 | NR3C2 | 0.716 |
| Q9UK80 | Q5D1E8 | USP21 | ZC3H12A | 0.714 |
| Q9UK80 | Q6Q0C0 | USP21 | TRAF7 | 0.712 |
| Q9UK80 | Q9UKI2 | USP21 | CDC42EP3 | 0.712 |
| Q9UK80 | P0C5Y9 | USP21 | H2AB1 | 0.711 |
| Q9UK80 | Q6TDP4 | USP21 | KLHL17 | 0.711 |

| | | | | |
|---|---|---|---|---|
| Q9UK80 | Q86VP6 | USP21 | CAND1 | 0.710 |
| Q9UK80 | Q7Z6Z7 | USP21 | HUWE1 | 0.708 |
| Q5VVQ6 | Q9UK80 | YOD1 | USP21 | 0.696 |
| Q9UK80 | Q9BQI6 | USP21 | SLF1 | 0.701 |
| Q9UK80 | O95817 | USP21 | BAG3 | 0.700 |
| Q9UK80 | Q4L235 | USP21 | AASDH | 0.697 |
| Q9UK80 | Q16539 | USP21 | MAPK14 | 0.694 |
| Q9UK80 | Q8N474 | USP21 | SFRP1 | 0.692 |
| Q9UK80 | Q8TCX5 | USP21 | RHPN1 | 0.688 |
| Q9UK80 | Q13158 | USP21 | FADD | 0.685 |
| Q9UK80 | Q96JH7 | USP21 | VCPIP1 | 0.680 |
| Q9UK80 | O60313 | USP21 | OPA1 | 0.679 |
| Q9UK80 | Q9HBF4 | USP21 | ZFYVE1 | 0.679 |
| Q9UK80 | Q01780 | USP21 | EXOSC10 | 0.677 |
| Q9UK80 | P43405 | USP21 | SYK | 0.672 |
| Q9UK80 | Q6P4R8 | USP21 | NFRKB | 0.670 |
| Q9UK80 | Q92905 | USP21 | COPS5 | 0.669 |
| Q9UK80 | Q96J02 | USP21 | ITCH | 0.668 |
| Q9UK80 | Q92783 | USP21 | STAM | 0.665 |
| Q9UK80 | Q9Y5Q5 | USP21 | CORIN | 0.664 |
| Q9UK80 | Q8IXQ5 | USP21 | KLHL7 | 0.661 |
| Q9UK80 | Q5T4F7 | USP21 | SFRP5 | 0.657 |
| Q9UK80 | Q8WYN0 | USP21 | ATG4A | 0.657 |
| Q9UK80 | O75771 | USP21 | RAD51D | 0.656 |
| Q9UK80 | Q13077 | USP21 | TRAF1 | 0.655 |
| Q9UK80 | Q92995 | USP21 | USP13 | 0.653 |
| Q9UK80 | P31152 | USP21 | MAPK4 | 0.651 |
| Q9UK80 | O95630 | USP21 | STAMBP | 0.649 |
| Q9UK80 | Q8N5Z5 | USP21 | KCTD17 | 0.645 |
| Q9UK80 | Q96PU5 | USP21 | NEDD4L | 0.643 |
| Q9UK80 | P11441 | USP21 | UBL4A | 0.643 |
| Q9UK80 | P35227 | USP21 | PCGF2 | 0.637 |
| Q9UK80 | Q6PHR2 | USP21 | ULK3 | 0.636 |
| Q9UK80 | Q14669 | USP21 | TRIP12 | 0.635 |
| Q9UK80 | P46934 | USP21 | NEDD4 | 0.634 |
| Q9UK80 | Q86TM6 | USP21 | SYVN1 | 0.633 |
| Q9UK80 | O00204 | USP21 | SULT2B1 | 0.632 |
| Q9UK80 | Q8N3Y1 | USP21 | FBXW8 | 0.631 |
| Q9UK80 | O00308 | USP21 | WWP2 | 0.631 |
| Q9UK80 | Q9Y4P1 | USP21 | ATG4B | 0.628 |
| Q9UK80 | O75164 | USP21 | KDM4A | 0.628 |
| Q9UK80 | P45983 | USP21 | MAPK8 | 0.626 |
| Q9UK80 | Q96T68 | USP21 | SETDB2 | 0.621 |
| Q9UK80 | P51668 | USP21 | UBE2D1 | 0.619 |
| Q9UK80 | Q15788 | USP21 | NCOA1 | 0.619 |
| Q9UK80 | Q9Y4E6 | USP21 | WDR7 | 0.615 |
| Q9UK80 | O94953 | USP21 | KDM4B | 0.615 |
| Q9UK80 | Q15047 | USP21 | SETDB1 | 0.615 |

| | | | | |
|---|---|---|---|---|
| Q9UK80 | Q6ZS86 | USP21 | GK5 | 0.614 |
| Q9UK80 | Q9HBJ7 | USP21 | USP29 | 0.614 |
| Q9UK80 | Q92830 | USP21 | KAT2A | 0.612 |
| Q9UK80 | Q9HD20 | USP21 | ATP13A1 | 0.611 |
| Q9UK80 | Q9H7B4 | USP21 | SMYD3 | 0.606 |
| Q9UK80 | Q96K21 | USP21 | ZFYVE19 | 0.606 |
| Q9UK80 | Q9P2P5 | USP21 | HECW2 | 0.604 |
| Q9UK80 | P21580 | USP21 | TNFAIP3 | 0.604 |
| Q9UK80 | P51610 | USP21 | HCFC1 | 0.593 |
| Q9UK80 | Q14139 | USP21 | UBE4A | 0.593 |
| Q9UK80 | Q96FJ0 | USP21 | STAMBPL1 | 0.588 |
| Q9UK80 | Q96FX2 | USP21 | DPH3 | 0.586 |
| Q9UK80 | O15047 | USP21 | SETD1A | 0.585 |
| Q9UK80 | Q8N7F7 | USP21 | UBL4B | 0.583 |
| Q9UK80 | Q6PCT2 | USP21 | FBXL19 | 0.581 |
| Q9UK80 | Q9NRR5 | USP21 | UBQLN4 | 0.580 |
| Q9UK80 | Q92831 | USP21 | KAT2B | 0.578 |
| Q9UK80 | Q6B0I6 | USP21 | KDM4D | 0.575 |
| Q9UK80 | Q5T6C5 | USP21 | ATXN7L2 | 0.575 |
| Q9UK80 | Q92974 | USP21 | ARHGEF2 | 0.574 |
| Q9UK80 | P15170 | USP21 | GSPT1 | 0.574 |
| Q9UK80 | Q9H0M0 | USP21 | WWP1 | 0.572 |
| Q9UK80 | Q15386 | USP21 | UBE3C | 0.570 |
| Q9UK80 | Q15075 | USP21 | EEA1 | 0.569 |
| Q9UK80 | Q96C11 | USP21 | FGGY | 0.568 |
| Q9UK80 | O75385 | USP21 | ULK1 | 0.565 |
| Q9UK80 | Q86TL0 | USP21 | ATG4D | 0.564 |
| Q9UK80 | Q15596 | USP21 | NCOA2 | 0.562 |
| Q9UK80 | P27448 | USP21 | MARK3 | 0.560 |
| Q9UK80 | P80217 | USP21 | IFI35 | 0.559 |
| Q9UK80 | Q9UPU5 | USP21 | USP24 | 0.558 |
| Q9UK80 | P09936 | USP21 | UCHL1 | 0.549 |
| Q9UK80 | Q8WUM4 | USP21 | PDCD6IP | 0.549 |
| Q9UK80 | O00303 | USP21 | EIF3F | 0.549 |
| Q9UK80 | Q7KZI7 | USP21 | MARK2 | 0.547 |
| Q9UK80 | O14980 | USP21 | XPO1 | 0.546 |
| Q9UK80 | Q96HF1 | USP21 | SFRP2 | 0.545 |
| Q9UK80 | P61088 | USP21 | UBE2N | 0.545 |
| Q9UK80 | P54727 | USP21 | RAD23B | 0.544 |
| Q9UK80 | Q15843 | USP21 | NEDD8 | 0.543 |
| Q9UK80 | P54725 | USP21 | RAD23A | 0.540 |
| Q9UK80 | Q9UNE7 | USP21 | STUB1 | 0.539 |
| Q9UK80 | Q693B1 | USP21 | KCTD11 | 0.538 |
| Q9UK80 | Q96KP6 | USP21 | TNIP3 | 0.536 |
| Q9UK80 | Q9Y6Q9 | USP21 | NCOA3 | 0.535 |
| Q9UK80 | Q9BVQ7 | USP21 | SPATA5L1 | 0.531 |
| Q9UK80 | A0AVT1 | USP21 | UBA6 | 0.530 |
| Q9UK80 | P49760 | USP21 | CLK2 | 0.530 |

| | | | | |
|---|---|---|---|---|
| Q9UK80 | Q8N5C8 | USP21 | TAB3 | 0.528 |
| Q9UK80 | Q9NYJ8 | USP21 | TAB2 | 0.528 |
| Q9UK80 | O60861 | USP21 | GAS7 | 0.525 |
| Q9UK80 | Q9UPQ7 | USP21 | PDZRN3 | 0.524 |
| Q9UK80 | P15374 | USP21 | UCHL3 | 0.523 |
| Q9UK80 | Q96CB5 | USP21 | C8orf44 | 0.522 |
| Q9UK80 | P35125 | USP21 | USP6 | 0.521 |
| Q9UK80 | Q8IYD1 | USP21 | GSPT2 | 0.517 |
| Q9UK80 | Q9NZM3 | USP21 | ITSN2 | 0.515 |
| Q9UK80 | Q15291 | USP21 | RBBP5 | 0.514 |
| Q9UK80 | P39023 | USP21 | RPL3 | 0.508 |
| Q9UK80 | Q92901 | USP21 | RPL3L | 0.508 |
| Q9UK80 | Q14191 | USP21 | WRN | 0.507 |
| Q9UK80 | Q15646 | USP21 | OASL | 0.507 |
| Q9UK80 | Q9H4L4 | USP21 | SENP3 | 0.506 |
| Q9UK80 | Q9UBC2 | USP21 | EPS15L1 | 0.505 |
| Q5VVQ6 | Q9NQC7 | YOD1 | CYLD | 0.742 |
| Q9NQC7 | Q9HAW4 | CYLD | CLSPN | 0.994 |
| Q9NQC7 | Q08999 | CYLD | RBL2 | 0.991 |
| Q9NQC7 | Q96RL1 | CYLD | UIMC1 | 0.989 |
| Q9NQC7 | Q00987 | CYLD | MDM2 | 0.985 |
| Q9NQC7 | Q99708 | CYLD | RBBP8 | 0.983 |
| Q9NQC7 | Q9BXW9 | CYLD | FANCD2 | 0.982 |
| Q9NQC7 | P04626 | CYLD | ERBB2 | 0.981 |
| Q9NQC7 | P10070 | CYLD | GLI2 | 0.976 |
| Q9NQC7 | O15379 | CYLD | HDAC3 | 0.973 |
| Q9NQC7 | Q92889 | CYLD | ERCC4 | 0.973 |
| Q9NQC7 | Q13547 | CYLD | HDAC1 | 0.971 |
| Q9NQC7 | P05231 | CYLD | IL6 | 0.971 |
| Q9NQC7 | O95999 | CYLD | BCL10 | 0.969 |
| Q9NQC7 | Q6PIZ9 | CYLD | TRAT1 | 0.968 |
| Q9NQC7 | Q01201 | CYLD | RELB | 0.968 |
| Q9NQC7 | Q96P20 | CYLD | NLRP3 | 0.967 |
| Q9NQC7 | O60674 | CYLD | JAK2 | 0.967 |
| Q9NQC7 | Q9Y5Q3 | CYLD | MAFB | 0.967 |
| Q9NQC7 | Q9HC29 | CYLD | NOD2 | 0.966 |
| Q9NQC7 | P01106 | CYLD | MYC | 0.965 |
| Q9NQC7 | P52701 | CYLD | MSH6 | 0.965 |
| Q9NQC7 | Q86UT6 | CYLD | NLRX1 | 0.964 |
| Q9NQC7 | P43246 | CYLD | MSH2 | 0.964 |
| Q9NQC7 | Q9NPP4 | CYLD | NLRC4 | 0.964 |
| Q9NQC7 | P00533 | CYLD | EGFR | 0.963 |
| Q9NQC7 | O43524 | CYLD | FOXO3 | 0.963 |
| Q9NQC7 | P35222 | CYLD | CTNNB1 | 0.961 |
| Q9NQC7 | Q9UJX3 | CYLD | ANAPC7 | 0.960 |
| Q9NQC7 | Q99683 | CYLD | MAP3K5 | 0.958 |
| Q9NQC7 | O15111 | CYLD | CHUK | 0.956 |

| | | | | |
|---|---|---|---|---|
| Q9NQC7 | Q9Y239 | CYLD | NOD1 | 0.956 |
| Q9NQC7 | P52333 | CYLD | JAK3 | 0.954 |
| Q9NQC7 | Q16620 | CYLD | NTRK2 | 0.954 |
| Q9NQC7 | Q16665 | CYLD | HIF1A | 0.954 |
| Q9NQC7 | Q12778 | CYLD | FOXO1 | 0.953 |
| Q9NQC7 | P62877 | CYLD | RBX1 | 0.953 |
| Q9NQC7 | Q13114 | CYLD | TRAF3 | 0.953 |
| Q9NQC7 | Q96EP0 | CYLD | RNF31 | 0.953 |
| Q9NQC7 | Q13485 | CYLD | SMAD4 | 0.953 |
| Q9NQC7 | O00534 | CYLD | VWA5A | 0.952 |
| Q9NQC7 | O95980 | CYLD | RECK | 0.951 |
| Q9NQC7 | Q14258 | CYLD | TRIM25 | 0.951 |
| Q9NQC7 | Q9BYX4 | CYLD | IFIH1 | 0.951 |
| Q9NQC7 | Q9UPY3 | CYLD | DICER1 | 0.950 |
| Q9NQC7 | Q969S8 | CYLD | HDAC10 | 0.949 |
| Q9NQC7 | Q13315 | CYLD | ATM | 0.949 |
| Q9NQC7 | P37173 | CYLD | TGFBR2 | 0.949 |
| Q9NQC7 | Q01974 | CYLD | ROR2 | 0.949 |
| Q9NQC7 | P48736 | CYLD | PIK3CG | 0.948 |
| Q9NQC7 | P42336 | CYLD | PIK3CA | 0.948 |
| Q9NQC7 | O15169 | CYLD | AXIN1 | 0.948 |
| Q9NQC7 | O75444 | CYLD | MAF | 0.948 |
| Q9NQC7 | P04141 | CYLD | CSF2 | 0.947 |
| Q9NQC7 | Q92844 | CYLD | TANK | 0.947 |
| Q9NQC7 | P25963 | CYLD | NFKBIA | 0.946 |
| Q9NQC7 | Q92993 | CYLD | KAT5 | 0.945 |
| Q9NQC7 | Q86WI3 | CYLD | NLRC5 | 0.945 |
| Q9NQC7 | Q59H18 | CYLD | TNNI3K | 0.945 |
| Q9NQC7 | P23396 | CYLD | RPS3 | 0.945 |
| Q9NQC7 | Q9UM73 | CYLD | ALK | 0.944 |
| Q9NQC7 | Q15208 | CYLD | STK38 | 0.944 |
| Q9NQC7 | Q13309 | CYLD | SKP2 | 0.944 |
| Q9NQC7 | P35228 | CYLD | NOS2 | 0.943 |
| Q9NQC7 | P10914 | CYLD | IRF1 | 0.943 |
| Q9NQC7 | Q9UKV0 | CYLD | HDAC9 | 0.943 |
| Q9NQC7 | P19838 | CYLD | NFKB1 | 0.941 |
| Q9NQC7 | Q9NYL2 | CYLD | MAP3K20 | 0.941 |
| Q9NQC7 | P11831 | CYLD | SRF | 0.941 |
| Q9NQC7 | Q9NUU6 | CYLD | OTULINL | 0.940 |
| Q9NQC7 | Q96BN8 | CYLD | OTULIN | 0.940 |
| Q9NQC7 | O75460 | CYLD | ERN1 | 0.940 |
| Q9NQC7 | P42345 | CYLD | MTOR | 0.940 |
| Q9NQC7 | Q9NSC2 | CYLD | SALL1 | 0.939 |
| Q9NQC7 | Q38SD2 | CYLD | LRRK1 | 0.939 |
| Q9NQC7 | Q07820 | CYLD | MCL1 | 0.938 |
| Q9NQC7 | Q99728 | CYLD | BARD1 | 0.938 |
| Q9NQC7 | O43521 | CYLD | BCL2L11 | 0.937 |
| Q9NQC7 | Q13233 | CYLD | MAP3K1 | 0.937 |

| | | | | |
|---|---|---|---|---|
| Q9NQC7 | P26045 | CYLD | PTPN3 | 0.937 |
| Q9NQC7 | O95644 | CYLD | NFATC1 | 0.937 |
| Q9NQC7 | P19235 | CYLD | EPOR | 0.936 |
| Q9NQC7 | Q9Y572 | CYLD | RIPK3 | 0.935 |
| Q9NQC7 | P00519 | CYLD | ABL1 | 0.935 |
| Q9NQC7 | Q13308 | CYLD | PTK7 | 0.934 |
| Q9NQC7 | P60568 | CYLD | IL2 | 0.933 |
| Q9NQC7 | P08069 | CYLD | IGF1R | 0.933 |
| Q9NQC7 | O95863 | CYLD | SNAI1 | 0.933 |
| Q9NQC7 | Q16288 | CYLD | NTRK3 | 0.932 |
| Q9NQC7 | O00463 | CYLD | TRAF5 | 0.932 |
| Q9NQC7 | Q8WUI4 | CYLD | HDAC7 | 0.931 |
| Q9NQC7 | Q15306 | CYLD | IRF4 | 0.931 |
| Q9NQC7 | O14920 | CYLD | IKBKB | 0.930 |
| Q9NQC7 | Q86VP1 | CYLD | TAX1BP1 | 0.929 |
| Q9NQC7 | Q9P2D0 | CYLD | IBTK | 0.929 |
| Q9NQC7 | Q5W0Q7 | CYLD | USPL1 | 0.929 |
| Q9NQC7 | P57078 | CYLD | RIPK4 | 0.928 |
| Q9NQC7 | Q9NRR4 | CYLD | DROSHA | 0.928 |
| Q9NQC7 | Q99558 | CYLD | MAP3K14 | 0.927 |
| Q9NQC7 | Q01973 | CYLD | ROR1 | 0.926 |
| Q9NQC7 | P30279 | CYLD | CCND2 | 0.926 |
| Q9NQC7 | Q9Y5W3 | CYLD | KLF2 | 0.926 |
| Q9NQC7 | O95352 | CYLD | ATG7 | 0.926 |
| Q9NQC7 | Q6PFW1 | CYLD | PPIP5K1 | 0.926 |
| Q9NQC7 | P46734 | CYLD | MAP2K3 | 0.925 |
| Q9NQC7 | O15519 | CYLD | CFLAR | 0.924 |
| Q9NQC7 | Q92466 | CYLD | DDB2 | 0.924 |
| Q9NQC7 | P51617 | CYLD | IRAK1 | 0.924 |
| Q9NQC7 | P15056 | CYLD | BRAF | 0.923 |
| Q9NQC7 | Q9Y6X2 | CYLD | PIAS3 | 0.922 |
| Q9NQC7 | Q13769 | CYLD | THOC5 | 0.922 |
| Q9NQC7 | Q02556 | CYLD | IRF8 | 0.921 |
| Q9NQC7 | O15524 | CYLD | SOCS1 | 0.920 |
| Q9NQC7 | P17097 | CYLD | ZNF7 | 0.920 |
| Q9NQC7 | P04049 | CYLD | RAF1 | 0.919 |
| Q9NQC7 | Q99759 | CYLD | MAP3K3 | 0.918 |
| Q9NQC7 | Q8NB16 | CYLD | MLKL | 0.917 |
| Q9NQC7 | Q9NP92 | CYLD | MRPS30 | 0.917 |
| Q9NQC7 | Q96CV9 | CYLD | OPTN | 0.916 |
| Q9NQC7 | Q8N2W9 | CYLD | PIAS4 | 0.916 |
| Q9NQC7 | P14373 | CYLD | TRIM27 | 0.916 |
| Q9NQC7 | Q9UBN7 | CYLD | HDAC6 | 0.916 |
| Q9NQC7 | Q9Y5S8 | CYLD | NOX1 | 0.914 |
| Q9NQC7 | Q9BYH8 | CYLD | NFKBIZ | 0.913 |
| Q9NQC7 | Q96DT7 | CYLD | ZBTB10 | 0.912 |
| Q9NQC7 | P45984 | CYLD | MAPK9 | 0.912 |
| Q9NQC7 | O14733 | CYLD | MAP2K7 | 0.912 |

| | | | | |
|---|---|---|---|---|
| Q9NQC7 | Q02779 | CYLD | MAP3K10 | 0.911 |
| Q9NQC7 | Q14790 | CYLD | CASP8 | 0.911 |
| Q9NQC7 | Q14653 | CYLD | IRF3 | 0.910 |
| Q9NQC7 | P43403 | CYLD | ZAP70 | 0.909 |
| Q9NQC7 | P22607 | CYLD | FGFR3 | 0.908 |
| Q9NQC7 | P60484 | CYLD | PTEN | 0.908 |
| Q9NQC7 | Q6IA17 | CYLD | SIGIRR | 0.907 |
| Q9NQC7 | Q9HB75 | CYLD | PIDD1 | 0.907 |
| Q9NQC7 | Q92985 | CYLD | IRF7 | 0.907 |
| Q9NQC7 | Q86WV6 | CYLD | STING1 | 0.906 |
| Q9NQC7 | Q92851 | CYLD | CASP10 | 0.906 |
| Q9NQC7 | P52564 | CYLD | MAP2K6 | 0.906 |
| Q9NQC7 | Q7RTR2 | CYLD | NLRC3 | 0.905 |
| Q9NQC7 | Q15366 | CYLD | PCBP2 | 0.904 |
| Q9NQC7 | Q92664 | CYLD | GTF3A | 0.904 |
| Q9NQC7 | Q13464 | CYLD | ROCK1 | 0.904 |
| Q9NQC7 | Q02750 | CYLD | MAP2K1 | 0.903 |
| Q9NQC7 | P17181 | CYLD | IFNAR1 | 0.903 |
| Q9NQC7 | Q04864 | CYLD | REL | 0.903 |
| Q9NQC7 | P08631 | CYLD | HCK | 0.902 |
| Q9NQC7 | Q8NI38 | CYLD | NFKBID | 0.902 |
| Q9NQC7 | Q14457 | CYLD | BECN1 | 0.902 |
| Q9NQC7 | Q96GD4 | CYLD | AURKB | 0.900 |
| Q9NQC7 | O75626 | CYLD | PRDM1 | 0.900 |
| Q9NQC7 | P21439 | CYLD | ABCB4 | 0.900 |
| Q9NQC7 | Q9HCE7 | CYLD | SMURF1 | 0.899 |
| Q9NQC7 | P42574 | CYLD | CASP3 | 0.898 |
| Q9NQC7 | Q9NSE2 | CYLD | CISH | 0.897 |
| Q9NQC7 | P78524 | CYLD | DENND2B | 0.897 |
| Q9NQC7 | Q99081 | CYLD | TCF12 | 0.897 |
| Q9NQC7 | P28482 | CYLD | MAPK1 | 0.896 |
| Q9NQC7 | P21359 | CYLD | NF1 | 0.896 |
| Q9NQC7 | Q9UL68 | CYLD | MYT1L | 0.895 |
| Q9NQC7 | P42681 | CYLD | TXK | 0.895 |
| Q9NQC7 | Q9UQM7 | CYLD | CAMK2A | 0.894 |
| Q9NQC7 | Q15910 | CYLD | EZH2 | 0.894 |
| Q9NQC7 | Q9UEG4 | CYLD | ZNF629 | 0.894 |
| Q9NQC7 | Q9UQQ2 | CYLD | SH2B3 | 0.893 |
| Q9NQC7 | O14495 | CYLD | PLPP3 | 0.893 |
| Q9NQC7 | Q15653 | CYLD | NFKBIB | 0.891 |
| Q9NQC7 | P31751 | CYLD | AKT2 | 0.891 |
| Q9NQC7 | Q13077 | CYLD | TRAF1 | 0.891 |
| Q9NQC7 | P55212 | CYLD | CASP6 | 0.891 |
| Q9NQC7 | P45985 | CYLD | MAP2K4 | 0.890 |
| Q9NQC7 | P24385 | CYLD | CCND1 | 0.890 |
| Q9NQC7 | P35354 | CYLD | PTGS2 | 0.889 |
| Q9NQC7 | Q9C0F3 | CYLD | ZNF436 | 0.889 |
| Q9NQC7 | Q96CG3 | CYLD | TIFA | 0.889 |

| | | | | |
|---|---|---|---|---|
| Q9NQC7 | P27361 | CYLD | MAPK3 | 0.888 |
| Q9NQC7 | O95376 | CYLD | ARIH2 | 0.887 |
| Q9NQC7 | P15407 | CYLD | FOSL1 | 0.887 |
| Q9NQC7 | O15146 | CYLD | MUSK | 0.886 |
| Q9NQC7 | O96017 | CYLD | CHEK2 | 0.886 |
| Q9NQC7 | O96028 | CYLD | NSD2 | 0.885 |
| Q9NQC7 | Q9UL17 | CYLD | TBX21 | 0.885 |
| Q9NQC7 | Q9H4P4 | CYLD | RNF41 | 0.885 |
| Q9NQC7 | Q96DX4 | CYLD | RSPRY1 | 0.884 |
| Q9NQC7 | P10398 | CYLD | ARAF | 0.884 |
| Q9NQC7 | O60603 | CYLD | TLR2 | 0.883 |
| Q9NQC7 | P43405 | CYLD | SYK | 0.883 |
| Q9NQC7 | P05412 | CYLD | JUN | 0.883 |
| Q9NQC7 | Q8NFD2 | CYLD | ANKK1 | 0.882 |
| Q9NQC7 | P01133 | CYLD | EGF | 0.881 |
| Q9NQC7 | P42771 | CYLD | CDKN2A | 0.880 |
| Q9NQC7 | Q96C10 | CYLD | DHX58 | 0.880 |
| Q9NQC7 | P46531 | CYLD | NOTCH1 | 0.879 |
| Q9NQC7 | Q9P289 | CYLD | STK26 | 0.879 |
| Q9NQC7 | O14641 | CYLD | DVL2 | 0.878 |
| Q9NQC7 | O43187 | CYLD | IRAK2 | 0.877 |
| Q9NQC7 | Q92997 | CYLD | DVL3 | 0.876 |
| Q9NQC7 | Q14164 | CYLD | IKBKE | 0.876 |
| Q9NQC7 | P49841 | CYLD | GSK3B | 0.875 |
| Q9NQC7 | Q9HAU4 | CYLD | SMURF2 | 0.875 |
| Q9NQC7 | Q8IYU2 | CYLD | HACE1 | 0.875 |
| Q9NQC7 | P41279 | CYLD | MAP3K8 | 0.874 |
| Q9NQC7 | Q9NWZ3 | CYLD | IRAK4 | 0.874 |
| Q9NQC7 | P84022 | CYLD | SMAD3 | 0.874 |
| Q9NQC7 | Q5TCX8 | CYLD | MAP3K21 | 0.873 |
| Q9NQC7 | O00635 | CYLD | TRIM38 | 0.873 |
| Q9NQC7 | Q5MCW4 | CYLD | ZNF569 | 0.871 |
| Q9NQC7 | Q9UPQ7 | CYLD | PDZRN3 | 0.871 |
| Q9NQC7 | P56817 | CYLD | BACE1 | 0.871 |
| Q9NQC7 | Q7Z6K4 | CYLD | NRARP | 0.871 |
| Q9NQC7 | P98170 | CYLD | XIAP | 0.870 |
| Q9NQC7 | Q9UK97 | CYLD | FBXO9 | 0.870 |
| Q9NQC7 | Q96CX3 | CYLD | ZNF501 | 0.870 |
| Q9NQC7 | Q2VWP7 | CYLD | PRTG | 0.869 |
| Q9NQC7 | Q15796 | CYLD | SMAD2 | 0.868 |
| Q9NQC7 | Q3MII6 | CYLD | TBC1D25 | 0.867 |
| Q9NQC7 | Q96RT1 | CYLD | ERBIN | 0.867 |
| Q9NQC7 | Q96KB5 | CYLD | PBK | 0.865 |
| Q9NQC7 | Q15349 | CYLD | RPS6KA2 | 0.864 |
| Q9NQC7 | Q9NQ35 | CYLD | NRIP3 | 0.863 |
| Q9NQC7 | Q86YT6 | CYLD | MIB1 | 0.863 |
| Q9NQC7 | Q15942 | CYLD | ZYX | 0.862 |
| Q9NQC7 | P49815 | CYLD | TSC2 | 0.862 |

| | | | | |
|---|---|---|---|---|
| Q9NQC7 | Q9Y243 | CYLD | AKT3 | 0.860 |
| Q9NQC7 | P38398 | CYLD | BRCA1 | 0.860 |
| Q9NQC7 | P58753 | CYLD | TIRAP | 0.859 |
| Q9NQC7 | O14497 | CYLD | ARID1A | 0.857 |
| Q9NQC7 | Q9NR09 | CYLD | BIRC6 | 0.856 |
| Q9NQC7 | Q8TEP8 | CYLD | CEP192 | 0.855 |
| Q9NQC7 | P49281 | CYLD | SLC11A2 | 0.855 |
| Q9NQC7 | O94822 | CYLD | LTN1 | 0.853 |
| Q9NQC7 | Q06187 | CYLD | BTK | 0.853 |
| Q9NQC7 | O43432 | CYLD | EIF4G3 | 0.851 |
| Q9NQC7 | Q8IVT5 | CYLD | KSR1 | 0.851 |
| Q9NQC7 | P19320 | CYLD | VCAM1 | 0.851 |
| Q9NQC7 | P53355 | CYLD | DAPK1 | 0.851 |
| Q9NQC7 | Q99836 | CYLD | MYD88 | 0.851 |
| Q9NQC7 | Q9H0E2 | CYLD | TOLLIP | 0.850 |
| Q9NQC7 | P50750 | CYLD | CDK9 | 0.850 |
| Q9NQC7 | Q6XUX3 | CYLD | DSTYK | 0.849 |
| Q9NQC7 | Q16552 | CYLD | IL17A | 0.849 |
| Q9NQC7 | Q9GZX7 | CYLD | AICDA | 0.848 |
| Q9NQC7 | Q00653 | CYLD | NFKB2 | 0.848 |
| Q9NQC7 | Q6ZMN7 | CYLD | PDZRN4 | 0.847 |
| Q9NQC7 | Q13635 | CYLD | PTCH1 | 0.847 |
| Q9NQC7 | Q9NTX7 | CYLD | RNF146 | 0.845 |
| Q9NQC7 | Q9BYV9 | CYLD | BACH2 | 0.844 |
| Q9NQC7 | Q06124 | CYLD | PTPN11 | 0.844 |
| Q9NQC7 | Q96JY6 | CYLD | PDLIM2 | 0.844 |
| Q9NQC7 | Q16548 | CYLD | BCL2A1 | 0.843 |
| Q9NQC7 | Q07817 | CYLD | BCL2L1 | 0.842 |
| Q9NQC7 | O15033 | CYLD | AREL1 | 0.842 |
| Q9NQC7 | P10275 | CYLD | AR | 0.842 |
| Q9NQC7 | Q9Y616 | CYLD | IRAK3 | 0.841 |
| Q9NQC7 | P16410 | CYLD | CTLA4 | 0.841 |
| Q9NQC7 | Q9Y462 | CYLD | ZNF711 | 0.840 |
| Q9NQC7 | P22681 | CYLD | CBL | 0.838 |
| Q9NQC7 | P13232 | CYLD | IL7 | 0.837 |
| Q9NQC7 | O75936 | CYLD | BBOX1 | 0.837 |
| Q9NQC7 | Q9UMX9 | CYLD | SLC45A2 | 0.836 |
| Q9NQC7 | Q96EQ8 | CYLD | RNF125 | 0.835 |
| Q9NQC7 | Q8WV24 | CYLD | PHLDA1 | 0.834 |
| Q9NQC7 | Q9UBE8 | CYLD | NLK | 0.834 |
| Q9NQC7 | P04062 | CYLD | GBA | 0.834 |
| Q9NQC7 | P29597 | CYLD | TYK2 | 0.833 |
| Q9NQC7 | Q13490 | CYLD | BIRC2 | 0.833 |
| Q9NQC7 | Q96JH7 | CYLD | VCPIP1 | 0.833 |
| Q9NQC7 | Q15465 | CYLD | SHH | 0.832 |
| Q9NQC7 | Q13751 | CYLD | LAMB3 | 0.832 |
| Q9NQC7 | P01568 | CYLD | IFNA21 | 0.832 |
| Q9NQC7 | Q9ULT8 | CYLD | HECTD1 | 0.831 |

| | | | | |
|---|---|---|---|---|
| Q9NQC7 | Q969H0 | CYLD | FBXW7 | 0.830 |
| Q9NQC7 | Q6ZVD8 | CYLD | PHLPP2 | 0.829 |
| Q9NQC7 | O75204 | CYLD | TMEM127 | 0.829 |
| Q9NQC7 | P55210 | CYLD | CASP7 | 0.828 |
| Q9NQC7 | Q96EB6 | CYLD | SIRT1 | 0.827 |
| Q9NQC7 | P42081 | CYLD | CD86 | 0.826 |
| Q9NQC7 | Q15628 | CYLD | TRADD | 0.826 |
| Q9NQC7 | Q01780 | CYLD | EXOSC10 | 0.821 |
| Q9NQC7 | P25445 | CYLD | FAS | 0.819 |
| Q9NQC7 | Q96P31 | CYLD | FCRL3 | 0.818 |
| Q9NQC7 | Q2KHN1 | CYLD | RNF151 | 0.815 |
| Q9NQC7 | Q9NR97 | CYLD | TLR8 | 0.815 |
| Q9NQC7 | P98164 | CYLD | LRP2 | 0.811 |
| Q9NQC7 | P09769 | CYLD | FGR | 0.811 |
| Q9NQC7 | P01563 | CYLD | IFNA2 | 0.811 |
| Q9NQC7 | O60346 | CYLD | PHLPP1 | 0.811 |
| Q9NQC7 | O75113 | CYLD | N4BP1 | 0.810 |
| Q9NQC7 | P17275 | CYLD | JUNB | 0.810 |
| Q9NQC7 | O14964 | CYLD | HGS | 0.810 |
| Q9NQC7 | P04156 | CYLD | PRNP | 0.808 |
| Q9NQC7 | Q12888 | CYLD | TP53BP1 | 0.808 |
| Q9NQC7 | P48200 | CYLD | IREB2 | 0.807 |
| Q9NQC7 | P06241 | CYLD | FYN | 0.805 |
| Q9NQC7 | Q16584 | CYLD | MAP3K11 | 0.805 |
| Q9NQC7 | Q96Q89 | CYLD | KIF20B | 0.804 |
| Q9NQC7 | Q9Y2D8 | CYLD | SSX2IP | 0.800 |
| Q9NQC7 | P42575 | CYLD | CASP2 | 0.800 |
| Q9NQC7 | P51812 | CYLD | RPS6KA3 | 0.798 |
| Q9NQC7 | Q9UPN4 | CYLD | CEP131 | 0.798 |
| Q9NQC7 | P23458 | CYLD | JAK1 | 0.795 |
| Q9NQC7 | Q5D1E8 | CYLD | ZC3H12A | 0.793 |
| Q9NQC7 | Q13158 | CYLD | FADD | 0.789 |
| Q9NQC7 | Q15750 | CYLD | TAB1 | 0.788 |
| Q9NQC7 | Q5S007 | CYLD | LRRK2 | 0.786 |
| Q9NQC7 | Q7Z6Z7 | CYLD | HUWE1 | 0.784 |
| Q9NQC7 | P28562 | CYLD | DUSP1 | 0.783 |
| Q9NQC7 | Q71RS6 | CYLD | SLC24A5 | 0.782 |
| Q9NQC7 | Q05823 | CYLD | RNASEL | 0.782 |
| Q9NQC7 | Q76N89 | CYLD | HECW1 | 0.782 |
| Q9NQC7 | P45983 | CYLD | MAPK8 | 0.781 |
| Q9NQC7 | P01571 | CYLD | IFNA17 | 0.780 |
| Q9NQC7 | Q16539 | CYLD | MAPK14 | 0.775 |
| Q9NQC7 | P25116 | CYLD | F2R | 0.775 |
| Q9NQC7 | Q9UJU2 | CYLD | LEF1 | 0.775 |
| Q9NQC7 | P12931 | CYLD | SRC | 0.773 |
| Q9NQC7 | Q8N6H7 | CYLD | ARFGAP2 | 0.772 |
| Q9NQC7 | P51965 | CYLD | UBE2E1 | 0.770 |
| Q9NQC7 | Q96CH1 | CYLD | GPR146 | 0.770 |

| | | | | |
|---|---|---|---|---|
| Q9NQC7 | P10747 | CYLD | CD28 | 0.769 |
| Q9NQC7 | P07948 | CYLD | LYN | 0.768 |
| Q9NQC7 | Q13489 | CYLD | BIRC3 | 0.768 |
| Q9NQC7 | P55211 | CYLD | CASP9 | 0.763 |
| Q9NQC7 | P30405 | CYLD | PPIF | 0.763 |
| Q9NQC7 | O95235 | CYLD | KIF20A | 0.762 |
| Q9NQC7 | P35625 | CYLD | TIMP3 | 0.762 |
| Q9NQC7 | P01584 | CYLD | IL1B | 0.762 |
| Q9NQC7 | Q96PU5 | CYLD | NEDD4L | 0.761 |
| Q9NQC7 | P53539 | CYLD | FOSB | 0.760 |
| Q9NQC7 | P80192 | CYLD | MAP3K9 | 0.760 |
| Q9NQC7 | Q14344 | CYLD | GNA13 | 0.757 |
| Q9NQC7 | Q9Y4X5 | CYLD | ARIH1 | 0.755 |
| Q9NQC7 | P04003 | CYLD | C4BPA | 0.754 |
| Q9NQC7 | Q70EL1 | CYLD | USP54 | 0.754 |
| Q9NQC7 | Q9BWF2 | CYLD | TRAIP | 0.751 |
| Q9NQC7 | Q96RQ1 | CYLD | ERGIC2 | 0.750 |
| Q9NQC7 | P09874 | CYLD | PARP1 | 0.748 |
| Q9NQC7 | P61073 | CYLD | CXCR4 | 0.747 |
| Q9NQC7 | Q96CA5 | CYLD | BIRC7 | 0.747 |
| Q9NQC7 | O00308 | CYLD | WWP2 | 0.746 |
| Q9NQC7 | Q16777 | CYLD | H2AC20 | 0.746 |
| Q9NQC7 | Q9Y2E6 | CYLD | DTX4 | 0.744 |
| Q9NQC7 | Q53EL6 | CYLD | PDCD4 | 0.744 |
| Q9NQC7 | P00540 | CYLD | MOS | 0.743 |
| Q9NQC7 | P29466 | CYLD | CASP1 | 0.743 |
| Q9NQC7 | Q15399 | CYLD | TLR1 | 0.741 |
| Q9NQC7 | Q7Z6A9 | CYLD | BTLA | 0.739 |
| Q9NQC7 | O15455 | CYLD | TLR3 | 0.739 |
| Q9NQC7 | Q6ZUJ8 | CYLD | PIK3AP1 | 0.736 |
| Q9NQC7 | Q96J84 | CYLD | KIRREL1 | 0.733 |
| Q9NQC7 | Q8TBB1 | CYLD | LNX1 | 0.729 |
| Q9NQC7 | P46934 | CYLD | NEDD4 | 0.729 |
| Q9NQC7 | P53779 | CYLD | MAPK10 | 0.729 |
| Q9NQC7 | P62993 | CYLD | GRB2 | 0.727 |
| Q9NQC7 | P01574 | CYLD | IFNB1 | 0.721 |
| Q9NQC7 | Q05086 | CYLD | UBE3A | 0.716 |
| Q9NQC7 | P15621 | CYLD | ZNF44 | 0.716 |
| Q9NQC7 | Q96J02 | CYLD | ITCH | 0.716 |
| Q9NQC7 | P21580 | CYLD | TNFAIP3 | 0.712 |
| Q9NQC7 | Q9HBL0 | CYLD | TNS1 | 0.712 |
| Q9NQC7 | Q9UDY8 | CYLD | MALT1 | 0.709 |
| Q9NQC7 | Q92830 | CYLD | KAT2A | 0.707 |
| Q9NQC7 | Q92783 | CYLD | STAM | 0.707 |
| Q9NQC7 | Q9NYK1 | CYLD | TLR7 | 0.705 |
| Q9NQC7 | Q7L1W4 | CYLD | LRRC8D | 0.705 |
| Q9NQC7 | Q13291 | CYLD | SLAMF1 | 0.705 |
| Q9NQC7 | Q9Y5E7 | CYLD | PCDHB2 | 0.704 |

| | | | | |
|---|---|---|---|---|
| Q9NQC7 | Q9BZS1 | CYLD | FOXP3 | 0.936 |
| A6NNY8 | Q9NQC7 | USP27X | CYLD | 0.503 |
| O75317 | Q9NQC7 | USP12 | CYLD | 0.507 |
| O75604 | Q9NQC7 | USP2 | CYLD | 0.509 |
| O94782 | Q9NQC7 | USP1 | CYLD | 0.564 |
| O94966 | Q9NQC7 | USP19 | CYLD | 0.516 |
| P09936 | Q9NQC7 | UCHL1 | CYLD | 0.530 |
| P15374 | Q9NQC7 | UCHL3 | CYLD | 0.532 |
| P54252 | Q9NQC7 | ATXN3 | CYLD | 0.597 |
| P62068 | Q9NQC7 | USP46 | CYLD | 0.516 |
| Q15040 | Q9NQC7 | JOSD1 | CYLD | 0.550 |
| Q53GS9 | Q9NQC7 | USP39 | CYLD | 0.537 |
| Q70CQ1 | Q9NQC7 | USP49 | CYLD | 0.518 |
| Q70CQ3 | Q9NQC7 | USP30 | CYLD | 0.565 |
| Q70EL2 | Q9NQC7 | USP45 | CYLD | 0.560 |
| Q70EL3 | Q9NQC7 | USP50 | CYLD | 0.518 |
| Q70EL4 | Q9NQC7 | USP43 | CYLD | 0.510 |
| Q8TEY7 | Q9NQC7 | USP33 | CYLD | 0.557 |
| Q92560 | Q9NQC7 | BAP1 | CYLD | 0.538 |
| Q9H0E7 | Q9NQC7 | USP44 | CYLD | 0.533 |
| Q9H3M9 | Q9NQC7 | ATXN3L | CYLD | 0.599 |
| Q9NQC7 | P50591 | CYLD | TNFSF10 | 0.701 |
| Q9NQC7 | P19474 | CYLD | TRIM21 | 0.701 |
| Q9NQC7 | Q15418 | CYLD | RPS6KA1 | 0.698 |
| Q9NQC7 | Q53G59 | CYLD | KLHL12 | 0.698 |
| Q9NQC7 | Q5XPI4 | CYLD | RNF123 | 0.697 |
| Q9NQC7 | P01130 | CYLD | LDLR | 0.696 |
| Q9NQC7 | Q9BTL4 | CYLD | IER2 | 0.695 |
| Q9NQC7 | Q92831 | CYLD | KAT2B | 0.693 |
| Q9NQC7 | P29965 | CYLD | CD40LG | 0.693 |
| Q9NQC7 | O95071 | CYLD | UBR5 | 0.692 |
| Q9NQC7 | P51878 | CYLD | CASP5 | 0.692 |
| Q9NQC7 | P12830 | CYLD | CDH1 | 0.692 |
| Q9NQC7 | Q9H0M0 | CYLD | WWP1 | 0.690 |
| Q9NQC7 | Q92995 | CYLD | USP13 | 0.687 |
| Q9NQC7 | Q5VT06 | CYLD | CEP350 | 0.685 |
| Q9NQC7 | Q9Y6Q6 | CYLD | TNFRSF11A | 0.684 |
| Q9NQC7 | Q9HBJ7 | CYLD | USP29 | 0.682 |
| Q9NQC7 | O75594 | CYLD | PGLYRP1 | 0.682 |
| Q9NQC7 | P31944 | CYLD | CASP14 | 0.680 |
| Q9NQC7 | P10145 | CYLD | CXCL8 | 0.679 |
| Q9NQC7 | O14974 | CYLD | PPP1R12A | 0.678 |
| Q9NQC7 | Q9UI42 | CYLD | CPA4 | 0.677 |
| Q9NQC7 | P01100 | CYLD | FOS | 0.676 |
| Q9NQC7 | P35125 | CYLD | USP6 | 0.676 |
| Q9NQC7 | Q8N1G4 | CYLD | LRRC47 | 0.675 |
| Q9NQC7 | Q9UPI3 | CYLD | FLVCR2 | 0.674 |
| Q9NQC7 | P16220 | CYLD | CREB1 | 0.672 |

| | | | | |
|---|---|---|---|---|
| Q9NQC7 | P43489 | CYLD | TNFRSF4 | 0.672 |
| Q9NQC7 | Q9BXL7 | CYLD | CARD11 | 0.670 |
| Q9NQC7 | Q9P0U3 | CYLD | SENP1 | 0.670 |
| Q9NQC7 | Q9UPU5 | CYLD | USP24 | 0.670 |
| Q9NQC7 | P40763 | CYLD | STAT3 | 0.669 |
| Q9NQC7 | P11441 | CYLD | UBL4A | 0.667 |
| Q9NQC7 | Q13501 | CYLD | SQSTM1 | 0.667 |
| Q9NQC7 | P48728 | CYLD | AMT | 0.666 |
| Q9NQC7 | Q96RD9 | CYLD | FCRL5 | 0.661 |
| Q9NQC7 | Q96BY6 | CYLD | DOCK10 | 0.661 |
| Q9NQC7 | Q96DZ5 | CYLD | CLIP3 | 0.660 |
| Q9NQC7 | Q8IUD6 | CYLD | RNF135 | 0.659 |
| Q9NQC7 | P42224 | CYLD | STAT1 | 0.657 |
| Q9NQC7 | P03372 | CYLD | ESR1 | 0.656 |
| Q9NQC7 | O95630 | CYLD | STAMBP | 0.653 |
| Q9NQC7 | Q9H257 | CYLD | CARD9 | 0.653 |
| Q9NQC7 | Q9BXR5 | CYLD | TLR10 | 0.651 |
| Q9NQC7 | Q9BWT7 | CYLD | CARD10 | 0.647 |
| Q9NQC7 | Q9UNE0 | CYLD | EDAR | 0.646 |
| Q9NQC7 | P18847 | CYLD | ATF3 | 0.642 |
| Q9NQC7 | Q5T4S7 | CYLD | UBR4 | 0.641 |
| Q9NQC7 | Q14765 | CYLD | STAT4 | 0.639 |
| Q9NQC7 | P19022 | CYLD | CDH2 | 0.638 |
| Q9NQC7 | Q15025 | CYLD | TNIP1 | 0.635 |
| Q9NQC7 | Q96S82 | CYLD | UBL7 | 0.634 |
| Q9NQC7 | Q8IUD2 | CYLD | ERC1 | 0.633 |
| Q9NQC7 | Q9UGI0 | CYLD | ZRANB1 | 0.633 |
| Q9NQC7 | Q02241 | CYLD | KIF23 | 0.632 |
| Q9NQC7 | Q16520 | CYLD | BATF | 0.630 |
| Q9NQC7 | P48052 | CYLD | CPA2 | 0.626 |
| Q9NQC7 | Q9BUZ4 | CYLD | TRAF4 | 0.625 |
| Q9NQC7 | Q13838 | CYLD | DDX39B | 0.625 |
| Q9NQC7 | Q07011 | CYLD | TNFRSF9 | 0.624 |
| Q9NQC7 | P52630 | CYLD | STAT2 | 0.621 |
| Q9NQC7 | P17535 | CYLD | JUND | 0.620 |
| Q9NQC7 | Q9UN67 | CYLD | PCDHB10 | 0.616 |
| Q9NQC7 | Q9NYA1 | CYLD | SPHK1 | 0.610 |
| Q9NQC7 | Q9NQT8 | CYLD | KIF13B | 0.608 |
| Q9NQC7 | Q9UNE7 | CYLD | STUB1 | 0.606 |
| Q9NQC7 | Q13829 | CYLD | TNFAIP1 | 0.605 |
| Q9NQC7 | O14788 | CYLD | TNFSF11 | 0.602 |
| Q9NQC7 | P01375 | CYLD | TNF | 0.600 |
| Q9NQC7 | P12980 | CYLD | LYL1 | 0.599 |
| Q9NQC7 | Q8IUC6 | CYLD | TICAM1 | 0.598 |
| Q9NQC7 | Q9UJV9 | CYLD | DDX41 | 0.597 |
| Q9NQC7 | O14490 | CYLD | DLGAP1 | 0.596 |
| Q9NQC7 | Q92905 | CYLD | COPS5 | 0.595 |
| Q9NQC7 | O00571 | CYLD | DDX3X | 0.589 |

| | | | | |
|---|---|---|---|---|
| Q9NQC7 | Q9Y5H2 | CYLD | PCDHGA11 | 0.588 |
| Q9NQC7 | O00566 | CYLD | MPHOSPH10 | 0.586 |
| Q9NQC7 | Q9C0C9 | CYLD | UBE2O | 0.585 |
| Q9NQC7 | P02741 | CYLD | CRP | 0.581 |
| Q9NQC7 | Q9BZL1 | CYLD | UBL5 | 0.581 |
| Q9NQC7 | O00206 | CYLD | TLR4 | 0.579 |
| Q9NQC7 | A0AVT1 | CYLD | UBA6 | 0.579 |
| Q9NQC7 | P0DP08 | CYLD | IGHV4-38-2 | 0.577 |
| Q9NQC7 | Q9Y566 | CYLD | SHANK1 | 0.577 |
| Q9NQC7 | Q4LE39 | CYLD | ARID4B | 0.570 |
| Q9NQC7 | Q96FJ0 | CYLD | STAMBPL1 | 0.569 |
| Q9NQC7 | O75688 | CYLD | PPM1B | 0.567 |
| Q9NQC7 | P14151 | CYLD | SELL | 0.567 |
| Q9NQC7 | Q8NHG8 | CYLD | ZNRF2 | 0.563 |
| Q9NQC7 | O00213 | CYLD | APBB1 | 0.560 |
| Q9NQC7 | Q86V81 | CYLD | ALYREF | 0.558 |
| Q9NQC7 | Q07343 | CYLD | PDE4B | 0.558 |
| Q9NQC7 | Q9NS68 | CYLD | TNFRSF19 | 0.558 |
| Q9NQC7 | Q8N7F7 | CYLD | UBL4B | 0.557 |
| Q9NQC7 | Q7L622 | CYLD | G2E3 | 0.552 |
| Q9NQC7 | P29279 | CYLD | CCN2 | 0.550 |
| Q9NQC7 | Q15646 | CYLD | OASL | 0.548 |
| Q9NQC7 | Q9UQ90 | CYLD | SPG7 | 0.546 |
| Q9NQC7 | Q9H158 | CYLD | PCDHAC1 | 0.545 |
| Q9NQC7 | Q9NPI1 | CYLD | BRD7 | 0.545 |
| Q9NQC7 | P56945 | CYLD | BCAR1 | 0.543 |
| Q9NQC7 | P27986 | CYLD | PIK3R1 | 0.539 |
| Q9NQC7 | P16455 | CYLD | MGMT | 0.537 |
| Q9NQC7 | P15516 | CYLD | HTN3 | 0.536 |
| Q9NQC7 | Q01804 | CYLD | OTUD4 | 0.536 |
| Q9NQC7 | Q96KP6 | CYLD | TNIP3 | 0.534 |
| Q9NQC7 | Q676U5 | CYLD | ATG16L1 | 0.533 |
| Q9NQC7 | O60602 | CYLD | TLR5 | 0.532 |
| Q9NQC7 | Q9Y4C4 | CYLD | MFHAS1 | 0.530 |
| Q9NQC7 | Q8IZR5 | CYLD | CMTM4 | 0.530 |
| Q9NQC7 | P26358 | CYLD | DNMT1 | 0.521 |
| Q9NQC7 | Q8TAF3 | CYLD | WDR48 | 0.517 |
| Q9NQC7 | Q13191 | CYLD | CBLB | 0.517 |
| Q9NQC7 | Q8NFA0 | CYLD | USP32 | 0.513 |
| Q9NQC7 | Q02548 | CYLD | PAX5 | 0.509 |
| Q9NQC7 | P45974 | CYLD | USP5 | 0.508 |
| Q9NQC7 | Q8WVY7 | CYLD | UBLCP1 | 0.507 |
| Q9NQC7 | Q9BVI0 | CYLD | PHF20 | 0.507 |
| Q9NQC7 | P08138 | CYLD | NGFR | 0.505 |
| Q9NQC7 | Q9NZC7 | CYLD | WWOX | 0.502 |
| Q9P275 | Q9NQC7 | USP36 | CYLD | 0.541 |
| Q9P2H5 | Q9NQC7 | USP35 | CYLD | 0.543 |
| Q9UMW8 | Q9NQC7 | USP18 | CYLD | 0.545 |

| | | | | |
|---|---|---|---|---|
| Q9UPT9 | Q9NQC7 | USP22 | CYLD | 0.575 |
| Q9Y5K5 | Q9NQC7 | UCHL5 | CYLD | 0.541 |
| P46736 | P51587 | BRCC3 | BRCA2 | 0.993 |
| P46736 | Q9HAW4 | BRCC3 | CLSPN | 0.992 |
| P46736 | P12956 | BRCC3 | XRCC6 | 0.989 |
| P46736 | Q9NVI1 | BRCC3 | FANCI | 0.972 |
| P46736 | Q96RL1 | BRCC3 | UIMC1 | 0.965 |
| P46736 | P04637 | BRCC3 | TP53 | 0.965 |
| P46736 | O43679 | BRCC3 | LDB2 | 0.965 |
| P46736 | Q86U70 | BRCC3 | LDB1 | 0.965 |
| P46736 | Q00987 | BRCC3 | MDM2 | 0.963 |
| P46736 | O15151 | BRCC3 | MDM4 | 0.963 |
| P46736 | O60934 | BRCC3 | NBN | 0.961 |
| P46736 | Q13547 | BRCC3 | HDAC1 | 0.959 |
| P46736 | P30304 | BRCC3 | CDC25A | 0.959 |
| P46736 | P62877 | BRCC3 | RBX1 | 0.958 |
| P46736 | Q92769 | BRCC3 | HDAC2 | 0.956 |
| P46736 | Q99708 | BRCC3 | RBBP8 | 0.955 |
| P46736 | P30307 | BRCC3 | CDC25C | 0.955 |
| P46736 | Q01831 | BRCC3 | XPC | 0.950 |
| P46736 | Q13042 | BRCC3 | CDC16 | 0.949 |
| P46736 | P17812 | BRCC3 | CTPS1 | 0.946 |
| P46736 | Q9NRF8 | BRCC3 | CTPS2 | 0.946 |
| P46736 | Q92535 | BRCC3 | PIGC | 0.945 |
| P46736 | P04626 | BRCC3 | ERBB2 | 0.941 |
| P46736 | Q9BXW9 | BRCC3 | FANCD2 | 0.940 |
| P46736 | Q9Y4K3 | BRCC3 | TRAF6 | 0.939 |
| P46736 | P06746 | BRCC3 | POLB | 0.939 |
| P46736 | Q14094 | BRCC3 | CCNI | 0.936 |
| P46736 | P62249 | BRCC3 | RPS16 | 0.929 |
| P46736 | Q86YC2 | BRCC3 | PALB2 | 0.928 |
| P46736 | Q16589 | BRCC3 | CCNG2 | 0.928 |
| P46736 | Q96BN8 | BRCC3 | OTULIN | 0.926 |
| P46736 | Q8IYW5 | BRCC3 | RNF168 | 0.926 |
| P46736 | P78396 | BRCC3 | CCNA1 | 0.922 |
| P46736 | P14635 | BRCC3 | CCNB1 | 0.921 |
| P46736 | P04179 | BRCC3 | SOD2 | 0.921 |
| P46736 | Q9UJX3 | BRCC3 | ANAPC7 | 0.916 |
| P46736 | P15880 | BRCC3 | RPS2 | 0.915 |
| P46736 | O96020 | BRCC3 | CCNE2 | 0.914 |
| P46736 | Q6ZMN8 | BRCC3 | CCNI2 | 0.914 |
| P46736 | Q96BM1 | BRCC3 | ANKRD9 | 0.912 |
| P46736 | P17181 | BRCC3 | IFNAR1 | 0.909 |
| P46736 | Q9NS91 | BRCC3 | RAD18 | 0.908 |
| P46736 | Q9UBF6 | BRCC3 | RNF7 | 0.907 |
| P46736 | Q8WTU0 | BRCC3 | DDI1 | 0.907 |
| P46736 | P42356 | BRCC3 | PI4KA | 0.907 |

| | | | | |
|---|---|---|---|---|
| P46736 | Q5T1A1 | BRCC3 | DCST2 | 0.906 |
| P46736 | P23396 | BRCC3 | RPS3 | 0.906 |
| P46736 | O95067 | BRCC3 | CCNB2 | 0.906 |
| P46736 | P51959 | BRCC3 | CCNG1 | 0.905 |
| P46736 | Q92466 | BRCC3 | DDB2 | 0.905 |
| P46736 | P05412 | BRCC3 | JUN | 0.904 |
| P46736 | Q99496 | BRCC3 | RNF2 | 0.903 |
| P46736 | P22674 | BRCC3 | CCNO | 0.902 |
| P46736 | Q5TDH0 | BRCC3 | DDI2 | 0.902 |
| P46736 | Q16667 | BRCC3 | CDKN3 | 0.899 |
| P46736 | P60228 | BRCC3 | EIF3E | 0.896 |
| P46736 | Q92993 | BRCC3 | KAT5 | 0.896 |
| P46736 | P27694 | BRCC3 | RPA1 | 0.894 |
| P46736 | P42677 | BRCC3 | RPS27 | 0.894 |
| P46736 | Q71UM5 | BRCC3 | RPS27L | 0.894 |
| P46736 | Q9HB75 | BRCC3 | PIDD1 | 0.893 |
| P46736 | P20248 | BRCC3 | CCNA2 | 0.893 |
| P46736 | Q06210 | BRCC3 | GFPT1 | 0.893 |
| P46736 | O94808 | BRCC3 | GFPT2 | 0.891 |
| P46736 | Q02556 | BRCC3 | IRF8 | 0.890 |
| P46736 | Q07002 | BRCC3 | CDK18 | 0.888 |
| P46736 | O60729 | BRCC3 | CDC14B | 0.886 |
| P46736 | Q9H8S5 | BRCC3 | CCNP | 0.886 |
| P46736 | P43246 | BRCC3 | MSH2 | 0.883 |
| P46736 | P25963 | BRCC3 | NFKBIA | 0.882 |
| P46736 | Q5W0Q7 | BRCC3 | USPL1 | 0.882 |
| P46736 | Q9BQI9 | BRCC3 | NRIP2 | 0.881 |
| P46736 | Q5M7Z0 | BRCC3 | RNFT1 | 0.881 |
| P46736 | Q5T197 | BRCC3 | DCST1 | 0.881 |
| P46736 | Q9BVJ7 | BRCC3 | DUSP23 | 0.879 |
| P46736 | A2A3K4 | BRCC3 | PTPDC1 | 0.878 |
| P46736 | Q9NY27 | BRCC3 | PPP4R2 | 0.877 |
| P46736 | Q9NQ35 | BRCC3 | NRIP3 | 0.876 |
| P46736 | Q9Y5L0 | BRCC3 | TNPO3 | 0.875 |
| P46736 | O00329 | BRCC3 | PIK3CD | 0.874 |
| P46736 | Q96EP1 | BRCC3 | CHFR | 0.873 |
| P46736 | Q8N2W9 | BRCC3 | PIAS4 | 0.872 |
| P46736 | Q9UQ84 | BRCC3 | EXO1 | 0.872 |
| P46736 | Q6VAB6 | BRCC3 | KSR2 | 0.870 |
| P46736 | O43542 | BRCC3 | XRCC3 | 0.870 |
| P46736 | O14757 | BRCC3 | CHEK1 | 0.869 |
| P46736 | Q8NB59 | BRCC3 | SYT14 | 0.869 |
| P46736 | P62277 | BRCC3 | RPS13 | 0.869 |
| P46736 | O15315 | BRCC3 | RAD51B | 0.868 |
| P46736 | Q00526 | BRCC3 | CDK3 | 0.865 |
| P46736 | Q9BQG2 | BRCC3 | NUDT12 | 0.862 |
| P46736 | P06493 | BRCC3 | CDK1 | 0.860 |
| P46736 | Q8N635 | BRCC3 | MEIOB | 0.859 |

| | | | | |
|---|---|---|---|---|
| P46736 | Q8TDX7 | BRCC3 | NEK7 | 0.857 |
| P46736 | Q9BXS5 | BRCC3 | AP1M1 | 0.855 |
| P46736 | P51965 | BRCC3 | UBE2E1 | 0.854 |
| P46736 | Q9H4P4 | BRCC3 | RNF41 | 0.853 |
| P46736 | Q9BTX1 | BRCC3 | NDC1 | 0.852 |
| P46736 | P52701 | BRCC3 | MSH6 | 0.852 |
| P46736 | O60733 | BRCC3 | PLA2G6 | 0.852 |
| P46736 | Q8WXK4 | BRCC3 | ASB12 | 0.851 |
| P46736 | P30260 | BRCC3 | CDC27 | 0.850 |
| P46736 | Q8TCY5 | BRCC3 | MRAP | 0.850 |
| P46736 | Q9C0B5 | BRCC3 | ZDHHC5 | 0.850 |
| P46736 | Q9UNH5 | BRCC3 | CDC14A | 0.848 |
| P46736 | O43242 | BRCC3 | PSMD3 | 0.848 |
| P46736 | Q9UNS2 | BRCC3 | COPS3 | 0.848 |
| P46736 | Q8NEB9 | BRCC3 | PIK3C3 | 0.847 |
| P46736 | Q99879 | BRCC3 | H2BC14 | 0.844 |
| P46736 | Q8IYD8 | BRCC3 | FANCM | 0.844 |
| P46736 | Q7LBR1 | BRCC3 | CHMP1B | 0.843 |
| P46736 | O60814 | BRCC3 | H2BC12 | 0.842 |
| P46736 | Q99880 | BRCC3 | H2BC13 | 0.841 |
| P46736 | O95352 | BRCC3 | ATG7 | 0.840 |
| P46736 | Q8NC69 | BRCC3 | KCTD6 | 0.840 |
| P46736 | Q96EX2 | BRCC3 | RNFT2 | 0.839 |
| P46736 | P33778 | BRCC3 | H2BC3 | 0.839 |
| P46736 | P19838 | BRCC3 | NFKB1 | 0.839 |
| P46736 | O00750 | BRCC3 | PIK3C2B | 0.837 |
| P46736 | Q96EB6 | BRCC3 | SIRT1 | 0.837 |
| P46736 | Q9Y259 | BRCC3 | CHKB | 0.837 |
| P46736 | Q13144 | BRCC3 | EIF2B5 | 0.836 |
| P46736 | Q9UKV5 | BRCC3 | AMFR | 0.836 |
| P46736 | P35226 | BRCC3 | BMI1 | 0.836 |
| P46736 | Q9HD42 | BRCC3 | CHMP1A | 0.835 |
| P46736 | Q6IN85 | BRCC3 | PPP4R3A | 0.835 |
| P46736 | Q9Y575 | BRCC3 | ASB3 | 0.834 |
| P46736 | Q99728 | BRCC3 | BARD1 | 0.834 |
| P46736 | P45984 | BRCC3 | MAPK9 | 0.833 |
| P46736 | P58876 | BRCC3 | H2BC5 | 0.833 |
| P46736 | Q99877 | BRCC3 | H2BC15 | 0.833 |
| P46736 | Q8TD20 | BRCC3 | SLC2A12 | 0.832 |
| P46736 | Q93079 | BRCC3 | H2BC9 | 0.830 |
| P46736 | P24864 | BRCC3 | CCNE1 | 0.830 |
| P46736 | P06899 | BRCC3 | H2BC11 | 0.829 |
| P46736 | Q13418 | BRCC3 | ILK | 0.829 |
| P46736 | P57053 | BRCC3 | H2BS1 | 0.828 |
| P46736 | Q9UBF8 | BRCC3 | PI4KB | 0.828 |
| P46736 | Q495B1 | BRCC3 | ANKDD1A | 0.827 |
| P46736 | O00443 | BRCC3 | PIK3C2A | 0.827 |
| P46736 | Q9H672 | BRCC3 | ASB7 | 0.827 |

| | | | | |
|---|---|---|---|---|
| P46736 | Q7Z6K1 | BRCC3 | THAP5 | 0.826 |
| P46736 | Q969W1 | BRCC3 | ZDHHC16 | 0.826 |
| P46736 | Q13976 | BRCC3 | PRKG1 | 0.825 |
| P46736 | Q96A08 | BRCC3 | H2BC1 | 0.825 |
| P46736 | O94921 | BRCC3 | CDK14 | 0.824 |
| P46736 | P57078 | BRCC3 | RIPK4 | 0.823 |
| P46736 | P60484 | BRCC3 | PTEN | 0.823 |
| P46736 | P22102 | BRCC3 | GART | 0.823 |
| P46736 | P20749 | BRCC3 | BCL3 | 0.822 |
| P46736 | Q8WU17 | BRCC3 | RNF139 | 0.821 |
| P46736 | Q9H765 | BRCC3 | ASB8 | 0.819 |
| P46736 | O75943 | BRCC3 | RAD17 | 0.818 |
| P46736 | Q16778 | BRCC3 | H2BC21 | 0.817 |
| P46736 | P23527 | BRCC3 | H2BC17 | 0.817 |
| P46736 | Q8IUH5 | BRCC3 | ZDHHC17 | 0.816 |
| P46736 | P42338 | BRCC3 | PIK3CB | 0.816 |
| P46736 | Q99942 | BRCC3 | RNF5 | 0.816 |
| P46736 | Q96Q05 | BRCC3 | TRAPPC9 | 0.816 |
| P46736 | Q8WXK3 | BRCC3 | ASB13 | 0.816 |
| P46736 | Q8WXK1 | BRCC3 | ASB15 | 0.815 |
| P46736 | Q53RE8 | BRCC3 | ANKRD39 | 0.814 |
| P46736 | Q7Z4I7 | BRCC3 | LIMS2 | 0.814 |
| P46736 | Q17RQ9 | BRCC3 | NKPD1 | 0.814 |
| P46736 | Q96MT1 | BRCC3 | RNF145 | 0.813 |
| P46736 | Q504Q3 | BRCC3 | PAN2 | 0.812 |
| P46736 | Q96Q40 | BRCC3 | CDK15 | 0.812 |
| P46736 | P42336 | BRCC3 | PIK3CA | 0.811 |
| P46736 | Q495M9 | BRCC3 | USH1G | 0.809 |
| P46736 | P48736 | BRCC3 | PIK3CG | 0.809 |
| P46736 | B4E2M5 | BRCC3 | ANKRD66 | 0.809 |
| P46736 | Q15653 | BRCC3 | NFKBIB | 0.808 |
| P46736 | Q7L7L0 | BRCC3 | H2AW | 0.808 |
| P46736 | P20671 | BRCC3 | H2AC7 | 0.808 |
| P46736 | P52429 | BRCC3 | DGKE | 0.808 |
| P46736 | Q9Y6H5 | BRCC3 | SNCAIP | 0.808 |
| P46736 | Q8NI38 | BRCC3 | NFKBID | 0.806 |
| P46736 | A6NHY2 | BRCC3 | ANKDD1B | 0.806 |
| P46736 | P38398 | BRCC3 | BRCA1 | 0.806 |
| P46736 | Q8WWX0 | BRCC3 | ASB5 | 0.805 |
| P46736 | O75925 | BRCC3 | PIAS1 | 0.805 |
| P46736 | P50570 | BRCC3 | DNM2 | 0.804 |
| P46736 | Q8N5Q1 | BRCC3 | FAM71E2 | 0.804 |
| P46736 | Q8NAG6 | BRCC3 | ANKLE1 | 0.804 |
| P46736 | P31930 | BRCC3 | UQCRC1 | 0.803 |
| P46736 | Q92547 | BRCC3 | TOPBP1 | 0.803 |
| P46736 | O75832 | BRCC3 | PSMD10 | 0.802 |
| P46736 | Q5VYY1 | BRCC3 | ANKRD22 | 0.802 |
| P46736 | O75747 | BRCC3 | PIK3C2G | 0.801 |

| | | | | |
|---|---|---|---|---|
| P46736 | Q9BZ19 | BRCC3 | ANKRD60 | 0.801 |
| P46736 | P78406 | BRCC3 | RAE1 | 0.800 |
| P46736 | Q13315 | BRCC3 | ATM | 0.800 |
| P46736 | Q9H9E1 | BRCC3 | ANKRA2 | 0.800 |
| P46736 | E5RJM6 | BRCC3 | ANKRD65 | 0.799 |
| P46736 | Q93077 | BRCC3 | H2AC6 | 0.799 |
| P46736 | P42773 | BRCC3 | CDKN2C | 0.798 |
| P46736 | Q8TC84 | BRCC3 | FANK1 | 0.798 |
| P46736 | Q96KK5 | BRCC3 | H2AC12 | 0.798 |
| P46736 | Q9Y574 | BRCC3 | ASB4 | 0.796 |
| P46736 | Q6FI13 | BRCC3 | H2AC18 | 0.796 |
| P46736 | O76064 | BRCC3 | RNF8 | 0.795 |
| P46736 | Q96DX5 | BRCC3 | ASB9 | 0.795 |
| P46736 | Q9Y576 | BRCC3 | ASB1 | 0.795 |
| P46736 | Q17RD7 | BRCC3 | SYT16 | 0.794 |
| P46736 | Q8IUH4 | BRCC3 | ZDHHC13 | 0.794 |
| P46736 | Q96HA7 | BRCC3 | TONSL | 0.793 |
| P46736 | Q8WXI3 | BRCC3 | ASB10 | 0.792 |
| P46736 | Q8WWL7 | BRCC3 | CCNB3 | 0.792 |
| P46736 | Q96QE2 | BRCC3 | SLC2A13 | 0.791 |
| P46736 | Q86WC6 | BRCC3 | PPP1R27 | 0.791 |
| P46736 | Q9ULH0 | BRCC3 | KIDINS220 | 0.791 |
| P46736 | P42345 | BRCC3 | MTOR | 0.791 |
| P46736 | Q99878 | BRCC3 | H2AC14 | 0.791 |
| P46736 | Q8NCN4 | BRCC3 | RNF169 | 0.790 |
| P46736 | P55273 | BRCC3 | CDKN2D | 0.789 |
| P46736 | Q8WXH4 | BRCC3 | ASB11 | 0.788 |
| P46736 | B5ME19 | BRCC3 | EIF3CL | 0.788 |
| P46736 | Q99613 | BRCC3 | EIF3C | 0.788 |
| P46736 | Q9H3S7 | BRCC3 | PTPN23 | 0.788 |
| P46736 | Q9Y468 | BRCC3 | L3MBTL1 | 0.787 |
| P46736 | Q7Z713 | BRCC3 | ANKRD37 | 0.786 |
| P46736 | Q00537 | BRCC3 | CDK17 | 0.785 |
| P46736 | Q96QV6 | BRCC3 | H2AC1 | 0.785 |
| P46736 | O75390 | BRCC3 | CS | 0.784 |
| P46736 | P61421 | BRCC3 | ATP6V0D1 | 0.784 |
| P46736 | Q8N8Y2 | BRCC3 | ATP6V0D2 | 0.784 |
| P46736 | Q96LR5 | BRCC3 | UBE2E2 | 0.783 |
| P46736 | Q8WXE0 | BRCC3 | CASKIN2 | 0.783 |
| P46736 | Q9Y385 | BRCC3 | UBE2J1 | 0.780 |
| P46736 | Q7Z6K4 | BRCC3 | NRARP | 0.780 |
| P46736 | A6NK59 | BRCC3 | ASB14 | 0.780 |
| P46736 | Q86W74 | BRCC3 | ANKRD46 | 0.779 |
| P46736 | Q969K4 | BRCC3 | ABTB1 | 0.779 |
| P46736 | Q9H9Y6 | BRCC3 | POLR1B | 0.778 |
| P46736 | Q9NWX5 | BRCC3 | ASB6 | 0.778 |
| P46736 | Q16777 | BRCC3 | H2AC20 | 0.778 |
| P46736 | O43502 | BRCC3 | RAD51C | 0.776 |

| | | | | |
|---|---|---|---|---|
| P46736 | Q9Y2I7 | BRCC3 | PIKFYVE | 0.774 |
| P46736 | Q9ULC8 | BRCC3 | ZDHHC8 | 0.773 |
| P46736 | P42772 | BRCC3 | CDKN2B | 0.771 |
| P46736 | O96017 | BRCC3 | CHEK2 | 0.771 |
| P46736 | Q8N9B4 | BRCC3 | ANKRD42 | 0.771 |
| P46736 | Q969T4 | BRCC3 | UBE2E3 | 0.770 |
| P46736 | P24941 | BRCC3 | CDK2 | 0.770 |
| P46736 | P58546 | BRCC3 | MTPN | 0.769 |
| P46736 | Q8IUE6 | BRCC3 | H2AC21 | 0.769 |
| P46736 | O60313 | BRCC3 | OPA1 | 0.768 |
| P46736 | O95528 | BRCC3 | SLC2A10 | 0.767 |
| P46736 | O14593 | BRCC3 | RFXANK | 0.766 |
| P46736 | Q6NXT1 | BRCC3 | ANKRD54 | 0.766 |
| P46736 | P0C0S5 | BRCC3 | H2AZ1 | 0.766 |
| P46736 | Q00536 | BRCC3 | CDK16 | 0.765 |
| P46736 | P52434 | BRCC3 | POLR2H | 0.765 |
| P46736 | Q8TBB6 | BRCC3 | SLC7A14 | 0.764 |
| P46736 | Q8N8V4 | BRCC3 | ANKS4B | 0.763 |
| P46736 | O43246 | BRCC3 | SLC7A4 | 0.763 |
| P46736 | Q9H832 | BRCC3 | UBE2Z | 0.763 |
| P46736 | Q15327 | BRCC3 | ANKRD1 | 0.763 |
| P46736 | Q8IUC4 | BRCC3 | RHPN2 | 0.762 |
| P46736 | A6NGH8 | BRCC3 | ANKRD61 | 0.761 |
| P46736 | P0C5Y9 | BRCC3 | H2AB1 | 0.761 |
| P46736 | Q6Q0C0 | BRCC3 | TRAF7 | 0.761 |
| P46736 | P62847 | BRCC3 | RPS24 | 0.760 |
| P46736 | Q96NW4 | BRCC3 | ANKRD27 | 0.760 |
| P46736 | P52569 | BRCC3 | SLC7A2 | 0.760 |
| P46736 | Q9NUQ2 | BRCC3 | AGPAT5 | 0.759 |
| P46736 | Q8IV38 | BRCC3 | ANKMY2 | 0.759 |
| P46736 | Q9UHW9 | BRCC3 | SLC12A6 | 0.757 |
| P46736 | Q06547 | BRCC3 | GABPB1 | 0.757 |
| P46736 | P08069 | BRCC3 | IGF1R | 0.756 |
| P46736 | P42771 | BRCC3 | CDKN2A | 0.756 |
| P46736 | Q96I13 | BRCC3 | ABHD8 | 0.756 |
| P46736 | O95429 | BRCC3 | BAG4 | 0.755 |
| P46736 | Q9BYH8 | BRCC3 | NFKBIZ | 0.754 |
| P46736 | Q8NB46 | BRCC3 | ANKRD52 | 0.753 |
| P46736 | Q7Z5J8 | BRCC3 | ANKAR | 0.753 |
| P46736 | O00221 | BRCC3 | NFKBIE | 0.752 |
| P46736 | Q9BSK4 | BRCC3 | FEM1A | 0.752 |
| P46736 | Q9GZX5 | BRCC3 | ZNF350 | 0.750 |
| P46736 | Q9NY64 | BRCC3 | SLC2A8 | 0.750 |
| P46736 | Q8WY07 | BRCC3 | SLC7A3 | 0.749 |
| P46736 | Q8N6D5 | BRCC3 | ANKRD29 | 0.748 |
| P46736 | Q9NR09 | BRCC3 | BIRC6 | 0.748 |
| P46736 | O14867 | BRCC3 | BACH1 | 0.748 |
| P46736 | Q9UJX5 | BRCC3 | ANAPC4 | 0.748 |

| | | | | |
|---|---|---|---|---|
| P46736 | Q9BQI6 | BRCC3 | SLF1 | 0.746 |
| P46736 | O75439 | BRCC3 | PMPCB | 0.746 |
| P46736 | Q8IV13 | BRCC3 | CCNJL | 0.746 |
| P46736 | Q92527 | BRCC3 | ANKRD7 | 0.745 |
| P46736 | P00740 | BRCC3 | F9 | 0.745 |
| P46736 | Q96JP0 | BRCC3 | FEM1C | 0.745 |
| P46736 | Q12834 | BRCC3 | CDC20 | 0.744 |
| P46736 | Q16204 | BRCC3 | CCDC6 | 0.744 |
| P46736 | Q12888 | BRCC3 | TP53BP1 | 0.743 |
| P46736 | A6NCQ9 | BRCC3 | RNF222 | 0.743 |
| P46736 | P61086 | BRCC3 | UBE2K | 0.743 |
| P46736 | Q92485 | BRCC3 | SMPDL3B | 0.743 |
| P46736 | Q96SD1 | BRCC3 | DCLRE1C | 0.743 |
| P46736 | Q5T9C9 | BRCC3 | PIP5KL1 | 0.742 |
| P46736 | Q5D1E8 | BRCC3 | ZC3H12A | 0.742 |
| P46736 | Q8WXD9 | BRCC3 | CASKIN1 | 0.742 |
| P46736 | Q9UL15 | BRCC3 | BAG5 | 0.739 |
| P46736 | Q6NY19 | BRCC3 | KANK3 | 0.738 |
| P46736 | Q9NQT4 | BRCC3 | EXOSC5 | 0.738 |
| P46736 | Q9BZL1 | BRCC3 | UBL5 | 0.737 |
| P46736 | Q9UGQ3 | BRCC3 | SLC2A6 | 0.736 |
| P46736 | Q86XT4 | BRCC3 | TRIM50 | 0.736 |
| P46736 | Q8WVL7 | BRCC3 | ANKRD49 | 0.735 |
| P46736 | Q6ZVZ8 | BRCC3 | ASB18 | 0.734 |
| P46736 | Q8TCX5 | BRCC3 | RHPN1 | 0.732 |
| P46736 | P11802 | BRCC3 | CDK4 | 0.732 |
| P46736 | O95630 | BRCC3 | STAMBP | 0.732 |
| P46736 | Q13561 | BRCC3 | DCTN2 | 0.731 |
| P46736 | O75771 | BRCC3 | RAD51D | 0.731 |
| P46736 | O95292 | BRCC3 | VAPB | 0.730 |
| P46736 | P30825 | BRCC3 | SLC7A1 | 0.729 |
| P46736 | Q9NR50 | BRCC3 | EIF2B3 | 0.728 |
| P46736 | P49643 | BRCC3 | PRIM2 | 0.728 |
| P46736 | O14578 | BRCC3 | CIT | 0.727 |
| P46736 | Q15406 | BRCC3 | NR6A1 | 0.727 |
| P46736 | P62993 | BRCC3 | GRB2 | 0.727 |
| P46736 | Q92625 | BRCC3 | ANKS1A | 0.727 |
| P46736 | Q8IYU2 | BRCC3 | HACE1 | 0.726 |
| P46736 | Q9H2X9 | BRCC3 | SLC12A5 | 0.726 |
| P46736 | Q00534 | BRCC3 | CDK6 | 0.725 |
| P46736 | Q9H9Q4 | BRCC3 | NHEJ1 | 0.725 |
| P46736 | Q5T7N3 | BRCC3 | KANK4 | 0.725 |
| P46736 | Q63ZY3 | BRCC3 | KANK2 | 0.724 |
| P46736 | Q96S59 | BRCC3 | RANBP9 | 0.723 |
| P46736 | Q9P0K7 | BRCC3 | RAI14 | 0.722 |
| P46736 | Q6QNK2 | BRCC3 | ADGRD1 | 0.721 |
| P46736 | Q9UJX2 | BRCC3 | CDC23 | 0.720 |
| P46736 | O95672 | BRCC3 | ECEL1 | 0.720 |

| | | | | |
|---|---|---|---|---|
| P46736 | Q9UGJ0 | BRCC3 | PRKAG2 | 0.719 |
| P46736 | Q96Q27 | BRCC3 | ASB2 | 0.718 |
| P46736 | Q96JH7 | BRCC3 | VCPIP1 | 0.718 |
| P46736 | Q8N9V6 | BRCC3 | ANKRD53 | 0.718 |
| P46736 | Q68DC2 | BRCC3 | ANKS6 | 0.718 |
| P46736 | Q9UP95 | BRCC3 | SLC12A4 | 0.718 |
| P46736 | Q5VVX9 | BRCC3 | UBE2U | 0.718 |
| P46736 | Q6P6B7 | BRCC3 | ANKRD16 | 0.717 |
| P46736 | Q9GZV1 | BRCC3 | ANKRD2 | 0.716 |
| P46736 | Q92905 | BRCC3 | COPS5 | 0.714 |
| P46736 | Q7L2H7 | BRCC3 | EIF3M | 0.713 |
| P46736 | O43390 | BRCC3 | HNRNPR | 0.712 |
| P46736 | P61758 | BRCC3 | VBP1 | 0.711 |
| P46736 | Q9NXR5 | BRCC3 | ANKRD10 | 0.710 |
| P46736 | Q9NU02 | BRCC3 | ANKEF1 | 0.710 |
| P46736 | Q5BJH7 | BRCC3 | YIF1B | 0.710 |
| P46736 | A7E2S9 | BRCC3 | ANKRD30BL | 0.709 |
| P46736 | Q9BXX2 | BRCC3 | ANKRD30B | 0.709 |
| P46736 | Q86SG2 | BRCC3 | ANKRD23 | 0.707 |
| P46736 | Q7Z6Z7 | BRCC3 | HUWE1 | 0.707 |
| P46736 | P35227 | BRCC3 | PCGF2 | 0.706 |
| P46736 | P45983 | BRCC3 | MAPK8 | 0.705 |
| P46736 | Q9UBT2 | BRCC3 | UBA2 | 0.704 |
| P46736 | Q9Y283 | BRCC3 | INVS | 0.704 |
| P46736 | Q6ZW76 | BRCC3 | ANKS3 | 0.703 |
| P46736 | O60237 | BRCC3 | PPP1R12B | 0.702 |
| Q5VVQ6 | P46736 | YOD1 | BRCC3 | 0.821 |
| O00487 | P46736 | PSMD14 | BRCC3 | 0.529 |
| O75317 | P46736 | USP12 | BRCC3 | 0.599 |
| O75604 | P46736 | USP2 | BRCC3 | 0.626 |
| O94782 | P46736 | USP1 | BRCC3 | 0.677 |
| O94966 | P46736 | USP19 | BRCC3 | 0.538 |
| O95630 | P46736 | STAMBP | BRCC3 | 0.505 |
| P09936 | P46736 | UCHL1 | BRCC3 | 0.619 |
| P15374 | P46736 | UCHL3 | BRCC3 | 0.624 |
| P35125 | P46736 | USP6 | BRCC3 | 0.590 |
| P46736 | Q96NS5 | BRCC3 | ASB16 | 0.701 |
| P46736 | Q92783 | BRCC3 | STAM | 0.701 |
| P46736 | Q92995 | BRCC3 | USP13 | 0.700 |
| P46736 | Q9UGI9 | BRCC3 | PRKAG3 | 0.700 |
| P46736 | Q8IXI1 | BRCC3 | RHOT2 | 0.698 |
| P46736 | P51668 | BRCC3 | UBE2D1 | 0.698 |
| P46736 | O95817 | BRCC3 | BAG3 | 0.698 |
| P46736 | Q3KP44 | BRCC3 | ANKRD55 | 0.697 |
| P46736 | Q6ZVH7 | BRCC3 | ESPNL | 0.696 |
| P46736 | P11441 | BRCC3 | UBL4A | 0.696 |
| P46736 | Q96Q15 | BRCC3 | SMG1 | 0.695 |
| P46736 | P54619 | BRCC3 | PRKAG1 | 0.694 |

| | | | | |
|---|---|---|---|---|
| P46736 | Q96J02 | BRCC3 | ITCH | 0.694 |
| P46736 | Q9P2R3 | BRCC3 | ANKFY1 | 0.693 |
| P46736 | Q8NCJ5 | BRCC3 | SPRYD3 | 0.692 |
| P46736 | Q14678 | BRCC3 | KANK1 | 0.692 |
| P46736 | Q13347 | BRCC3 | EIF3I | 0.691 |
| P46736 | Q6VN20 | BRCC3 | RANBP10 | 0.691 |
| P46736 | Q00653 | BRCC3 | NFKB2 | 0.690 |
| P46736 | Q7Z6G8 | BRCC3 | ANKS1B | 0.689 |
| P46736 | P54278 | BRCC3 | PMS2 | 0.688 |
| P46736 | Q8WWH4 | BRCC3 | ASZ1 | 0.687 |
| P46736 | Q96I34 | BRCC3 | PPP1R16A | 0.685 |
| P46736 | Q96SF2 | BRCC3 | CCT8L2 | 0.684 |
| P46736 | Q96S82 | BRCC3 | UBL7 | 0.684 |
| P46736 | Q9Y666 | BRCC3 | SLC12A7 | 0.682 |
| P46736 | Q8IWZ3 | BRCC3 | ANKHD1 | 0.682 |
| P46736 | P00451 | BRCC3 | F8 | 0.679 |
| P46736 | Q9UJG1 | BRCC3 | MOSPD1 | 0.678 |
| P46736 | O75425 | BRCC3 | MOSPD3 | 0.678 |
| P46736 | Q8N8B7 | BRCC3 | TCEANC | 0.677 |
| P46736 | Q96FJ0 | BRCC3 | STAMBPL1 | 0.677 |
| P46736 | Q05823 | BRCC3 | RNASEL | 0.677 |
| P46736 | Q15819 | BRCC3 | UBE2V2 | 0.677 |
| P46736 | Q86VP6 | BRCC3 | CAND1 | 0.676 |
| P46736 | Q9Y2G4 | BRCC3 | ANKRD6 | 0.674 |
| P46736 | Q8TAK5 | BRCC3 | GABPB2 | 0.671 |
| P46736 | Q9UIF8 | BRCC3 | BAZ2B | 0.670 |
| P46736 | O15084 | BRCC3 | ANKRD28 | 0.669 |
| P46736 | P04150 | BRCC3 | NR3C1 | 0.669 |
| P46736 | P03372 | BRCC3 | ESR1 | 0.668 |
| P46736 | Q9Y5Y6 | BRCC3 | ST14 | 0.668 |
| P46736 | O75762 | BRCC3 | TRPA1 | 0.666 |
| P46736 | P34896 | BRCC3 | SHMT1 | 0.665 |
| P46736 | P34897 | BRCC3 | SHMT2 | 0.665 |
| P46736 | Q13535 | BRCC3 | ATR | 0.664 |
| P46736 | Q9BRT3 | BRCC3 | MIEN1 | 0.664 |
| P46736 | Q86YT6 | BRCC3 | MIB1 | 0.663 |
| P46736 | P09936 | BRCC3 | UCHL1 | 0.663 |
| P46736 | Q13685 | BRCC3 | AAMP | 0.663 |
| P46736 | Q9BX63 | BRCC3 | BRIP1 | 0.661 |
| P46736 | Q9UI10 | BRCC3 | EIF2B4 | 0.659 |
| P46736 | O96028 | BRCC3 | NSD2 | 0.659 |
| P46736 | Q9HBJ7 | BRCC3 | USP29 | 0.658 |
| P46736 | Q96T49 | BRCC3 | PPP1R16B | 0.658 |
| P46736 | Q6P4R8 | BRCC3 | NFRKB | 0.658 |
| P46736 | P40692 | BRCC3 | MLH1 | 0.657 |
| P46736 | O96008 | BRCC3 | TOMM40 | 0.654 |
| P46736 | Q969M1 | BRCC3 | TOMM40L | 0.654 |
| P46736 | Q1L5Z9 | BRCC3 | LONRF2 | 0.653 |

| | | | | |
|---|---|---|---|---|
| P46736 | P55011 | BRCC3 | SLC12A2 | 0.650 |
| P46736 | Q4UJ75 | BRCC3 | ANKRD20A4l | 0.649 |
| P46736 | Q9UK73 | BRCC3 | FEM1B | 0.649 |
| P46736 | O75164 | BRCC3 | KDM4A | 0.648 |
| P46736 | Q8NA23 | BRCC3 | WDR31 | 0.646 |
| P46736 | A6NC57 | BRCC3 | ANKRD62 | 0.646 |
| P46736 | P61088 | BRCC3 | UBE2N | 0.646 |
| P46736 | Q01484 | BRCC3 | ANK2 | 0.645 |
| P46736 | Q5TYW2 | BRCC3 | ANKRD20A1 | 0.644 |
| P46736 | Q9H5N1 | BRCC3 | RABEP2 | 0.644 |
| P46736 | O43396 | BRCC3 | TXNL1 | 0.644 |
| P46736 | Q13404 | BRCC3 | UBE2V1 | 0.644 |
| P46736 | Q96AX9 | BRCC3 | MIB2 | 0.642 |
| P46736 | P80217 | BRCC3 | IFI35 | 0.642 |
| P46736 | Q8TBX8 | BRCC3 | PIP4K2C | 0.641 |
| P46736 | Q5VUR7 | BRCC3 | ANKRD20A3l | 0.641 |
| P46736 | P04275 | BRCC3 | VWF | 0.640 |
| P46736 | P15374 | BRCC3 | UCHL3 | 0.640 |
| P46736 | Q96TA2 | BRCC3 | YME1L1 | 0.638 |
| P46736 | Q8IXK0 | BRCC3 | PHC2 | 0.637 |
| P46736 | P19793 | BRCC3 | RXRA | 0.637 |
| P46736 | P13861 | BRCC3 | PRKAR2A | 0.635 |
| P46736 | Q9C0C9 | BRCC3 | UBE2O | 0.635 |
| P46736 | P48454 | BRCC3 | PPP3CC | 0.634 |
| P46736 | P51124 | BRCC3 | GZMM | 0.634 |
| P46736 | P63279 | BRCC3 | UBE2I | 0.634 |
| P46736 | Q5SQ80 | BRCC3 | ANKRD20A2l | 0.634 |
| P46736 | P62837 | BRCC3 | UBE2D2 | 0.633 |
| P46736 | P52198 | BRCC3 | RND2 | 0.632 |
| P46736 | Q8N7F7 | BRCC3 | UBL4B | 0.631 |
| P46736 | P35125 | BRCC3 | USP6 | 0.631 |
| P46736 | O15360 | BRCC3 | FANCA | 0.630 |
| P46736 | P48426 | BRCC3 | PIP4K2A | 0.630 |
| P46736 | P13521 | BRCC3 | SCG2 | 0.629 |
| P46736 | Q9HCD6 | BRCC3 | TANC2 | 0.629 |
| P46736 | Q96IJ6 | BRCC3 | GMPPA | 0.629 |
| P46736 | Q14669 | BRCC3 | TRIP12 | 0.629 |
| P46736 | P51530 | BRCC3 | DNA2 | 0.628 |
| P46736 | Q05086 | BRCC3 | UBE3A | 0.627 |
| P46736 | Q9UBS9 | BRCC3 | SUCO | 0.627 |
| P46736 | O00762 | BRCC3 | UBE2C | 0.625 |
| P46736 | Q8WUM4 | BRCC3 | PDCD6IP | 0.624 |
| P46736 | Q15018 | BRCC3 | ABRAXAS2 | 0.624 |
| P46736 | Q6UWZ7 | BRCC3 | ABRAXAS1 | 0.624 |
| P46736 | Q9Y5P6 | BRCC3 | GMPPB | 0.624 |
| P46736 | Q9NY72 | BRCC3 | SCN3B | 0.623 |
| P46736 | A6QL64 | BRCC3 | ANKRD36 | 0.621 |
| P46736 | Q9H2K2 | BRCC3 | TNKS2 | 0.620 |

| | | | | |
|---|---|---|---|---|
| P46736 | A0AVT1 | BRCC3 | UBA6 | 0.619 |
| P46736 | Q14139 | BRCC3 | UBE4A | 0.619 |
| P46736 | O75717 | BRCC3 | WDHD1 | 0.619 |
| P46736 | Q9BZL4 | BRCC3 | PPP1R12C | 0.617 |
| P46736 | Q92820 | BRCC3 | GGH | 0.617 |
| P46736 | B1AK53 | BRCC3 | ESPN | 0.612 |
| P46736 | Q9Y2X8 | BRCC3 | UBE2D4 | 0.610 |
| P46736 | P63220 | BRCC3 | RPS21 | 0.610 |
| P46736 | Q9UIF9 | BRCC3 | BAZ2A | 0.609 |
| P46736 | P43490 | BRCC3 | NAMPT | 0.608 |
| P46736 | Q6ZS86 | BRCC3 | GK5 | 0.608 |
| P46736 | Q9BVQ7 | BRCC3 | SPATA5L1 | 0.606 |
| P46736 | P46100 | BRCC3 | ATRX | 0.606 |
| P46736 | Q15843 | BRCC3 | NEDD8 | 0.605 |
| P46736 | Q86TM6 | BRCC3 | SYVN1 | 0.602 |
| P46736 | O95071 | BRCC3 | UBR5 | 0.600 |
| P46736 | Q99755 | BRCC3 | PIP5K1A | 0.599 |
| P46736 | P13010 | BRCC3 | XRCC5 | 0.596 |
| P46736 | P63241 | BRCC3 | EIF5A | 0.596 |
| P46736 | O00571 | BRCC3 | DDX3X | 0.593 |
| P46736 | Q9UPU5 | BRCC3 | USP24 | 0.592 |
| P46736 | Q9BXX3 | BRCC3 | ANKRD30A | 0.592 |
| P46736 | P41002 | BRCC3 | CCNF | 0.591 |
| P46736 | Q9ULJ7 | BRCC3 | ANKRD50 | 0.591 |
| P46736 | Q9GZP4 | BRCC3 | PITHD1 | 0.590 |
| P46736 | Q9GZV4 | BRCC3 | EIF5A2 | 0.588 |
| P46736 | Q9UHC1 | BRCC3 | MLH3 | 0.587 |
| P46736 | Q92830 | BRCC3 | KAT2A | 0.586 |
| P46736 | P04844 | BRCC3 | RPN2 | 0.585 |
| P46736 | O15372 | BRCC3 | EIF3H | 0.585 |
| P46736 | P63146 | BRCC3 | UBE2B | 0.583 |
| P46736 | O43252 | BRCC3 | PAPSS1 | 0.583 |
| P46736 | P82675 | BRCC3 | MRPS5 | 0.583 |
| P46736 | P62195 | BRCC3 | PSMC5 | 0.583 |
| P46736 | Q6IS14 | BRCC3 | EIF5AL1 | 0.582 |
| P46736 | Q14191 | BRCC3 | WRN | 0.582 |
| P46736 | Q8IYT4 | BRCC3 | KATNAL2 | 0.581 |
| P46736 | P55010 | BRCC3 | EIF5 | 0.581 |
| P46736 | P20042 | BRCC3 | EIF2S2 | 0.581 |
| P46736 | Q9Y244 | BRCC3 | POMP | 0.580 |
| P46736 | Q8IVF6 | BRCC3 | ANKRD18A | 0.580 |
| P46736 | Q15646 | BRCC3 | OASL | 0.579 |
| P46736 | Q9C0D5 | BRCC3 | TANC1 | 0.579 |
| P46736 | P54725 | BRCC3 | RAD23A | 0.579 |
| P46736 | A2A2Z9 | BRCC3 | ANKRD18B | 0.577 |
| P46736 | Q99549 | BRCC3 | MPHOSPH8 | 0.577 |
| P46736 | Q9HBL0 | BRCC3 | TNS1 | 0.577 |
| P46736 | Q8NDX5 | BRCC3 | PHC3 | 0.576 |

| | | | | |
|---|---|---|---|---|
| P46736 | Q86XP3 | BRCC3 | DDX42 | 0.576 |
| P46736 | P06401 | BRCC3 | PGR | 0.575 |
| P46736 | P51610 | BRCC3 | HCFC1 | 0.574 |
| P46736 | O75886 | BRCC3 | STAM2 | 0.574 |
| P46736 | Q96MN5 | BRCC3 | TCEANC2 | 0.573 |
| P46736 | P78356 | BRCC3 | PIP4K2B | 0.572 |
| P46736 | Q96B02 | BRCC3 | UBE2W | 0.571 |
| P46736 | P17980 | BRCC3 | PSMC3 | 0.570 |
| P46736 | Q2TAK8 | BRCC3 | PWWP3A | 0.570 |
| P46736 | Q01804 | BRCC3 | OTUD4 | 0.569 |
| P46736 | Q96C11 | BRCC3 | FGGY | 0.569 |
| P46736 | Q86T82 | BRCC3 | USP37 | 0.569 |
| P46736 | P45974 | BRCC3 | USP5 | 0.569 |
| P46736 | Q86Y37 | BRCC3 | CACUL1 | 0.568 |
| P46736 | Q9Y5K6 | BRCC3 | CD2AP | 0.567 |
| P46736 | Q9UPX8 | BRCC3 | SHANK2 | 0.567 |
| P46736 | Q8WVY7 | BRCC3 | UBLCP1 | 0.567 |
| P46736 | Q96DT6 | BRCC3 | ATG4C | 0.567 |
| P46736 | O43815 | BRCC3 | STRN | 0.567 |
| P46736 | Q16763 | BRCC3 | UBE2S | 0.564 |
| P46736 | Q96A44 | BRCC3 | SPSB4 | 0.564 |
| P46736 | Q9NPD8 | BRCC3 | UBE2T | 0.564 |
| P46736 | O43633 | BRCC3 | CHMP2A | 0.562 |
| P46736 | Q15059 | BRCC3 | BRD3 | 0.562 |
| P46736 | Q5JPF3 | BRCC3 | ANKRD36C | 0.561 |
| P46736 | Q7L8S5 | BRCC3 | OTUD6A | 0.561 |
| P46736 | P49427 | BRCC3 | CDC34 | 0.560 |
| P46736 | P38606 | BRCC3 | ATP6V1A | 0.560 |
| P46736 | P28070 | BRCC3 | PSMB4 | 0.560 |
| P46736 | Q9BYE7 | BRCC3 | PCGF6 | 0.559 |
| P46736 | O14980 | BRCC3 | XPO1 | 0.559 |
| P46736 | Q92831 | BRCC3 | KAT2B | 0.558 |
| P46736 | Q8TAF3 | BRCC3 | WDR48 | 0.558 |
| P46736 | Q8IXB1 | BRCC3 | DNAJC10 | 0.555 |
| P46736 | O75602 | BRCC3 | SPAG6 | 0.553 |
| P46736 | P54727 | BRCC3 | RAD23B | 0.552 |
| P46736 | O94826 | BRCC3 | TOMM70 | 0.550 |
| P46736 | O14974 | BRCC3 | PPP1R12A | 0.550 |
| P46736 | Q15386 | BRCC3 | UBE3C | 0.550 |
| P46736 | Q8TBZ3 | BRCC3 | WDR20 | 0.549 |
| P46736 | Q8WZ74 | BRCC3 | CTTNBP2 | 0.548 |
| P46736 | Q70CQ3 | BRCC3 | USP30 | 0.546 |
| P46736 | Q9H0E9 | BRCC3 | BRD8 | 0.545 |
| P46736 | O43513 | BRCC3 | MED7 | 0.545 |
| P46736 | Q58F21 | BRCC3 | BRDT | 0.544 |
| P46736 | P68036 | BRCC3 | UBE2L3 | 0.543 |
| P46736 | Q9UGI0 | BRCC3 | ZRANB1 | 0.543 |
| P46736 | Q9BQD3 | BRCC3 | KXD1 | 0.542 |

| | | | | |
|---|---|---|---|---|
| P46736 | O43172 | BRCC3 | PRPF4 | 0.542 |
| P46736 | Q70EL1 | BRCC3 | USP54 | 0.541 |
| P46736 | O60331 | BRCC3 | PIP5K1C | 0.541 |
| P46736 | O14933 | BRCC3 | UBE2L6 | 0.541 |
| P46736 | P62253 | BRCC3 | UBE2G1 | 0.541 |
| P46736 | Q9NZ32 | BRCC3 | ACTR10 | 0.539 |
| P46736 | O75179 | BRCC3 | ANKRD17 | 0.538 |
| P46736 | Q9UBS4 | BRCC3 | DNAJB11 | 0.538 |
| P46736 | P22570 | BRCC3 | FDXR | 0.536 |
| P46736 | P49459 | BRCC3 | UBE2A | 0.535 |
| P46736 | Q9BZF9 | BRCC3 | UACA | 0.535 |
| P46736 | P82664 | BRCC3 | MRPS10 | 0.534 |
| P46736 | Q12797 | BRCC3 | ASPH | 0.534 |
| P46736 | Q14999 | BRCC3 | CUL7 | 0.534 |
| P46736 | Q12955 | BRCC3 | ANK3 | 0.531 |
| P46736 | P52948 | BRCC3 | NUP98 | 0.530 |
| P46736 | Q92564 | BRCC3 | DCUN1D4 | 0.530 |
| P46736 | Q8N2N9 | BRCC3 | ANKRD36B | 0.529 |
| P46736 | Q8NB14 | BRCC3 | USP38 | 0.528 |
| P46736 | Q9NQI0 | BRCC3 | DDX4 | 0.528 |
| P46736 | Q9NRR5 | BRCC3 | UBQLN4 | 0.526 |
| P46736 | Q9NPI8 | BRCC3 | FANCF | 0.522 |
| P46736 | Q8NB90 | BRCC3 | SPATA5 | 0.522 |
| P46736 | P25440 | BRCC3 | BRD2 | 0.522 |
| P46736 | O94817 | BRCC3 | ATG12 | 0.521 |
| P46736 | Q05639 | BRCC3 | EEF1A2 | 0.520 |
| P46736 | Q9BXF3 | BRCC3 | CECR2 | 0.515 |
| P46736 | P23193 | BRCC3 | TCEA1 | 0.514 |
| P46736 | P62244 | BRCC3 | RPS15A | 0.513 |
| P46736 | O94776 | BRCC3 | MTA2 | 0.512 |
| P46736 | P78317 | BRCC3 | RNF4 | 0.510 |
| P46736 | Q8N283 | BRCC3 | ANKRD35 | 0.509 |
| P46736 | P55072 | BRCC3 | VCP | 0.507 |
| P46736 | Q14C86 | BRCC3 | GAPVD1 | 0.507 |
| P46736 | Q14596 | BRCC3 | NBR1 | 0.505 |
| P46736 | P60510 | BRCC3 | PPP4C | 0.505 |
| P46736 | O96015 | BRCC3 | DNAL4 | 0.505 |
| P46736 | P62979 | BRCC3 | RPS27A | 0.504 |
| P46736 | P25705 | BRCC3 | ATP5F1A | 0.504 |
| P46736 | Q8TEY7 | BRCC3 | USP33 | 0.504 |
| P46736 | O95402 | BRCC3 | MED26 | 0.500 |
| P54252 | P46736 | ATXN3 | BRCC3 | 0.690 |
| P54578 | P46736 | USP14 | BRCC3 | 0.531 |
| P62068 | P46736 | USP46 | BRCC3 | 0.611 |
| Q14694 | P46736 | USP10 | BRCC3 | 0.573 |
| Q15040 | P46736 | JOSD1 | BRCC3 | 0.628 |
| Q504Q3 | P46736 | PAN2 | BRCC3 | 0.638 |
| Q53GS9 | P46736 | USP39 | BRCC3 | 0.627 |

| | | | | |
|---|---|---|---|---|
| Q5VVJ2 | P46736 | MYSM1 | BRCC3 | 0.555 |
| Q70CQ1 | P46736 | USP49 | BRCC3 | 0.627 |
| Q70CQ3 | P46736 | USP30 | BRCC3 | 0.683 |
| Q70EL2 | P46736 | USP45 | BRCC3 | 0.657 |
| Q70EL3 | P46736 | USP50 | BRCC3 | 0.613 |
| Q70EL4 | P46736 | USP43 | BRCC3 | 0.548 |
| Q8NB14 | P46736 | USP38 | BRCC3 | 0.503 |
| Q8TEY7 | P46736 | USP33 | BRCC3 | 0.695 |
| Q92560 | P46736 | BAP1 | BRCC3 | 0.623 |
| Q92905 | P46736 | COPS5 | BRCC3 | 0.536 |
| Q96BN8 | P46736 | OTULIN | BRCC3 | 0.560 |
| Q9BXU7 | P46736 | USP26 | BRCC3 | 0.533 |
| Q9H0E7 | P46736 | USP44 | BRCC3 | 0.631 |
| Q9H3M9 | P46736 | ATXN3L | BRCC3 | 0.694 |
| Q9P275 | P46736 | USP36 | BRCC3 | 0.633 |
| Q9P2H5 | P46736 | USP35 | BRCC3 | 0.616 |
| Q9UMW8 | P46736 | USP18 | BRCC3 | 0.617 |
| Q9UPT9 | P46736 | USP22 | BRCC3 | 0.680 |
| Q9Y2K6 | P46736 | USP20 | BRCC3 | 0.525 |
| Q9Y5K5 | P46736 | UCHL5 | BRCC3 | 0.630 |
| Q9Y5T5 | P46736 | USP16 | BRCC3 | 0.611 |

| | | | | |
|---|---|---|---|---|
| Q96K76 | Q9HAW4 | USP47 | CLSPN | 0.992 |
| Q96K76 | P04637 | USP47 | TP53 | 0.967 |
| Q96K76 | Q15154 | USP47 | PCM1 | 0.965 |
| Q96K76 | Q96RL1 | USP47 | UIMC1 | 0.962 |
| Q96K76 | O15151 | USP47 | MDM4 | 0.962 |
| Q96K76 | Q00987 | USP47 | MDM2 | 0.962 |
| Q96K76 | P35659 | USP47 | DEK | 0.956 |
| Q96K76 | P30304 | USP47 | CDC25A | 0.952 |
| Q96K76 | P32314 | USP47 | FOXN2 | 0.949 |
| Q96K76 | P62877 | USP47 | RBX1 | 0.939 |
| Q96K76 | Q6IEE8 | USP47 | SLFN12L | 0.937 |
| Q96K76 | Q9BXW9 | USP47 | FANCD2 | 0.936 |
| Q96K76 | Q9BY41 | USP47 | HDAC8 | 0.935 |
| Q96K76 | Q68D06 | USP47 | SLFN13 | 0.933 |
| Q96K76 | Q08AF3 | USP47 | SLFN5 | 0.930 |
| Q96K76 | Q7Z7L1 | USP47 | SLFN11 | 0.925 |
| Q96K76 | Q13127 | USP47 | REST | 0.923 |
| Q96K76 | Q2YD98 | USP47 | UVSSA | 0.916 |
| Q96K76 | Q9UKA2 | USP47 | FBXL4 | 0.907 |
| Q96K76 | Q13309 | USP47 | SKP2 | 0.904 |
| Q96K76 | Q8IWA4 | USP47 | MFN1 | 0.904 |
| Q96K76 | O95140 | USP47 | MFN2 | 0.904 |
| Q96K76 | Q56UN5 | USP47 | MAP3K19 | 0.903 |
| Q96K76 | Q8IVP5 | USP47 | FUNDC1 | 0.903 |
| Q96K76 | Q9UKV0 | USP47 | HDAC9 | 0.899 |
| Q96K76 | Q96BM1 | USP47 | ANKRD9 | 0.898 |

| | | | | |
|---|---|---|---|---|
| Q96K76 | Q92889 | USP47 | ERCC4 | 0.897 |
| Q96K76 | Q9UBN7 | USP47 | HDAC6 | 0.894 |
| Q96K76 | Q8IYW5 | USP47 | RNF168 | 0.893 |
| Q96K76 | Q01167 | USP47 | FOXK2 | 0.890 |
| Q96K76 | Q9UBW7 | USP47 | ZMYM2 | 0.890 |
| Q96K76 | P85037 | USP47 | FOXK1 | 0.889 |
| Q96K76 | Q96BN8 | USP47 | OTULIN | 0.887 |
| Q96K76 | Q14146 | USP47 | URB2 | 0.880 |
| Q96K76 | Q9UKT4 | USP47 | FBXO5 | 0.876 |
| Q96K76 | Q9H0D2 | USP47 | ZNF541 | 0.876 |
| Q96K76 | Q6N043 | USP47 | ZNF280D | 0.874 |
| Q96K76 | Q9Y6B7 | USP47 | AP4B1 | 0.873 |
| Q96K76 | P15056 | USP47 | BRAF | 0.871 |
| Q96K76 | O43583 | USP47 | DENR | 0.870 |
| Q96K76 | Q8WTU0 | USP47 | DDI1 | 0.869 |
| Q96K76 | Q96F46 | USP47 | IL17RA | 0.869 |
| Q96K76 | P25963 | USP47 | NFKBIA | 0.866 |
| Q96K76 | Q9UBF6 | USP47 | RNF7 | 0.858 |
| Q96K76 | Q9NYY3 | USP47 | PLK2 | 0.858 |
| Q96K76 | P37840 | USP47 | SNCA | 0.858 |
| Q96K76 | Q6VAB6 | USP47 | KSR2 | 0.857 |
| Q96K76 | Q9UL58 | USP47 | ZNF215 | 0.857 |
| Q96K76 | P06213 | USP47 | INSR | 0.855 |
| Q96K76 | Q6FI81 | USP47 | CIAPIN1 | 0.855 |
| Q96K76 | Q9BQG2 | USP47 | NUDT12 | 0.853 |
| Q96K76 | P14373 | USP47 | TRIM27 | 0.849 |
| Q96K76 | Q9P2D0 | USP47 | IBTK | 0.848 |
| Q96K76 | Q7Z6E9 | USP47 | RBBP6 | 0.845 |
| Q96K76 | Q9Y4E5 | USP47 | ZNF451 | 0.845 |
| Q96K76 | Q17RY0 | USP47 | CPEB4 | 0.844 |
| Q96K76 | Q6P996 | USP47 | PDXDC1 | 0.842 |
| Q96K76 | Q8N4B4 | USP47 | FBXO39 | 0.841 |
| Q96K76 | Q8WXK4 | USP47 | ASB12 | 0.840 |
| Q96K76 | Q04609 | USP47 | FOLH1 | 0.838 |
| Q96K76 | Q9Y294 | USP47 | ASF1A | 0.837 |
| Q96K76 | Q8NDW4 | USP47 | ZNF248 | 0.835 |
| Q96K76 | O00203 | USP47 | AP3B1 | 0.834 |
| Q96K76 | P19838 | USP47 | NFKB1 | 0.834 |
| Q96K76 | Q8IYD8 | USP47 | FANCM | 0.834 |
| Q96K76 | P29459 | USP47 | IL12A | 0.834 |
| Q96K76 | Q9UJT9 | USP47 | FBXL7 | 0.833 |
| Q96K76 | P53350 | USP47 | PLK1 | 0.832 |
| Q96K76 | Q9UKC9 | USP47 | FBXL2 | 0.831 |
| Q96K76 | Q6PIY7 | USP47 | TENT2 | 0.830 |
| Q96K76 | Q86T24 | USP47 | ZBTB33 | 0.829 |
| Q96K76 | Q99558 | USP47 | MAP3K14 | 0.828 |
| Q96K76 | P17014 | USP47 | ZNF12 | 0.828 |
| Q96K76 | P60484 | USP47 | PTEN | 0.827 |

| | | | | |
|---|---|---|---|---|
| Q96K76 | Q99728 | USP47 | BARD1 | 0.824 |
| Q96K76 | P20749 | USP47 | BCL3 | 0.823 |
| Q96K76 | Q9NZJ5 | USP47 | EIF2AK3 | 0.823 |
| Q96K76 | Q13439 | USP47 | GOLGA4 | 0.821 |
| Q96K76 | Q9H165 | USP47 | BCL11A | 0.817 |
| Q96K76 | Q9Y575 | USP47 | ASB3 | 0.817 |
| Q96K76 | P50542 | USP47 | PEX5 | 0.816 |
| Q96K76 | P04049 | USP47 | RAF1 | 0.814 |
| Q96K76 | Q9H469 | USP47 | FBXL15 | 0.814 |
| Q96K76 | Q13367 | USP47 | AP3B2 | 0.814 |
| Q96K76 | Q86YH2 | USP47 | ZNF280B | 0.813 |
| Q96K76 | Q17RQ9 | USP47 | NKPD1 | 0.813 |
| Q96K76 | P57078 | USP47 | RIPK4 | 0.812 |
| Q96K76 | P39748 | USP47 | FEN1 | 0.812 |
| Q96K76 | Q86Z02 | USP47 | HIPK1 | 0.812 |
| Q96K76 | Q92499 | USP47 | DDX1 | 0.810 |
| Q96K76 | O00429 | USP47 | DNM1L | 0.810 |
| Q96K76 | P54274 | USP47 | TERF1 | 0.810 |
| Q96K76 | Q96EB6 | USP47 | SIRT1 | 0.810 |
| Q96K76 | O75155 | USP47 | CAND2 | 0.809 |
| Q96K76 | Q9NZS9 | USP47 | BFAR | 0.809 |
| Q96K76 | Q99759 | USP47 | MAP3K3 | 0.809 |
| Q96K76 | Q7Z3U7 | USP47 | MON2 | 0.808 |
| Q96K76 | Q9Y3I1 | USP47 | FBXO7 | 0.807 |
| Q96K76 | Q9UK97 | USP47 | FBXO9 | 0.807 |
| Q96K76 | Q96Q07 | USP47 | BTBD9 | 0.805 |
| Q96K76 | Q9Y6H5 | USP47 | SNCAIP | 0.804 |
| Q96K76 | Q13233 | USP47 | MAP3K1 | 0.804 |
| Q96K76 | Q16690 | USP47 | DUSP5 | 0.804 |
| Q96K76 | Q96HA7 | USP47 | TONSL | 0.802 |
| Q96K76 | Q9BT40 | USP47 | INPP5K | 0.800 |
| Q96K76 | Q05193 | USP47 | DNM1 | 0.800 |
| Q96K76 | Q9H0K1 | USP47 | SIK2 | 0.800 |
| Q96K76 | Q9H4P4 | USP47 | RNF41 | 0.799 |
| Q96K76 | O95785 | USP47 | WIZ | 0.798 |
| Q96K76 | Q96JP5 | USP47 | ZFP91 | 0.797 |
| Q96K76 | Q8NI38 | USP47 | NFKBID | 0.794 |
| Q96K76 | Q9NQC1 | USP47 | JADE2 | 0.793 |
| Q96K76 | Q8WXK1 | USP47 | ASB15 | 0.791 |
| Q96K76 | O94916 | USP47 | NFAT5 | 0.790 |
| Q96K76 | P10398 | USP47 | ARAF | 0.789 |
| Q96K76 | Q8WXK3 | USP47 | ASB13 | 0.787 |
| Q96K76 | Q15653 | USP47 | NFKBIB | 0.786 |
| Q96K76 | Q8WWX0 | USP47 | ASB5 | 0.786 |
| Q96K76 | Q9H765 | USP47 | ASB8 | 0.786 |
| Q96K76 | Q9ULH0 | USP47 | KIDINS220 | 0.785 |
| Q96K76 | Q8IYB4 | USP47 | PEX5L | 0.785 |
| Q96K76 | Q9Y468 | USP47 | L3MBTL1 | 0.784 |

| | | | | |
|---|---|---|---|---|
| Q96K76 | Q495M9 | USP47 | USH1G | 0.781 |
| Q96K76 | P04908 | USP47 | H2AC4 | 0.779 |
| Q96K76 | Q8WXI3 | USP47 | ASB10 | 0.778 |
| Q96K76 | Q969U6 | USP47 | FBXW5 | 0.777 |
| Q96K76 | Q9NR09 | USP47 | BIRC6 | 0.775 |
| Q96K76 | Q13177 | USP47 | PAK2 | 0.775 |
| Q96K76 | Q16778 | USP47 | H2BC21 | 0.774 |
| Q96K76 | B4E2M5 | USP47 | ANKRD66 | 0.773 |
| Q96K76 | O43781 | USP47 | DYRK3 | 0.772 |
| Q96K76 | Q7Z713 | USP47 | ANKRD37 | 0.771 |
| Q96K76 | Q9Y3Q0 | USP47 | NAALAD2 | 0.770 |
| Q96K76 | Q53RE8 | USP47 | ANKRD39 | 0.770 |
| Q96K76 | Q13627 | USP47 | DYRK1A | 0.770 |
| Q96K76 | Q9HCE7 | USP47 | SMURF1 | 0.769 |
| Q96K76 | Q9UKT5 | USP47 | FBXO4 | 0.769 |
| Q96K76 | Q8NE63 | USP47 | HIPK4 | 0.765 |
| Q96K76 | Q7Z3V4 | USP47 | UBE3B | 0.765 |
| Q96K76 | Q9HAU4 | USP47 | SMURF2 | 0.764 |
| Q96K76 | Q8NB46 | USP47 | ANKRD52 | 0.763 |
| Q96K76 | P50570 | USP47 | DNM2 | 0.763 |
| Q96K76 | Q9Y463 | USP47 | DYRK1B | 0.763 |
| Q96K76 | P42773 | USP47 | CDKN2C | 0.763 |
| Q96K76 | Q9UKA1 | USP47 | FBXL5 | 0.761 |
| Q96K76 | Q9UII4 | USP47 | HERC5 | 0.761 |
| Q96K76 | Q9Y2U5 | USP47 | MAP3K2 | 0.761 |
| Q96K76 | Q9NVX7 | USP47 | KBTBD4 | 0.761 |
| Q96K76 | P08069 | USP47 | IGF1R | 0.760 |
| Q96K76 | Q8WXH4 | USP47 | ASB11 | 0.760 |
| Q96K76 | Q5VYY1 | USP47 | ANKRD22 | 0.760 |
| Q96K76 | Q96DX5 | USP47 | ASB9 | 0.760 |
| Q96K76 | P30291 | USP47 | WEE1 | 0.759 |
| Q96K76 | Q9BYH8 | USP47 | NFKBIZ | 0.759 |
| Q96K76 | Q9H2X6 | USP47 | HIPK2 | 0.758 |
| Q96K76 | Q9H9E1 | USP47 | ANKRA2 | 0.758 |
| Q96K76 | P63010 | USP47 | AP2B1 | 0.757 |
| Q96K76 | P49915 | USP47 | GMPS | 0.756 |
| Q96K76 | Q15034 | USP47 | HERC3 | 0.755 |
| Q96K76 | Q9Y2K2 | USP47 | SIK3 | 0.755 |
| Q96K76 | Q9H422 | USP47 | HIPK3 | 0.754 |
| Q96K76 | Q9Y574 | USP47 | ASB4 | 0.753 |
| Q96K76 | Q504Q3 | USP47 | PAN2 | 0.752 |
| Q96K76 | O00221 | USP47 | NFKBIE | 0.752 |
| Q96K76 | A6NK59 | USP47 | ASB14 | 0.751 |
| Q96K76 | Q8NEW0 | USP47 | SLC30A7 | 0.751 |
| Q96K76 | Q12888 | USP47 | TP53BP1 | 0.750 |
| Q96K76 | Q9Y576 | USP47 | ASB1 | 0.750 |
| Q96K76 | Q8IVU3 | USP47 | HERC6 | 0.750 |
| Q96K76 | P51965 | USP47 | UBE2E1 | 0.750 |

| | | | | |
|---|---|---|---|---|
| Q96K76 | Q6XR72 | USP47 | SLC30A10 | 0.749 |
| Q96K76 | Q9NWX5 | USP47 | ASB6 | 0.749 |
| Q96K76 | Q6P3X3 | USP47 | TTC27 | 0.749 |
| Q96K76 | Q5H9U9 | USP47 | DDX60L | 0.749 |
| Q96K76 | Q4G0W2 | USP47 | DUSP28 | 0.748 |
| Q96K76 | Q92618 | USP47 | ZNF516 | 0.748 |
| Q96K76 | Q9BSK4 | USP47 | FEM1A | 0.747 |
| Q96K76 | Q8TC84 | USP47 | FANK1 | 0.747 |
| Q96K76 | P42772 | USP47 | CDKN2B | 0.746 |
| Q96K76 | P55273 | USP47 | CDKN2D | 0.745 |
| Q96K76 | Q6NXT4 | USP47 | SLC30A6 | 0.744 |
| Q96K76 | P17030 | USP47 | ZNF25 | 0.743 |
| Q96K76 | O95202 | USP47 | LETM1 | 0.742 |
| Q96K76 | O15033 | USP47 | AREL1 | 0.742 |
| Q96K76 | Q9HAU0 | USP47 | PLEKHA5 | 0.742 |
| Q96K76 | Q16777 | USP47 | H2AC20 | 0.741 |
| Q96K76 | Q86VP6 | USP47 | CAND1 | 0.741 |
| Q96K76 | Q8TDJ6 | USP47 | DMXL2 | 0.739 |
| Q96K76 | Q86W74 | USP47 | ANKRD46 | 0.738 |
| Q96K76 | Q9UQ16 | USP47 | DNM3 | 0.738 |
| Q96K76 | P42771 | USP47 | CDKN2A | 0.737 |
| Q96K76 | Q8N9B4 | USP47 | ANKRD42 | 0.737 |
| Q96K76 | Q8N8V4 | USP47 | ANKS4B | 0.736 |
| Q96K76 | Q9BQI6 | USP47 | SLF1 | 0.736 |
| Q96K76 | Q9GZV5 | USP47 | WWTR1 | 0.734 |
| Q96K76 | Q6KC79 | USP47 | NIPBL | 0.733 |
| Q96K76 | P54868 | USP47 | HMGCS2 | 0.732 |
| Q96K76 | Q01581 | USP47 | HMGCS1 | 0.732 |
| Q96K76 | Q6IE81 | USP47 | JADE1 | 0.731 |
| Q96K76 | P57059 | USP47 | SIK1 | 0.731 |
| Q96K76 | A0A0B4J2F2 | USP47 | SIK1B | 0.731 |
| Q96K76 | Q969H0 | USP47 | FBXW7 | 0.731 |
| Q96K76 | Q10567 | USP47 | AP1B1 | 0.730 |
| Q96K76 | Q9NVR0 | USP47 | KLHL11 | 0.726 |
| Q96K76 | Q8IYU2 | USP47 | HACE1 | 0.725 |
| Q96K76 | Q96KE9 | USP47 | BTBD6 | 0.724 |
| Q96K76 | Q8N961 | USP47 | ABTB2 | 0.723 |
| Q96K76 | Q13432 | USP47 | UNC119 | 0.723 |
| Q96K76 | Q68DC2 | USP47 | ANKS6 | 0.722 |
| Q96K76 | Q5T447 | USP47 | HECTD3 | 0.722 |
| Q96K76 | Q6NXT1 | USP47 | ANKRD54 | 0.721 |
| Q96K76 | Q7Z6Z7 | USP47 | HUWE1 | 0.721 |
| Q96K76 | P46937 | USP47 | YAP1 | 0.720 |
| Q96K76 | Q8TAD4 | USP47 | SLC30A5 | 0.720 |
| Q96K76 | Q9BV47 | USP47 | DUSP26 | 0.719 |
| Q96K76 | P19525 | USP47 | EIF2AK2 | 0.716 |
| Q96K76 | Q96JP0 | USP47 | FEM1C | 0.715 |
| Q96K76 | O94952 | USP47 | FBXO21 | 0.714 |

| | | | | |
|---|---|---|---|---|
| Q96K76 | Q15327 | USP47 | ANKRD1 | 0.713 |
| Q96K76 | Q5D1E8 | USP47 | ZC3H12A | 0.712 |
| Q96K76 | A6QL63 | USP47 | BTBD11 | 0.711 |
| Q96K76 | Q8IV38 | USP47 | ANKMY2 | 0.707 |
| Q96K76 | Q92882 | USP47 | OSTF1 | 0.707 |
| Q96K76 | Q9Y2Z4 | USP47 | YARS2 | 0.707 |
| Q96K76 | O14863 | USP47 | SLC30A4 | 0.706 |
| Q96K76 | Q5XX13 | USP47 | FBXW10 | 0.706 |
| Q96K76 | O14593 | USP47 | RFXANK | 0.705 |
| Q96K76 | Q9NSD9 | USP47 | FARSB | 0.705 |
| Q96K76 | Q6ZVH7 | USP47 | ESPNL | 0.704 |
| Q96K76 | O95817 | USP47 | BAG3 | 0.703 |
| Q96K76 | Q9Y2F9 | USP47 | BTBD3 | 0.703 |
| Q96K76 | Q9P2R3 | USP47 | ANKFY1 | 0.700330859 |
| Q96K76 | Q7L9B9 | USP47 | EEPD1 | 0.699959767 |
| Q96K76 | Q96JH7 | USP47 | VCPIP1 | 0.699601691 |
| Q96K76 | Q5XUX1 | USP47 | FBXW9 | 0.699457411 |
| Q96K76 | O95170 | USP47 | CDRT1 | 0.699056721 |
| Q96K76 | Q8IVT5 | USP47 | KSR1 | 0.697699121 |
| Q96K76 | Q969K4 | USP47 | ABTB1 | 0.697423317 |
| Q96K76 | Q15477 | USP47 | SKIV2L | 0.696663212 |
| Q96K76 | Q8N6D5 | USP47 | ANKRD29 | 0.696414437 |
| Q96K76 | P51522 | USP47 | ZNF83 | 0.695414973 |
| Q96K76 | O43390 | USP47 | HNRNPR | 0.694740493 |
| Q96K76 | Q9Y283 | USP47 | INVS | 0.694164913 |
| Q96K76 | Q5GLZ8 | USP47 | HERC4 | 0.693750431 |
| Q96K76 | Q00653 | USP47 | NFKB2 | 0.69331377 |
| Q96K76 | Q5U5R9 | USP47 | HECTD2 | 0.692654686 |
| Q96K76 | Q6ZVZ8 | USP47 | ASB18 | 0.692643844 |
| Q96K76 | Q8IWZ3 | USP47 | ANKHD1 | 0.692453773 |
| Q96K76 | O60907 | USP47 | TBL1X | 0.692424449 |
| Q96K76 | O60313 | USP47 | OPA1 | 0.689200434 |
| Q96K76 | P48436 | USP47 | SOX9 | 0.688755672 |
| Q96K76 | Q14678 | USP47 | KANK1 | 0.686704861 |
| Q96K76 | Q6IQ16 | USP47 | SPOPL | 0.686242948 |
| Q96K76 | Q92527 | USP47 | ANKRD7 | 0.685939264 |
| Q96K76 | Q8TCX5 | USP47 | RHPN1 | 0.685058772 |
| Q96K76 | Q17RB8 | USP47 | LONRF1 | 0.684577259 |
| Q96K76 | Q9ULT8 | USP47 | HECTD1 | 0.683576237 |
| Q96K76 | Q9NVW2 | USP47 | RLIM | 0.683528687 |
| Q96K76 | Q99592 | USP47 | ZBTB18 | 0.683072544 |
| Q96K76 | Q96S82 | USP47 | UBL7 | 0.681800098 |
| Q96K76 | O15084 | USP47 | ANKRD28 | 0.681140943 |
| Q96K76 | Q9Y485 | USP47 | DMXL1 | 0.680449028 |
| Q96K76 | Q9HAZ1 | USP47 | CLK4 | 0.679172968 |
| Q96K76 | Q76N89 | USP47 | HECW1 | 0.678799464 |
| Q96K76 | Q96NS5 | USP47 | ASB16 | 0.678090815 |
| Q96K76 | Q9NZL4 | USP47 | HSPBP1 | 0.677864947 |

| | | | | |
|---|---|---|---|---|
| Q96K76 | O76064 | USP47 | RNF8 | 0.677659121 |
| Q96K76 | Q12834 | USP47 | CDC20 | 0.677290114 |
| Q96K76 | Q9BXX2 | USP47 | ANKRD30B | 0.675916887 |
| Q96K76 | P27708 | USP47 | CAD | 0.675283821 |
| Q96K76 | P49643 | USP47 | PRIM2 | 0.674088385 |
| Q96K76 | Q6P4R8 | USP47 | NFRKB | 0.673445496 |
| Q96K76 | Q6ZW76 | USP47 | ANKS3 | 0.67291644 |
| Q96K76 | Q9Y6M5 | USP47 | SLC30A1 | 0.672876599 |
| Q96K76 | Q9UKT8 | USP47 | FBXW2 | 0.672662161 |
| Q96K76 | Q86YT6 | USP47 | MIB1 | 0.668859669 |
| Q96K76 | Q9NXR5 | USP47 | ANKRD10 | 0.66831265 |
| Q96K76 | Q8NCN4 | USP47 | RNF169 | 0.667939272 |
| Q96K76 | Q96Q27 | USP47 | ASB2 | 0.665813087 |
| Q96K76 | Q3KP44 | USP47 | ANKRD55 | 0.665577354 |
| Q96K76 | Q9GZV1 | USP47 | ANKRD2 | 0.665363773 |
| Q96K76 | Q86SG2 | USP47 | ANKRD23 | 0.665040291 |
| Q96K76 | Q9C0D0 | USP47 | PHACTR1 | 0.662670529 |
| Q96K76 | Q16890 | USP47 | TPD52L1 | 0.662406581 |
| Q96K76 | O94759 | USP47 | TRPM2 | 0.661428593 |
| Q96K76 | Q9HCD6 | USP47 | TANC2 | 0.661344142 |
| Q96K76 | Q5XPI4 | USP47 | RNF123 | 0.661291988 |
| Q96K76 | Q9H6Y5 | USP47 | MAGIX | 0.660054554 |
| Q96K76 | O75167 | USP47 | PHACTR2 | 0.659718557 |
| Q96K76 | Q96AX9 | USP47 | MIB2 | 0.65962275 |
| Q96K76 | Q6P6B7 | USP47 | ANKRD16 | 0.659068483 |
| Q96K76 | Q309B1 | USP47 | TRIM16L | 0.658832109 |
| Q96K76 | Q96PU5 | USP47 | NEDD4L | 0.65760268 |
| Q96K76 | O43861 | USP47 | ATP9B | 0.657347101 |
| Q96K76 | Q14669 | USP47 | TRIP12 | 0.656407866 |
| Q96K76 | P53618 | USP47 | COPB1 | 0.656082472 |
| Q96K76 | P55081 | USP47 | MFAP1 | 0.65606281 |
| Q96K76 | Q8NHS4 | USP47 | CLHC1 | 0.655551686 |
| Q96K76 | Q9C0D7 | USP47 | ZC3H12C | 0.652412326 |
| Q96K76 | Q8IY21 | USP47 | DDX60 | 0.652004795 |
| Q96K76 | Q15744 | USP47 | CEBPE | 0.65121643 |
| Q96K76 | Q05086 | USP47 | UBE3A | 0.649820172 |
| Q96K76 | Q6PIW4 | USP47 | FIGNL1 | 0.649520883 |
| Q96K76 | Q96J02 | USP47 | ITCH | 0.649192001 |
| Q96K76 | P46934 | USP47 | NEDD4 | 0.648402341 |
| Q96K76 | Q9UNP9 | USP47 | PPIE | 0.646640641 |
| Q96K76 | Q9ULE0 | USP47 | WWC3 | 0.646619885 |
| Q96K76 | Q92630 | USP47 | DYRK2 | 0.645506143 |
| Q96K76 | Q9P2P5 | USP47 | HECW2 | 0.645316183 |
| Q96K76 | O00308 | USP47 | WWP2 | 0.644447274 |
| Q96K76 | Q8NFZ0 | USP47 | FBH1 | 0.644016779 |
| Q96K76 | Q9H832 | USP47 | UBE2Z | 0.643799975 |
| Q96K76 | Q6AWC2 | USP47 | WWC2 | 0.643385115 |
| Q96K76 | O43933 | USP47 | PEX1 | 0.642960512 |

| | | | | |
|---|---|---|---|---|
| Q96K76 | Q07890 | USP47 | SOS2 | 0.641018482 |
| Q96K76 | Q70EK8 | USP47 | USP53 | 0.640376704 |
| Q96K76 | Q05823 | USP47 | RNASEL | 0.638625628 |
| Q96K76 | B1AK53 | USP47 | ESPN | 0.636628745 |
| Q96K76 | Q5HYM0 | USP47 | ZC3H12B | 0.63563406 |
| Q96K76 | P53567 | USP47 | CEBPG | 0.633921786 |
| Q96K76 | O95071 | USP47 | UBR5 | 0.630325992 |
| Q96K76 | A7E2S9 | USP47 | ANKRD30BL | 0.629294148 |
| Q96K76 | Q9Y2G4 | USP47 | ANKRD6 | 0.629024324 |
| Q96K76 | Q6PGP7 | USP47 | TTC37 | 0.628648061 |
| Q96K76 | Q96L34 | USP47 | MARK4 | 0.628038414 |
| Q96K76 | Q96LR5 | USP47 | UBE2E2 | 0.628000747 |
| Q96K76 | P11441 | USP47 | UBL4A | 0.626997911 |
| Q96K76 | Q8WWR9 | USP47 | PPDPFL | 0.62647969 |
| Q96K76 | Q86UL8 | USP47 | MAGI2 | 0.625322161 |
| Q96K76 | Q8WWH4 | USP47 | ASZ1 | 0.624380109 |
| Q96K76 | P20592 | USP47 | MX2 | 0.623783946 |
| Q96K76 | P11717 | USP47 | IGF2R | 0.620531503 |
| Q96K76 | Q92905 | USP47 | COPS5 | 0.620394012 |
| Q96K76 | Q9HC77 | USP47 | CENPJ | 0.618929394 |
| Q96K76 | P31327 | USP47 | CPS1 | 0.617869695 |
| Q96K76 | Q5T4S7 | USP47 | UBR4 | 0.616771885 |
| Q96K76 | Q15075 | USP47 | EEA1 | 0.614932469 |
| Q96K76 | P52735 | USP47 | VAV2 | 0.614734829 |
| Q96K76 | Q9BSC4 | USP47 | NOL10 | 0.614263697 |
| Q96K76 | Q9C0D5 | USP47 | TANC1 | 0.613855233 |
| Q96K76 | P49716 | USP47 | CEBPD | 0.613318753 |
| Q96K76 | Q9BVG8 | USP47 | KIFC3 | 0.612787878 |
| Q96K76 | A6QL64 | USP47 | ANKRD36 | 0.612670696 |
| Q96K76 | Q6PCT2 | USP47 | FBXL19 | 0.611351843 |
| Q96K76 | P27448 | USP47 | MARK3 | 0.607586353 |
| Q96K76 | A6NC57 | USP47 | ANKRD62 | 0.607392006 |
| Q96K76 | Q9UK73 | USP47 | FEM1B | 0.605918328 |
| Q96K76 | Q4UJ75 | USP47 | ANKRD20A4 | 0.605376829 |
| Q96K76 | Q969T4 | USP47 | UBE2E3 | 0.604701567 |
| Q96K76 | Q15436 | USP47 | SEC23A | 0.604451196 |
| Q96K76 | Q92995 | USP47 | USP13 | 0.602975426 |
| Q96K76 | P61009 | USP47 | SPCS3 | 0.602521126 |
| Q96K76 | Q9UPX8 | USP47 | SHANK2 | 0.60211151 |
| Q96K76 | Q6TFL4 | USP47 | KLHL24 | 0.601773954 |
| Q96K76 | Q9H0C5 | USP47 | BTBD1 | 0.6016044 |
| Q96K76 | P21580 | USP47 | TNFAIP3 | 0.601141063 |
| Q96K76 | Q8N3Y1 | USP47 | FBXW8 | 0.600952638 |
| Q96K76 | Q9H0M0 | USP47 | WWP1 | 0.600677812 |
| Q96K76 | Q8IWU4 | USP47 | SLC30A8 | 0.600219632 |
| Q96K76 | Q9Y6D6 | USP47 | ARFGEF1 | 0.600068213 |
| Q96K76 | Q15386 | USP47 | UBE3C | 0.600051834 |
| Q96K76 | Q496Y0 | USP47 | LONRF3 | 0.599879316 |

| | | | | |
|---|---|---|---|---|
| Q96K76 | Q9HBJ7 | USP47 | USP29 | 0.598930123 |
| Q96K76 | Q9UQR1 | USP47 | ZNF148 | 0.597935936 |
| Q96K76 | P52739 | USP47 | ZNF131 | 0.596938446 |
| Q96K76 | Q5TYW2 | USP47 | ANKRD20A1 | 0.596101335 |
| Q96K76 | Q14999 | USP47 | CUL7 | 0.595926784 |
| Q96K76 | Q5VUR7 | USP47 | ANKRD20A3I | 0.593990466 |
| Q96K76 | P55786 | USP47 | NPEPPS | 0.592907418 |
| Q96K76 | P49715 | USP47 | CEBPA | 0.591214104 |
| Q96K76 | P20936 | USP47 | RASA1 | 0.588433455 |
| Q96K76 | Q7KZI7 | USP47 | MARK2 | 0.586562035 |
| Q96K76 | Q5SQ80 | USP47 | ANKRD20A2I | 0.586303922 |
| Q96K76 | O00189 | USP47 | AP4M1 | 0.586210782 |
| Q96K76 | P53621 | USP47 | COPA | 0.585436577 |
| Q96K76 | Q8WYQ9 | USP47 | ZCCHC14 | 0.585196811 |
| Q96K76 | Q5T655 | USP47 | CFAP58 | 0.584634846 |
| Q96K76 | O95630 | USP47 | STAMBP | 0.584446955 |
| Q96K76 | Q5TGS1 | USP47 | HES3 | 0.583678819 |
| Q96K76 | Q9UBP4 | USP47 | DKK3 | 0.582461927 |
| Q96K76 | Q8IYW4 | USP47 | ENTHD1 | 0.581725427 |
| Q96K76 | Q9ULJ7 | USP47 | ANKRD50 | 0.580208307 |
| Q96K76 | Q9C0B9 | USP47 | ZCCHC2 | 0.580200208 |
| Q96K76 | Q15437 | USP47 | SEC23B | 0.579907825 |
| Q96K76 | Q8N7F7 | USP47 | UBL4B | 0.578353548 |
| Q96K76 | Q5HY92 | USP47 | FIGN | 0.577636852 |
| Q96K76 | P49792 | USP47 | RANBP2 | 0.576069841 |
| Q96K76 | O75592 | USP47 | MYCBP2 | 0.574917057 |
| Q96K76 | O43567 | USP47 | RNF13 | 0.574297888 |
| Q96K76 | Q8IVE3 | USP47 | PLEKHH2 | 0.574287868 |
| Q96K76 | P49759 | USP47 | CLK1 | 0.573643437 |
| Q96K76 | Q9NR20 | USP47 | DYRK4 | 0.573390848 |
| Q96K76 | Q9P0J7 | USP47 | KCMF1 | 0.57282336 |
| Q96K76 | Q9H201 | USP47 | EPN3 | 0.571837441 |
| Q96K76 | O75179 | USP47 | ANKRD17 | 0.571611088 |
| Q96K76 | Q9UPU5 | USP47 | USP24 | 0.570581322 |
| Q96K76 | O95208 | USP47 | EPN2 | 0.570471918 |
| Q96K76 | Q05481 | USP47 | ZNF91 | 0.565313893 |
| Q96K76 | Q9Y566 | USP47 | SHANK1 | 0.564760072 |
| Q96K76 | Q8IVF6 | USP47 | ANKRD18A | 0.564193324 |
| Q96K76 | O15067 | USP47 | PFAS | 0.562684302 |
| Q96K76 | P17676 | USP47 | CEBPB | 0.561912213 |
| Q96K76 | P61086 | USP47 | UBE2K | 0.559521507 |
| Q96K76 | Q00610 | USP47 | CLTC | 0.558453111 |
| Q96K76 | Q13501 | USP47 | SQSTM1 | 0.55734652 |
| Q96K76 | Q96TA2 | USP47 | YME1L1 | 0.555779775 |
| Q96K76 | P53675 | USP47 | CLTCL1 | 0.55443751 |
| Q96K76 | Q8NCL8 | USP47 | TMEM116 | 0.554328356 |
| Q96K76 | Q99726 | USP47 | SLC30A3 | 0.554243333 |
| Q96K76 | P49760 | USP47 | CLK2 | 0.554062595 |

| | | | | |
|---|---|---|---|---|
| Q96K76 | Q99986 | USP47 | VRK1 | 0.554022569 |
| Q96K76 | Q15645 | USP47 | TRIP13 | 0.552903194 |
| Q96K76 | Q8IX03 | USP47 | WWC1 | 0.552648766 |
| Q96K76 | A2A2Z9 | USP47 | ANKRD18B | 0.551907491 |
| Q96K76 | Q70EL1 | USP47 | USP54 | 0.551422353 |
| Q96K76 | Q9BXX3 | USP47 | ANKRD30A | 0.547273644 |
| Q96K76 | Q5JPF3 | USP47 | ANKRD36C | 0.545956385 |
| Q96K76 | O14974 | USP47 | PPP1R12A | 0.544399103 |
| Q96K76 | Q96N64 | USP47 | PWWP2A | 0.543936171 |
| Q96K76 | P22670 | USP47 | RFX1 | 0.54308591 |
| Q96K76 | O43791 | USP47 | SPOP | 0.540898131 |
| Q96K76 | Q9C0C9 | USP47 | UBE2O | 0.537866799 |
| Q96K76 | Q9Y6I3 | USP47 | EPN1 | 0.5367459 |
| Q96K76 | P51991 | USP47 | HNRNPA3 | 0.535311141 |
| Q96K76 | Q96PU4 | USP47 | UHRF2 | 0.535117026 |
| Q96K76 | Q99549 | USP47 | MPHOSPH8 | 0.534574956 |
| Q96K76 | P49761 | USP47 | CLK3 | 0.531323554 |
| Q96K76 | P42285 | USP47 | MTREX | 0.531275118 |
| Q96K76 | Q9NRR5 | USP47 | UBQLN4 | 0.530578139 |
| Q96K76 | Q9BZF9 | USP47 | UACA | 0.529361805 |
| Q96K76 | Q9BX70 | USP47 | BTBD2 | 0.529291686 |
| Q96K76 | Q13608 | USP47 | PEX6 | 0.528766888 |
| Q96K76 | O15145 | USP47 | ARPC3 | 0.528444828 |
| Q96K76 | Q5T200 | USP47 | ZC3H13 | 0.528376368 |
| Q96K76 | Q96FJ0 | USP47 | STAMBPL1 | 0.528049094 |
| Q96K76 | Q9BVQ7 | USP47 | SPATA5L1 | 0.527159934 |
| Q96K76 | Q008S8 | USP47 | ECT2L | 0.523063186 |
| Q96K76 | O15372 | USP47 | EIF3H | 0.521457144 |
| Q96K76 | Q01804 | USP47 | OTUD4 | 0.520993308 |
| Q96K76 | Q9Y2K7 | USP47 | KDM2A | 0.519458375 |
| Q96K76 | Q1L5Z9 | USP47 | LONRF2 | 0.519350318 |
| Q96K76 | A0AVT1 | USP47 | UBA6 | 0.518648204 |
| Q96K76 | Q14596 | USP47 | NBR1 | 0.514817722 |
| Q96K76 | Q8IYE0 | USP47 | CCDC146 | 0.512496526 |
| Q96K76 | Q15646 | USP47 | OASL | 0.512074057 |
| Q96K76 | O75152 | USP47 | ZC3H11A | 0.509704279 |
| Q96K76 | Q9UBP0 | USP47 | SPAST | 0.508648061 |
| Q96K76 | O14980 | USP47 | XPO1 | 0.507722071 |
| Q96K76 | Q6IQ23 | USP47 | PLEKHA7 | 0.506154457 |
| Q96K76 | P54727 | USP47 | RAD23B | 0.505879738 |
| Q96K76 | Q96T88 | USP47 | UHRF1 | 0.504979894 |
| Q96K76 | Q9UKB1 | USP47 | FBXW11 | 0.504185463 |
| Q96K76 | Q15843 | USP47 | NEDD8 | 0.504024661 |
| Q96K76 | O94776 | USP47 | MTA2 | 0.503864101 |
| Q96K76 | Q9Y3R5 | USP47 | DOP1B | 0.502473911 |
| Q96K76 | Q5JWR5 | USP47 | DOP1A | 0.502473911 |
| Q96K76 | Q15738 | USP47 | NSDHL | 0.502375029 |
| Q96K76 | Q9UGI0 | USP47 | ZRANB1 | 0.501506864 |

| | | | | |
|---|---|---|---|---|
| Q96K76 | Q8IV36 | USP47 | HID1 | 0.501025204 |
| Q96K76 | Q8WWQ0 | USP47 | PHIP | 0.500772417 |
| Q96K76 | P20591 | USP47 | MX1 | 0.5002461 |

**Table 1S: Primer sequences used for qRT-PCR analysis (*Mus musculus*)**

| Gene | Forward Primer | Reverse Primer |
|---|---|---|
| Brcc3 | GGCGGTTCATCTTGAGTCTG | GCGCATTTCCGTTCCAGTG |
| Usp4 | GACCTGAACCGCGTAAAG | AGCACATTCTGGGCAAACC |
| Usp46 | GCTTCTGGCTGTTGTTTGA | TGAATGAGTCTCGCACTCC |
| Usp47 | ACCCTGAATGTTTGGCCTCT | ATGTCCCGTCTTCTGCAGTC |
| Usp40 | GAGAACCAGAAGCGGCTAG | TGGAGAACAAATCGCTGCTC |
| β-actin | GGCTGTATTCCCCTCCATCG | CCAGTTGGTAACAATGCCATGT |
